# An apical membrane complex controls rhoptry exocytosis and invasion in *Toxoplasma*

**DOI:** 10.1101/2022.02.25.481937

**Authors:** Daniela Sparvoli, Jason Delabre, Diana Marcela Penarete-Vargas, Shrawan Kumar Mageswaran, Lev M. Tsypin, Justine Heckendorn, Liam Theveny, Marjorie Maynadier, Marta Mendonça Cova, Laurence Berry-Sterkers, Amandine Guérin, Jean-François Dubremetz, Serge Urbach, Boris Striepen, Aaron P. Turkewitz, Yi-Wei Chang, Maryse Lebrun

## Abstract

Apicomplexan parasites possess secretory organelles called rhoptries that undergo regulated exocytosis upon contact with the host. This process is essential for the parasitic lifestyle of these pathogens and relies on an exocytic machinery sharing structural features and molecular components with free-living ciliates. Here, we performed a *Tetrahymena*-based transcriptomic screen to uncover novel exocytic factors in Ciliata and Apicomplexa. We identified membrane-bound proteins, named CRMPs, forming part of a large complex essential for rhoptry secretion and invasion in *Toxoplasma*. In contrast to previously described rhoptry exocytic factors, TgCRMPs are not required for the assembly of the rhoptry secretion machinery and only transiently associated with the exocytic site - prior to invasion. CRMPs and their partners contain putative host cell-binding domains, and CRMPa shares similarity to GPCR proteins. We propose that the CRMP complex acts as host-molecular sensor to ensure that rhoptry exocytosis occurs when the parasite contacts the host cell.

## INTRODUCTION

Apicomplexan parasites can cause life-threatening diseases including malaria, cryptosporidiosis, and toxoplasmosis. They are obligate intracellular organisms that invade and subvert functions of diverse host cells by releasing multiple adhesins, perforins and effectors from three different secretory organelles: micronemes, rhoptries and dense granules (Lebrun, et al., 2020). The content of rhoptries is secreted directly into the host cell (Besteiro et al., 2009; Gilbert et al., 2007), typically at the onset of host cell contact (Carruthers and Sibley, 1997; Riglar et al., 2011). The signaling pathways that mediate rhoptry discharge are unknown, but they might depend on the initial secretion of microneme proteins (Kessler et al., 2008; Singh et al., 2010). Upon injection into the host cell, rhoptry proteins facilitate invasion by establishing a structure called the moving junction (MJ), which anchors the parasite invasion machinery into the host cell cortex (Besteiro et al., 2011; Guerin et al., 2017). Rhoptry proteins also contribute to the formation of the parasitophorous vacuole (Ghosh et al., 2017) and play key roles in subverting host immune responses (Hakimi et al., 2017; Kemp et al., 2012). How rhoptry content is delivered into the host cell cytoplasm has been a vexing question for decades. Delivery requires docking and fusion of the organelle with the parasite plasma membrane (PPM); this process of exocytosis is coupled with the translocation of rhoptry content across the host plasma membrane (HPM). The latter likely involves the formation of a pore at the junction between the PPM and HPM (Burrell et al., 2021; Dubremetz, 1998; Hanssen et al., 2013; Nichols et al., 1983; Suss-Toby et al., 1996), but its nature and composition are unknown. Excitingly, recent studies revealed new insights into structure and molecular players essential for the exocytic step (Aquilini et al., 2021; Mageswaran et al., 2021; Suarez et al., 2019). Rhoptry exocytosis relies on the proper assembly of a “rosette” of eight particles embedded in the PPM at the apex of the parasite (Aquilini et al., 2021). A similar rosette is present at the exocytic site of ciliate secretory organelles known as trichocysts in *Paramecium tetraurelia* and mucocysts in *Tetrahymena thermophila* (Plattner et al., 1973; Satir et al., 1972) and its presence is a firm requirement for the release of organelle content (Beisson et al., 1976). Cryo-electron tomography (Cryo-ET) of the apical tips of *Toxoplasma* and *Cryptosporidium* zoites revealed the rosette to be part of an elaborate machinery named Rhoptry Secretory Apparatus (RSA). This complex molecular machine connects the rhoptry to the PPM via an intermediate apical vesicle (AV) (Aquilini et al., 2021; Mageswaran et al., 2021). A group of Alveolata-restricted “<u>n</u>on-<u>d</u>ischarge” proteins (Nd6, Nd9, NdP1, NdP2) is required for the formation of the rosette in both Ciliata and Apicomplexa (Aquilini et al., 2021; Froissard et al., 2001; Gogendeau et al., 2005), demonstrating a conserved mechanism for exocytic fusion in Alveolata. However, several aspects of rhoptry secretion remain unknown, including the exact function of Nd proteins in this process, and how rhoptry discharge is regulated and triggered by host cell contact to inject content inside the host.

Here we extend the use of ciliate models, specifically *Tetrahymena thermophila*, to further uncover the mechanism of rhoptry secretion. *Tetrahymena* possesses hundreds of mucocysts concentrated at the plasma membrane which are capable of rapid and synchronous release upon stimulation (Satir, 1977). Following mucocyst exocytosis, the organelles are regenerated *de novo* and docked at the plasma membrane in a highly synchronous process (Haddad and Turkewitz, 1997). These organelles are dispensable for cell survival in laboratory conditions, allowing the mechanisms leading to their formation and release to be analyzed by disruption of genes essential for this pathway. Genes involved in the mucocyst pathway are tightly co-regulated, and new biogenesis-related factors have been identified by the analysis of their expression profiles (Briguglio et al., 2013; Kumar et al., 2014). To further exploit this phenomenon, we used the Coregulation Data Harvester (CDH) tool (Tsypin and Turkewitz, 2017) to automate the search of genes co-regulated with *Tetrahymena Nd* genes and also conserved in Apicomplexa. By this approach, we identified two novel *Tetrahymena* proteins with a role in mucocyst exocytosis. Both proteins show similarities with the Cysteine Repeat Modular Proteins (CRMPs) previously described in *Plasmodium* (Douradinha et al., 2011; Thompson et al., 2007) and two uncharacterized proteins in *Toxoplasma*, named hereafter TgCRMPa and TgCRMPb. We investigated the two uncharacterized *Toxoplasma* homologs and found that, similarly to their ciliate counterparts, they are necessary for rhoptry exocytosis and subsequent parasite invasion. TgCRMPa and TgCRMPb are part of a complex comprising at least two additional yet uncharcaterized proteins, and we demonstrated that one of them is also involved in rhoptry secretion. Unlike the exocytic Nd complex, we found that TgCRMPs are not essential for the assembly of the RSA or the anchoring of the AV to the RSA, and they only accumulate at the exocytic site just prior to invasion and subsequently disappear at the onset of host invasion. Sequence analyses of TgCRMPs showed that they are multipass-transmembrane proteins, contain putative host cell binding domains and they are related to G protein-coupled receptor (GPCR). These features, together with their transient localization to exocytic sites, support a role for this complex in the signaling pathway that coordinates rhoptry content discharge with host contact.

## RESULTS

### A *Tetrahymena*-based strategy to search for new exocytic factors conserved in Apicomplexa

We recently demonstrated that a group of Alveolata-restricted proteins, Nd6, Nd9, NdP1 and NdP2, regulate mucocyst/trichocyst and rhoptry exocytosis in ciliates and apicomplexans, respectively (Aquilini et al., 2021). In addition, we found that *Toxoplasma* protein ferlin 2 (TgFer2), which has a role in rhoptry secretion (Coleman et al., 2018), is associated with the Nd complex. To test a conserved role of Fer2 in the two systems, we searched for the *Tetrahymena* homolog of TgFer2 and verified its role in exocytosis. Phylogenetic analysis of the four *Tetrahymena ferlin* genes revealed TTHERM_00886960 as the closest homolog of TgFer2 (Figure S1A). We deleted the expressed (macronuclear) copies of *TtFer2* (Figures S1B, S1C) and found that the *TtΔfer2* mutant cells have a defect in mucocyst release when stimulated with the secretagogue dibucaine (Figure S1D). The organelles appeared properly formed and docked at the plasma membrane (Figure S1E). Also arguing against any defect in biogenesis was our finding that the content protein Grl1 was proteolytically processed (Figure S1F), an essential step in mucocyst maturation (Chilcoat et al., 1996). These results demonstrate a role for TtFer2 in exocytosis, further supporting the conservation of exocytic mechanisms in Alveolata.

Genes involved in mucocyst exocytosis are tightly co-regulated, as shown by the transcriptional profiles of *Tetrahymena Nd6*, *Nd9*, *NdP1*, *NdP2* and *Fer2* genes in different life stages (Figure 1A), and we hypothesized that yet uncharacterized secretion factors might be similarly co-regulated, and therefore identifiable by their distinct expression profiles. We took advantage of a bioinformatic tool specifically developed for *Tetrahymena*, called the Coregulation Data Harvester (CDH) (Tsypin and Turkewitz, 2017), to automate the search for co-regulated genes. To identify exocytosis-related genes with orthologs in Apicomplexa, and as such potentially involved in rhoptry secretion, we set up the CDH search to look for genes conserved in *T. gondii* and *P. falciparum* (Figure 1B). We performed the CDH analysis using *Tetrahymena* Nd6, NdP1, NdP2 and Fer2 as separate queries, but excluded TtNd9 due to its very low expression level. For each query we identified those *Tetrahymena* genes with homologs in *T. gondii* and *P. falciparum*, and prioritized a list of candidates shared by at least three of the four queries (Figure 1B; Table S1).

**Figure 1.**
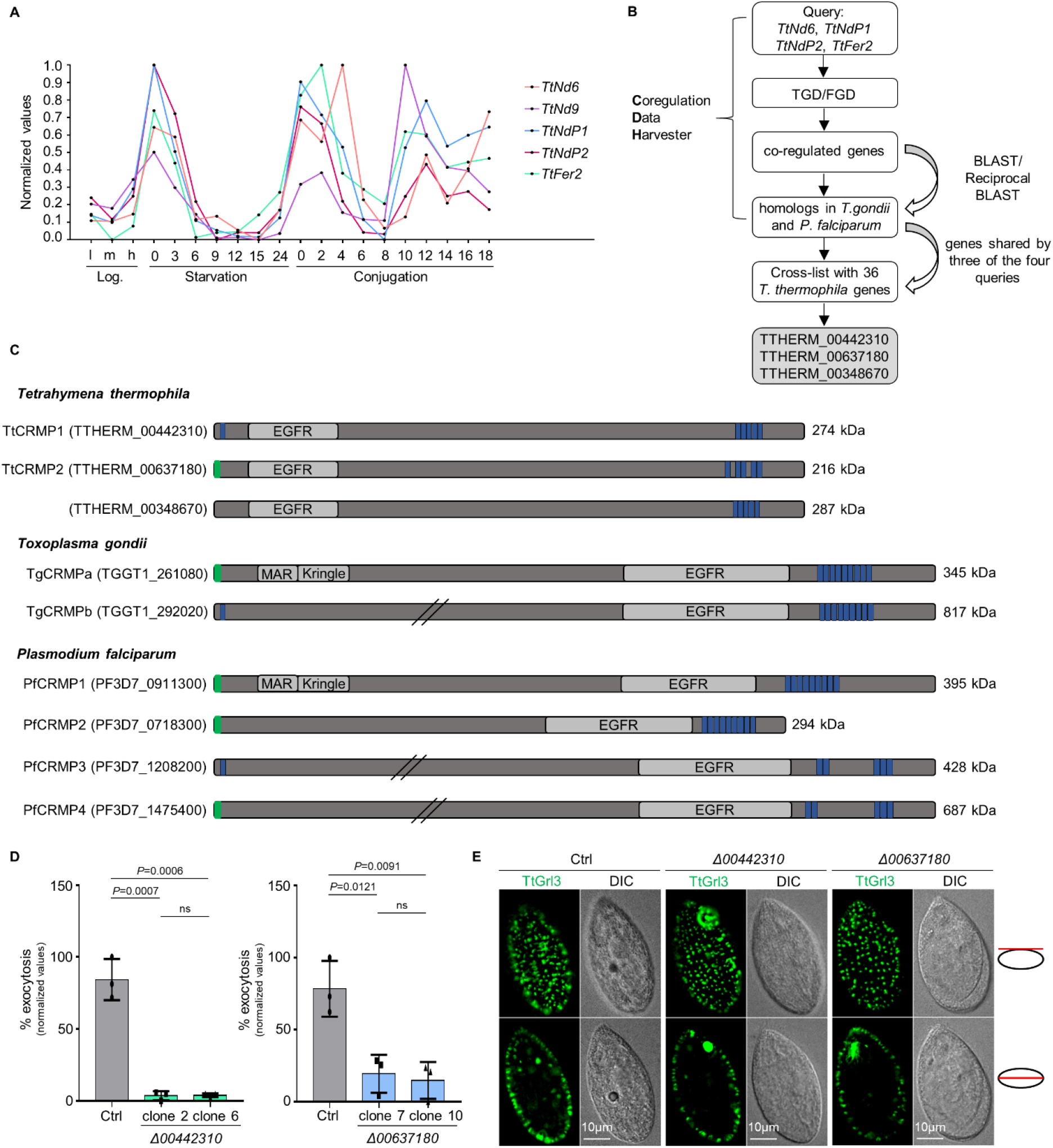
A *Tetrahymena*-based strategy identified two new non-discharge genes conserved in *Toxoplasma gondii* and *Plasmodium falciparum*. (A) Expression profiles of *Tetrahymena Nd* genes involved in mucocysts exocytosis. The plot values were downloaded from http://tfgd.ihb.ac.cn and normalized to that of the gene’s maximum expression level. The data were collected from growing (low, <u>m</u>edium and <u>h</u>igh culture density) and starved (S0– S24) cultures, and different time points during conjugation (C0–C18). (B) *Tetrahymena*-based bioinformatics approach for identifying new exocytic factors. (C) Protein domains in *T. thermophila*, *T. gondii* and *P. falciparum* CRMPs. EGFR (Epidermal Growth Factor Receptor), MAR (Microneme Adhesive Repeat) and Kringle domains are shown in grey. Green: predicted signal peptide; blue: transmembrane domains; slanted lines: truncation of the full length protein sequence. (D) Quantification of the exocytic response of *Tetrahymena Δ00442310* and *Δ00637180* mutants to dibucaine stimulation. Mean ± SD (n = 3 biological replicates, each with 2 technical replicates). *P* values were measured by two-tailed *t*-test. (E) Immunofluorescence images of *Tetrahymena* cells. Mucocysts in wildtype (Ctrl), *Δ00442310* and *Δ00637180* cells were immunostained with mAbs 5E9 which recognizes the granule protein Grl3. The mucocyst pattern in the mutants was similar to wildtype. Single focal planes of surface and cross sections are shown for each cell. DIC: differential interference contrast.

Among the 37 candidates identified, three (TTHERM_00442310, TTHERM_00637180, TTHERM_00348670) encode proteins containing similar features including an Epidermal-Growth-Factor-receptor domain and multiple C-terminal transmembrane domains. These domains are shared by the putative homologs found in *T. gondii* and *P. falciparum* (Figure 1C). The *Plasmodium* homologs were previously described as members of a family of four genes named CRMPs for Cysteine-Repeat Modular Proteins (Douradinha et al., 2011; Thompson et al., 2007), but the two *Toxoplasma* counterparts, which we called TgCRMPa (TGGT1_261080) and TgCRMPb (TGGT1_292020) had not been previously studied. In addition to the common features, TgCRMPa and PfCRMP1 possess a Kringle domain known to bind proteins (Patthy et al., 1984). Secondary structure-based predictions (Zimmermann et al., 2018) revealed that TgCRMPa and PfCRMP1 also possess a MAR (Microneme Adhesive Repeat) domain at the N-terminus, which is a novel carbohydrate-binding domain found in microneme proteins of enteroparasitic coccidians and known to interact with sialic acids (Blumenschein et al., 2007; Friedrich et al., 2010). Interestingly, TgCRMPa and PfCRMP1 are also predicted to be G protein-coupled receptor (GPCR)-like proteins by PANTHER analysis (Mi et al., 2021). These similarities between TgCRMPa and PfCRMP1 are consistent with their evolutionary relatedness (Figure S2A).

To validate the *in-silico* screen, we first knocked-out the three *Tetrahymena* genes (Figure S1B). We obtained complete knockout lines for the genes TTHERM_00442310 and TTHERM_00637180 (Figure S1G) but not for TTHERM_00348670 (not shown). *Δ00442310* and *Δ00637180* cells were impaired in mucocyst discharge (Figure 1D) but not in biogenesis, as judged by normal mucocyst staining (Figure 1E) and correct processing of the Grl1 precursor (Figure S1H). We concluded that the affected step was exocytosis. These data showed that TTHERM_00442310 and TTHERM_00637180 are two novel non-discharge proteins, and prompted us to study the function of their apicomplexan homologs.

### TgCRMPa and TgCRMPb are essential for rhoptry secretion and host cell invasion

PfCRMP1 and PfCRMP2 are not essential for the asexual stage of *P. berghei*, but they appear to control sporozoite invasion of the mosquito salivary glands (Douradinha et al., 2011; Thompson et al., 2007). We tested the function of CRMP proteins in the apicomplexan model *T. gondii*. TgCRMPa and TgCRMPb are predicted to be fitness-conferring genes in tachyzoites (Sidik et al., 2016); thus, we generated inducible knockdown lines (iKD). We introduced a triple HA tag at the C-terminus of TgCRMPa and TgCRMPb (Figures S2B, S2C) and then replaced the endogenous promoter of each gene with the anhydrotetracycline (ATc)-regulatable TetOSag4 promoter (Figures S2D, S2E) to switch off gene expression by using ATc (Meissner et al., 2002). Two bands were detected by western blot for both TgCRMPa-HA_3_ and TgCRMPb-HA_3_ and appeared less abundant in the ATc-untreated (0h) iKD lines compared to the solely HA_3_-tagged lines (Figure 2A), indicating that the promoter switch reduced transcription of both *TgCRMPs* genes. The two bands might reflect proteolytic processing, and both disappeared in the iKD lines upon ATc treatment (24-48h) (Figure 2A). Expression and efficient depletion of the tagged proteins were also confirmed by immunofluorescence microscopy (Figures 2B, S2F). We observed diffuse vesicle-like staining of TgCRMPa-HA_3_ and TgCRMPb-HA_3_ throughout the entire parasite that disappears upon ATc incubation. Occasionally, the staining appeared more concentrated at the apex of the tachyzoite, similar to micronemes visualized using antibodies to AMA1 (Figure 2B). The apical concentration of TgCRMPs was more evident in the untreated iKD lines (-ATc; Figures 2B, 2C, S2F), likely due to lower levels of the proteins, as shown in figure 2A. However, they did not extensively co-localize with the microneme proteins AMA1, MIC2, GAMA and PLP1 by confocal microscopy, as shown for TgCRMPb-HA_3__iKD (Figure 2D, S2G, S2H).

**Figure 2.**
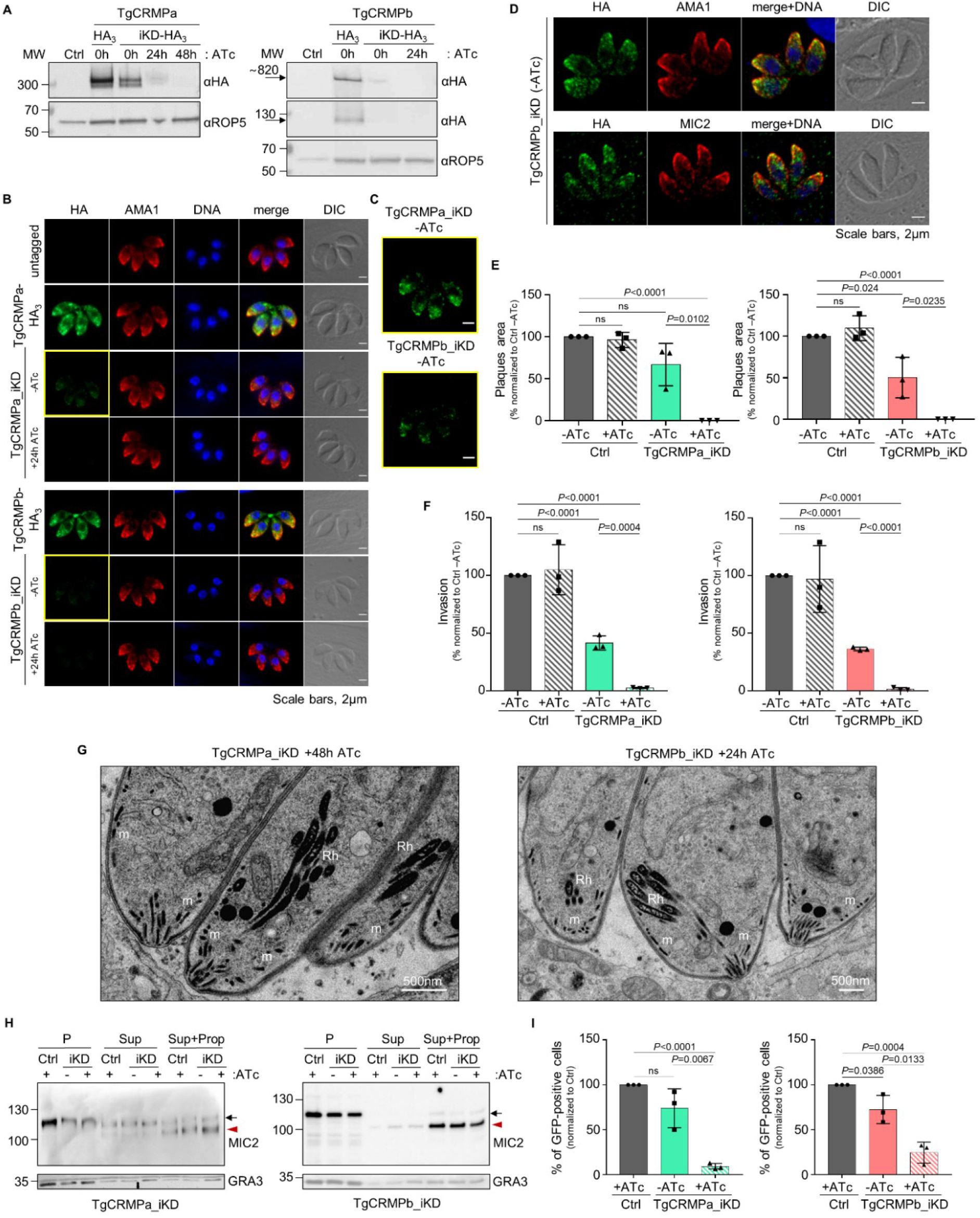
TgCRMPa and TgCRMPb are essential for rhoptry secretion and host cell invasion in *Toxoplasma*. **(A)** Immunoblot of lysates from parental (Ctrl) and tagged lines (TgCRMPa-HA_3_ and TgCRMPb-HA_3_) together with inducible-knockdown lines (TgCRMPa-HA_3__iKD and TgCRMPb-HA_3__iKD) treated with ATc for 0, 24 or 48h. Blots were probed with anti-HA Abs to visualize tagged CRMPa and CRMPb. TgROP5 was used as loading control and visualized by anti-ROP5 Abs. Two close bands around 300kDa were detected for TgCRMPa. A ∼820kDa protein, corresponding to the predicted size for TgCRMPb, was observed together with a ∼130kDa band. Both bands for CRMPa and CRMPb disappeared upon 48 and 24h ATc incubation, respectively. **(B)** Immunofluorescence microscopy of intracellular parasites (untagged, TgCRMPa-HA_3_, TgCRMPb-HA_3_, and TgCRMPs-depleted (iKD) lines). Parasites were labeled with anti-HA and anti-AMA1 Abs to visualize CRMPs-HA_3_ and micronemes, respectively. The nuclei (DNA) are stained with Hoechst. DIC: differential interference contrast. TgCRMPs-HA_3_ shows a heterogeneous distribution within the parasite cytosol, occasionally showing a microneme-like apical gradient (yellow boxes), highlighted in **(C)** (contrast and brightness increased). CRMPa and CRMPb fluorescence significantly disappeared upon 48 and 24h ATc treatment, respectively. Shown are single focal planes. **(D)** Confocal immunofluorescence images of TgCRMPb-depleted (iKD) intracellular tachyzoites. Parasites were stained with anti-HA and with anti-AMA1 and anti-MIC2 Abs to visualize TgCRMPb and micronemes, respectively. The nuclei (DNA) are stained with Hoechst. Shown are single focal planes. **(E)** Quantification of plaques areas for control, TgCRMPa_iKD and TgCRMPb_iKD in absence of ATc, and upon 24 and 48h ATc treatment for TgCRMPb and TgCRMPa, respectively. Values are reported as mean ± SD (n = 3 biological replicates, each with 3 technical replicates). **(F)** Invasion of TgCRMPa- and TgCRMPb-depleted tachyzoites upon 48 and 24h treatment with ATc, respectively. Data are reported as mean ± SD (n = 3 biological replicates, each with 3 technical replicates). **(G)** Electron micrographs of TgCRMPa_iKD and TgCRMPb_iKD intravacuolar parasites treated with ATc for 48 and 24h, respectively. Micronemes (m) and rhoptries (Rh) appeared properly localized and shaped in both mutants. **(H)** Quantification of microneme secretion in TgCRMPa- and TgCRMPb-depleted tachyzoites was measured by detecting the processed form (arrowhead) of TgMIC2 (arrow) in the media. Control, TgCRMPa_iKD and TgCRMPb_iKD parasites, ATc-treated (+) and untreated (-), were stimulated with propranolol to release microneme contents. Blots were probed with anti-MIC2 (secretion of micronemes) and anti-GRA3 (constitutive secretion of dense granules). P: Parasites pellet. Sup: Supernatant from untreated parasites. Sup+Prop: Supernatant from parasites treated with propranolol. The results are representative of two independent experiments. **(I)** Quantification of rhoptry secretion in TgCRMPa_iKD and TgCRMPb_iKD parasites upon 48 and 24h ATc-treatment, respectively, using the SeCreEt system (Koshy et al., 2010). Successful secretion of rhoptry proteins into the host causes a switch from red to green fluorescence in a reporter host cell line. CRMPs-depleted parasites were unable to efficiently deliver rhoptry content into the host cytosol. Data are represented as mean ± SD (n = 3 biological replicates). *P* values in E, F, I, were measured by two-tailed *t*-test.

We tested the overall ability of TgCRMPs_iKD lines to proliferate and lyse host cells, and found that treatment with ATc (+ATc) resulted in loss of plaque formation; TgCRMPb_iKD parasites exhibited significant defects in plaque formation even in the absence of ATc (-ATc) (Figure 2E). Importantly, parasites could efficiently replicate, egress from the PV and attach to host cells (Figures S2I, J, K) but were severely impaired in host cell invasion (Figure 2F). Invasion depends on the sequential secretion of microneme and rhoptry proteins. Since the morphology and positioning of both organelles appeared unaltered by ATc treatment (Figures 2B, 2G, S2F), we tested whether their discharge was disrupted. While microneme secretion occurred normally in TgCRMPs-depleted parasites (Figure 2H), the discharge of rhoptry contents into the host cell was greatly impaired (Figure 2I). We conclude that CRMP proteins serve a crucial function, conserved between Ciliata and Apicomplexa, in the regulated discharge of secretory organelles. We named *Tetrahymena* TTHERM_00442310 and TTHERM_00637180, TtCRMP1 and TtCRMP2, respectively.

### TgCRMPa and TgCRMPb form a complex with two additional membrane proteins

TgCRMPa and TgCRMPb have similar organization and function, suggesting that they might collaborate in regulating rhoptry secretion. To test this, we isolated each TgCRMP-HA_3_ and its associated proteins by affinity capture (Figure S3A) and analyzed the associated proteins by liquid chromatography-tandem mass spectrometry (Tables S2 and S3). Indeed, TgCRMPa and TgCRMPb were associated with each other (Figure 3A), a result also confirmed by co-immunoprecipitation experiments with parasites co-expressing TgCRMPa-FLAG_3_ and TgCRMPb-HA_3_ (Figures 3B, S3B-D). Moreover, TgCRMPs robustly associate with two additional uncharacterized membrane proteins (Figure 3A, C), Tg247195 and Tg277910. Tg277910 and Tg247195 possess one and three thrombospondin type 1 (TSP-1) domains, respectively (Figure 3C), known to participate in cell adhesion (Adams and Tucker, 2000). In addition, Tg247195 possesses a H-type lectin domain (Pietrzyk-Brzezinska and Bujacz, 2020) and, interestingly has a role in rhoptry secretion (Furio Spano, personal communication). To determine the function of Tg277910, we generated an inducible knockdown HA_3_-tagged line (Tg277910_iKD) (Figure S3E-G). A single Tg277910-HA_3_ band was detected by western blot in the absence of ATc, and the protein completely disappeared after ATc treatment in both western blot (Figure S3H) and IFA (Figures 3D, S3I). We observed a consistent reduction in the area of lytic plaques in ATc-treated tachyzoites (Figure S3J), that was not related to the disruption of parasite replication, egress or attachment (Figure S3K-M) but a consequence of the inability of the parasites to invade the host cell (Figure 3E). This defect was associated with loss of rhoptries discharge (Figure 3F), but not that of micronemes (Figure 3G). We note that again the morphology and localization of these two secretory organelles were not affected by protein depletion (Figures 3D, S3I).

**Figure 3.**
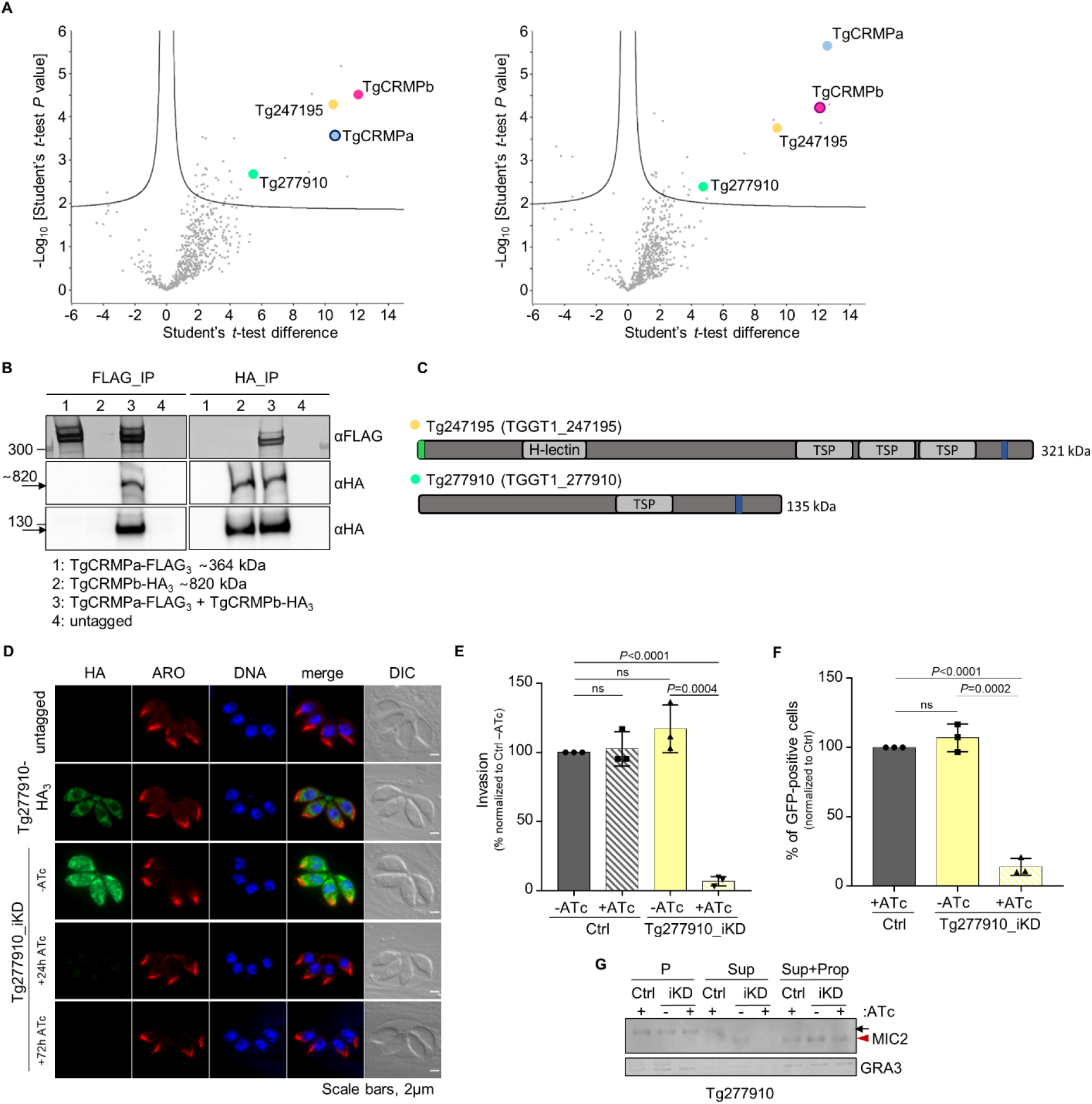
TgCRMPa and TgCRMPb are in complex with two additional membrane proteins, one of which is required for rhoptry exocytosis. **(A)** Mass spectrometric identification of proteins co-isolated with HA_3_-tagged TgCRMPa (left plot) and TgCRMPb (right plot). The volcano plots shown here were generated by plotting the −log_10_ *t*-test *P*-value versus the *t*-test difference. The colored dots mark members of the CRMP complex. Baits are indicated by darker outlines. **(B)** Co-immunoprecipitation of TgCRMPa and TgCRMPb. Lysates from parasites co-expressing TgCRMPa-FLAG_3_ and TgCRMPb-HA_3_ were split and incubated with either anti-FLAG (left panels) or anti-HA beads (right panels). Eluates were subjected to SDS-PAGE and immunoblotted with anti-HA and anti-FLAG Abs. Untagged, TgCRMPa-FLAG_3_ and TgCRMPb-HA_3_ parasites were used as controls. TgCRMPs robustly associate with each other. **(C)** Protein domains of Tg247195 and Tg277910, co-purified with TgCRMPs. TSP: thrombospondin domain; H-lectin: lectin binding domain. Green: predicted signal peptides; blue: transmembrane domains. **(D)** Immunofluorescence images of untagged, Tg277910-HA_3_ and Tg277910_iKD intracellular tachyzoites. Parasites were stained with anti-HA and anti-ARO Abs to label Tg277910 and rhoptries, respectively. The nuclei (DNA) are stained with Hoechst. Tg277910 localization is similar to that of TgCRMPs, with an increased fluorescence in the iKD line (-ATc). Tg277910 signal almost completely disappeared upon 72h ATc-treatment. Single focal planes are shown. **(E)** Invasion of parental and Tg277910_iKD lines in absence of ATc, and upon 72h ATc treatment. Values are reported as mean ± SD (n=3 biological replicates, each with 3 technical replicates). **(F)** Quantification of rhoptry secretion in Tg277910_iKD by SeCreEt system as described in figure 2H. Tg277910-depleted parasites failed to deliver rhoptry content into the host cytosol. Data are shown as mean ± SD (n=3 biological replicates). *P* values in E, F, were measured by two-tailed *t*-test. **(G)** Quantification of microneme secretion in control and Tg277910-depleted tachyzoites was measured as in figure 2G. Blots were probed with anti-MIC2 (secretion of micronemes) and anti-GRA3 (constitutive secretion of dense granules). P: Parasites pellet. Sup: Supernatant from untreated parasites. Sup+Prop: Supernatant from parasites treated with propranolol. The results are representative of two independent experiments.

We did not find any of the rhoptry exocytic factors described previously (TgNd6, TgNd9, TgNdP1, TgNdP2 and TgFer2) among the proteins co-isolated with TgCRMPs, suggesting that CRMPs are part of a distinct complex regulating rhoptry secretion, a result also supported by the mass spectrometry analysis of Nd9 and NdP1 pulldowns (Aquilini et al., 2021).

### *Toxoplasma* and *Tetrahymena* CRMP proteins are not required for rosette formation, RSA assembly or AV positioning in *T. gondii*

Our findings on *Toxoplasma* and *Tetrahymena* CRMPs strongly suggest that they have a role in exocytosis, the last step of the secretory pathway, which depends on the proper assembly of the rosette at the plasma membrane (Aquilini et al., 2021; Plattner et al., 1973). Since CRMPs are predicted to be transmembrane proteins (Figure 1C), we considered that they might be rosette components. To test this hypothesis, we performed thin-section and freeze fracture electron microscopy (EM) analyses of CRMP mutants. *Tetrahymena* mutant *ΔCRMP1* accumulated well-formed rosettes at the plasma membrane as shown by freeze fracture EM of the cell surface (Figure 4A, B), arrayed in the known pattern of mucocyst docking sites (Figure 4C). In *Toxoplasma*, no apparent defects were observed in the positioning of the AV in CRMPs_iKD strains after ATc treatment (Figure 4D). Nor was there apparent defect in the assembly of the rosette, as shown for TgCRMPa-depleted tachyzoites (Figure 4E). To inspect possible minor defects affecting the RSA, we performed cryo-electron tomography (cryo-ET) on frozen-hydrated TgCRMPb-depleted cells. The subtomogram average of the RSA showed an 8-fold symmetry of defined densities holding the AV as seen previously in the wild-type (Mageswaran et al., 2021)(Figure 4F). We did not observe profound rearrangements of the RSA densities and their distance to the AV in the TgCRMPb-depleted parasites compared to wild-type (Figure 4G, H), in stark contrast to what we previously showed after TgNd9 depletion (Mageswaran et al., 2021). We only observed a minor alteration in the AV shape and anchoring angle (Figure 4G, I-K). In conclusion, since freeze-fracture EM and cryo-ET demonstrate that CRMPs are not essential for building the rhoptry secretion machinery, CRMPs have a function different from that of the previously described Nd complex.

**Figure 4.**
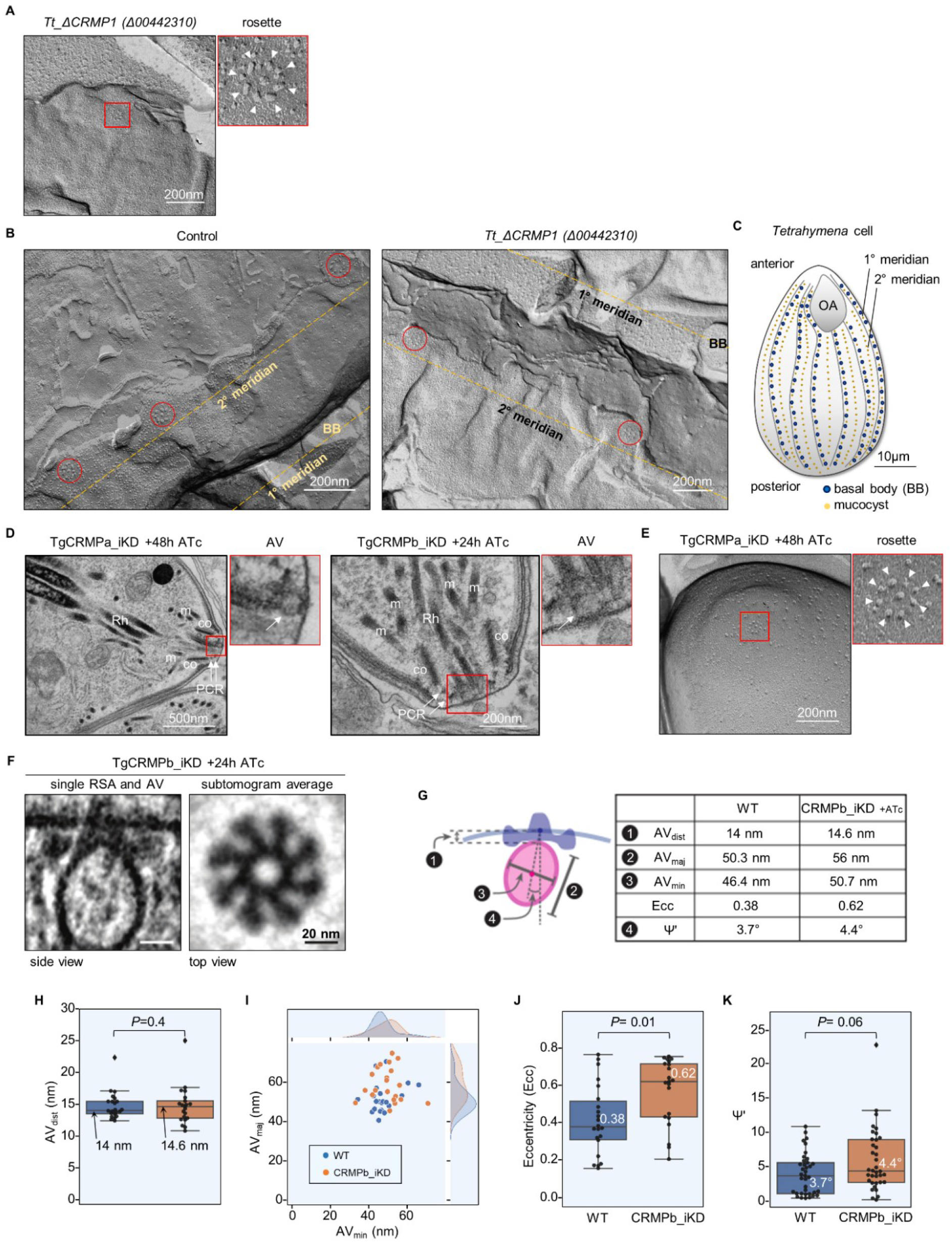
TgCRMPa and TgCRMPb are dispensable for apical vesicle positioning and, similarly to TtCRMP1, show a well-assembled rosette. **(A)** Freeze-fracture electron micrograph of *Tetrahymena ΔCRMP1* cell surface showing a representative rosette (red box) at the plasma membrane. On the right, the magnified rosette with the eight intramembranous particles (IMPs, arrowheads) surrounding the central one. **(B)** Freeze-fracture electron micrographs of larger cell surfaces for *TtΔCRMP1* and wildtype in which multiple rosettes, corresponding to multiple mucocyst docking sites, are correctly aligned along 2° meridians in both lines. **(C)** Cartoon of a *Tetrahymena* cell (surface section). Mucocysts primarily occupy sites along 2° meridians which mark the membrane spaced between two consecutive 1° meridians, defined in turn by longitudinal rows of basal bodies (BB). Mucocysts are also found at lower frequency between BBs. 1° and 2° meridians regularly span the length of the cell, from the anterior oral apparatus (OA) to the cell posterior. **(D)** Electron micrographs of TgCRMPa_iKD and TgCRMPb_iKD intracellular tachyzoites treated with ATc for 48 and 24h, respectively. A well-formed apical vesicle (AV) appears correctly positioned at the parasite apex in both mutants (magnified red box, arrow). Rh rhoptry; m microneme; co conoid; PCR pre-conoidal rings, indicated by arrows. **(E)** Freeze-fracture electron micrographs of the apex of a TgCRMPa_iKD extracellular tachyzoite treated with ATc for 48h. A well-assembled rosette (red box) was observed at the center of the parasite apex, and magnified on the right. **(F)** A tomogram slice showing the apical vesicle (AV)-anchoring on the plasma membrane in TgCRMPb-iKD line; anchoring is mediated by a rhoptry secretory apparatus (RSA) that is morphologically undistinguishable from the wildtype (WT). Left: side view presented by a central slice through the AV and the RSA. Right: top view presented by a slice through the subtomogram average, revealing the RSA densities that anchor the AV. **(G-K)** AV dimensions and anchoring parameters in WT and TgCRMPb-iKD. **(G)** Left: a schematic depicting the parameters. Right: a table summarizing their measurements. **(H)** AV anchoring distance (the shortest distance measured from the parasite apex to the AV membrane). **(I)** AV dimensions (AV_maj_ and AV_min_). **(J)** AV eccentricity (Ecc) calculated using the AV dimensions (AV_maj_ and AV_min_) in panel I. **(K)** AV orientation parameter (Ψ’) that measures the AV anchoring angle. Panels H, J, and K show a combination of boxplot and swarmplot for each dataset. For the boxplot, the lower and upper boundaries of the box represent the 1^st^ and 3^rd^ quartiles (Q1 and Q3), whiskers extend to 1.5 times the interquartile range (Q3-Q1) below and above Q1 and Q3, and points outside (diamonds) are regarded as outliers. The horizontal divider within the box represents the median with the value noted next to it. For the swarmplot, each data point represents a measurement from a tomogram. Panel I shows a jointplot, which is a combination of a bivariate scatterplot and two marginal univariate kernel density estimate plots (a.k.a. probability density plots), one each for AV_maj_ and AV_min_. Mann–Whitney U tests were used to calculate the *P*-values.

### TgCRMPa and TgCRMPb accumulate at the tip of the extruded conoid in extracellular tachyzoites

Since the CRMPs labeling was reminiscent of MICs, which are typically released on the surface of parasite upon egress, we analyzed the location of CRMPs on extracellular parasites. TgCRMPa-HA_3_ and TgCRMPb-HA_3_ were found to consistently accumulate at the tip of the extruded conoid in freshly egressed parasites kept in contact with host cells (Figure 5A, left panels) and in those treated with the calcium ionophore A23187 (Figure 5A, right panels; 5B), which artificially induces conoid extrusion (Mondragon and Frixione, 1996) and microneme secretion (Carruthers and Sibley, 1999). This staining appears as a tiny dot at the apex of the parasite and thus contrasts with the redistribution of MICs proteins at the surface of the parasite (Carruthers and Sibley, 1999). This accumulation did not occur upon TgCRMP depletion (Figure S4A) and was undetectable without permeabilization (not shown), indicating that it was not a staining artifact and that the C-terminus of TgCRMPa is likely cytoplasmic with the adhesion domains facing the extracellular space. This topology is also supported by TMHMM (http://www.cbs.dtu.dk/services/TMHMM/) and TOPCONS (http://topcons.cbr.su.se) modelling, and by the degradation of TgCRMPa when C-terminally tagged with the auxin-inducible degron (mAID) system (Figure S4B, C). The mAID tag targets the fusion protein to the proteasome upon addition of 3-indoleacetic acid (IAA or auxin), when it is topologically oriented towards the cytosol (Figure 5C) (Nishimura et al., 2009). After adding IAA to the medium, we observed depletion of TgCRMPa by western blot (Figure 5D) indicating that the C-terminus of TgCRMPa is indeed found in the cytosol. Confirming our previous findings, CRMPa IAA-dependent degradation blocked the mutant’s ability to form plaques in host cell monolayers (Figure S4D).

**Figure 5.**
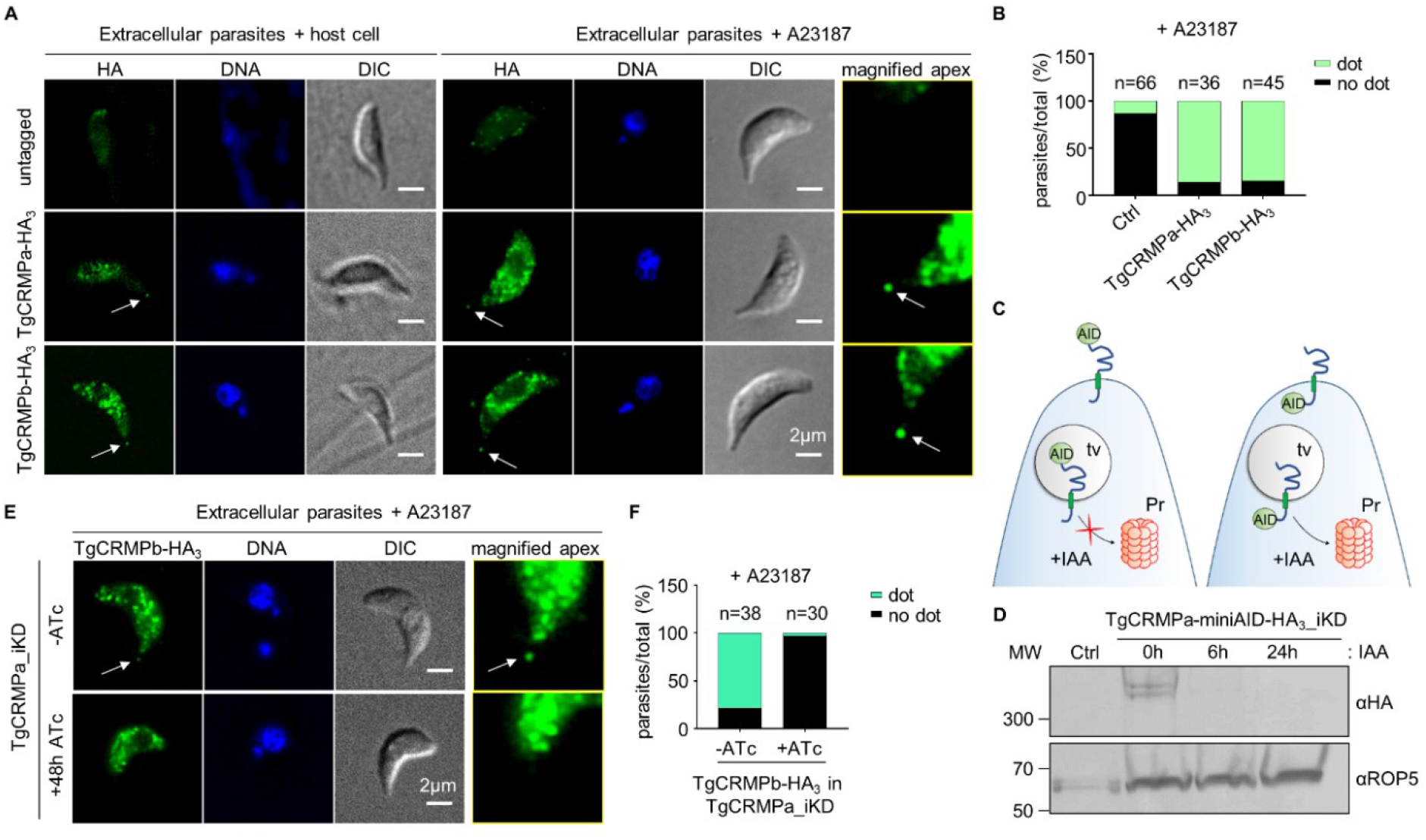
TgCRMPa and TgCRMPb accumulate at the apical tip of extracellular tachyzoites. **(A)** Immunofluorescence images of extracellular tachyzoites of untagged, TgCRMPa-HA_3_ and TgCRMPb-HA_3_ parasites, incubated either with host cell monolayers for 2min, or with ionophore A23187, to induce natural or artificial conoid extrusion, respectively. Parasites were immunostained with anti-HA Abs; DNA was labeled by Hoechst. CRMPa and CRMPb consistently accumulate at the tip of extruded conoids (arrows). The apexes of A23187-treated parasites were magnified on the right, and increased in brightness and contrast to highlight the apical dots. DIC: differential interference contrast. Single focal planes are shown. **(B)** Quantification of the dot pattern upon A23187 treatment shown in A). Values are expressed as percentage of parasites showing (dot) or lacking (no dot) the apical accumulation of TgCRMPa and TgCRMPb; n= number of parasites analyzed per line. **(C)** Cartoon depicting the targeting of a membrane protein to the proteasome (Pr) by the AID-degron system when the AID-fused C-terminus is exposed to the cytosol. IAA: 3-indoleacetic acid or auxin; AID: auxin-inducible degron; tv: transport vesicle. **(D)** Whole cell lysates from TIR1-expressing parental line (Ctrl) and TgCRMPa-miniAID-HA_3__iKD lines were immunoblotted with anti-HA Abs to visualize tagged CRMPa in IAA-treated and untreated samples. CRMPa disappeared upon 24h incubation with IAA, suggesting that the miniAID-HA_3_-tagged C-terminus is exposed towards the cytosol. TgROP5 was used as loading control and detected with ROP5 Abs. MW: molecular weight standards. **(E)** Immunofluorescence images of extracellular TgCRMPb-HA_3_ tachyzoites in presence (+48h ATc) or absence (-ATc) of TgCRMPa-FLAG_3_ (iKD line). Parasites were stained with anti-HA Abs to visualize CRMPb. TgCRMPb localization at the tip of the extruded conoid (arrow) disappears upon TgCRMPa depletion, but it is still detected in the cytoplasm (lower panel). DNA is labeled by Hoechst. Single focal planes are shown. DIC: differential interference contrast. **(F)** Quantification of the dot pattern shown in (E). The values are reported as in (B).

To test whether the apical localization of TgCRMPa and TgCRMPb were inter-dependent, we generated an inducible knockdown (iKD) for TgCRMPa, in which TgCRMPa was tagged with a triple FLAG tag and TgCRMPb with a triple HA tag (Figure S4E, F). Upon depletion of TgCRMPa-FLAG_3_ by ATc treatment (Figure S4G), TgCRMPb-HA_3_ was readily detected in the cytosol but its apical accumulation disappeared (Figure 5E, F), suggesting that the localization of TgCRMPb-HA_3_ at the tip of extracellular parasites is dependent on proper complex assembly.

Altogether, these results suggest that TgCRMPa and TgCRMPb are potentially associated together with the site of rhoptry exocytosis.

### TgCRMPa and TgCRMPb accumulate at the site of exocytosis with TgNd6 but behave differently during invasion

The apical accumulation of TgCRMPa and TgCRMPb in extracellular parasites was reminiscent of that of TgNd6, a protein related to the rhoptry secretion machinery, in intracellular parasites (Aquilini et al., 2021). To assess if CRMPs and Nd6 co-localize at the site of rhoptry exocytosis, we generated *T. gondii* strains co-expressing TgCRMPa-HA3 or TgCRMPb-HA_3_ with TgNd6-TY_2_ (Figure S5A-C). TgCRMPs and TgNd6 appeared to occupy distinct compartments in intracellular parasites, with only Nd6 puncta at the apical ends of tachyzoites (Figure 6A, TgCRMPa; Figure S5D, TgCRMPb), as previously shown (Aquilini et al., 2021). Remarkably, we found TgNd6 overlapping with TgCRMPs at the tip of the extruded conoid in extracellular parasites (Figures 6A, S5D; lower panels), a result confirmed using ultrastructure expansion microscopy (U-ExM) (Figure 6B). Upon parasite expansion, we could measure a ∼40% overlap between TgCRMPs and TgNd6 at the tip of the extruded conoid (Figure 6C), indicating that the two proteins might be spatially very close but part of distinct complexes, in agreement with the mass spectrometry data and the observation that CRMPa and CRMPb persist at the apical tip in the Nd9 mutant defective in RSA assembly (Figures 6D; S5E-G).

**Figure 6.**
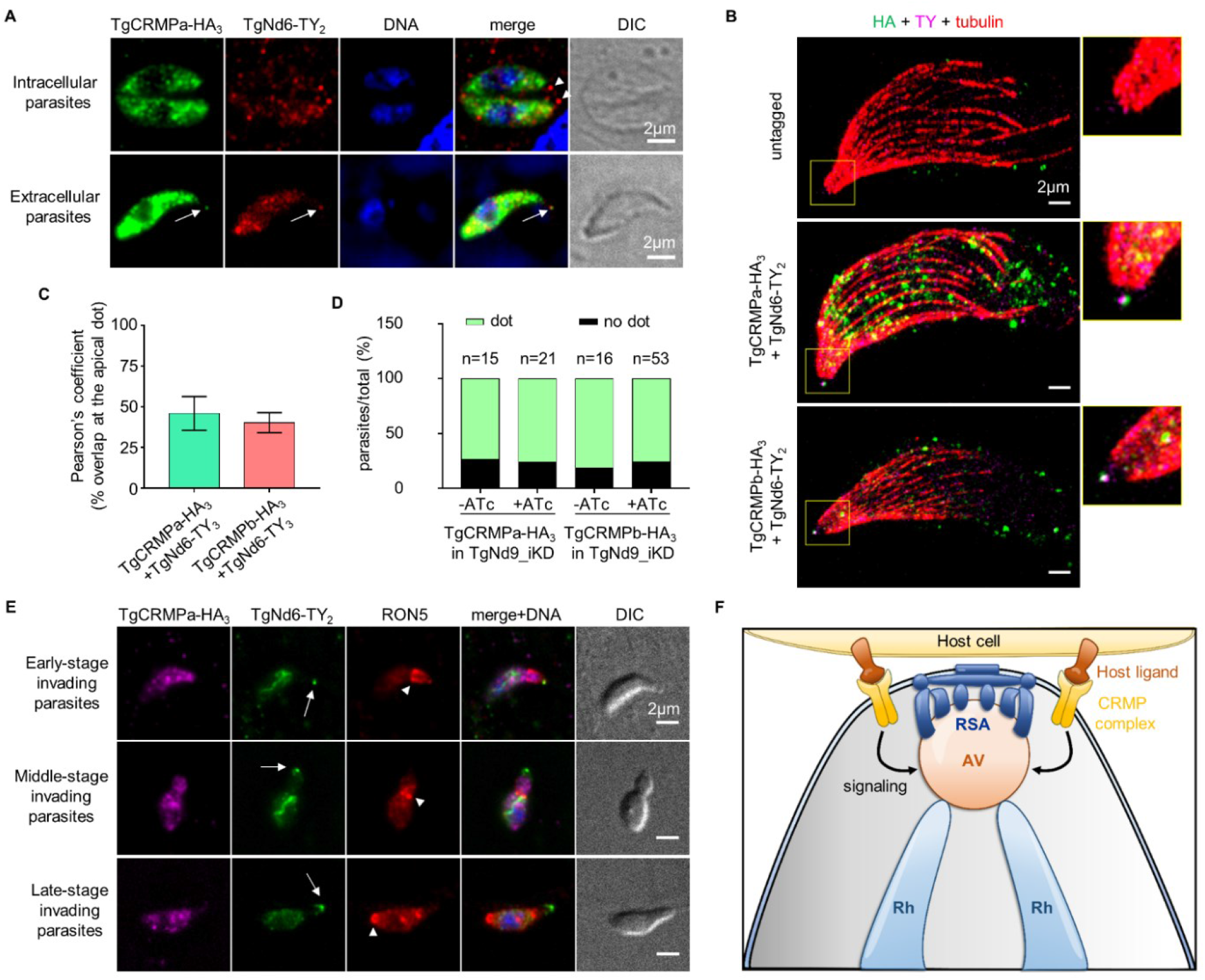
CRMP and Nd complexes differentially regulate rhoptry secretion at the exocytic site in *Toxoplasma gondii*. **(A)** Immunofluorescence images of intracellular (upper panel) and extracellular (lower panel) tachyzoites co-expressing TgCRMPa-HA_3_ and TgNd6-TY_2_. Extracellular parasites were incubated with host cell monolayers for 2min prior to fixation. Parasites were stained with anti-HA and anti-TY Abs to label CRMPa and Nd6, respectively. Nd6, but not CRMPa, accumulates at the tachyzoite apex in intracellular parasites (arrowheads), while both proteins localize at the tip of the extruded conoid in extracellular parasites (arrows). DNA is labeled by Hoechst. Single focal planes are shown. DIC: differential interference contrast. **(B)** Ultrastructure Expansion Microscopy of extracellular tachyzoites, either untagged or co-expressing TgCRMPa-HA_3_ or TgCRMPb-HA_3_ with TgNd6-TY_2_. Parasites were treated with A23187 prior to fixation and preparation for U-ExM, and stained with anti-HA, anti-TY and anti-α/β tubulin Abs to label CRMPs, Nd6 and microtubules, respectively. Shown are maximum intensity projections of z-stack confocal images. CRMPs overlap with Nd6 at the tip of the extruded conoid (yellow selection). A magnified image of the apical tip of each parasite is shown on the right. **(C)** Extent of colocalization between TgCRMPs-HA_3_ and TgNd6-TY_2_ in the apical dot shown in B). Pearson’s correlation coefficient was measured using the Fiji-JACoP plugin. Values are expressed as mean ± SD. Three parasites per line were analyzed. **(D)** Quantification of the dot pattern for TgCRMPa-HA_3_ and TgCRMPb-HA_3_ in TgNd9_iKD lines. CRMPs accumulation at the apical tip of extracellular parasites was measured as in figure S4A. TgCRMPs were still found in the apical dot in absence of TgNd9 (+ATc). Numbers are expressed as percentage of parasites showing (dot) or lacking (no dot) the tip accumulation of TgCRMPa and TgCRMPb. The number of parasites (n) analyzed for each line is reported at the top of the column. **(E)** Immunofluorescence and DIC images of parasites in early, middle and late stages of host cell invasion. Parasites co-expressing TgCRMPa-HA_3_ and TgNd6-TY_2_ were incubated with host cell monolayers and fixed after 2, 3, and 5min. Parasites were immunostained with anti-HA, anti-TY and anti-RON5 Abs to label CRMPs, Nd6 and the moving junction, respectively. DNA is labeled by Hoechst. In contrast with TgNd6 (arrow), the apical accumulation of TgCRMPa observed in extracellular parasites, disappears upon entering the host, a step marked by the formation of the moving junction (arrowhead), and for the entire process. Non-specific anti-TY labeling of mitochondria was detected for both untagged and tagged lines (Figure S5E). Single focal planes are shown. **(F)** Model depicting the proposed role for the TgCRMP complex in rhoptry exocytosis. Upon contacting the host cell via a host ligand, CRMPs participate in the activation of the signaling pathway targeting the AV-RSA system, and leading to the fusion events required for the timely discharge of rhoptry contents. Rh: rhoptry; AV: apical vesicle; RSA: rhoptry secretory apparatus.

We then asked whether the apical dot labeled by CRMPs and Nd6 was maintained throughout the entire invasion process or limited to the pre-entry step. We fixed and immunostained parasites co-expressing TgCRMPs and TgNd6, at different time points during the invasion process, and used anti-RON5 antibodies to label the moving junction and mark the progress of invasion. Interestingly, TgNd6 apical labeling was detected when the parasite started entering the host cell and remained visible throughout the entire process, until the parasite was completely inside the host cell (Figures 6E, S5H). However, TgCRMPa apical signal vanished as soon as the moving junction is formed (Figure 6E). The same results were obtained for TgCRMPb (Figure S5H).

To sum up, CRMPs form a complex required to trigger exocytosis that is spatially located in close proximity but distinct from the RSA-associated Nd complex. Both Nd and CRMP complexes have different fates during invasion, adding further support to a model where Nd and CRMP complexes play related but distinct roles in controlling rhoptry secretion at the exocytic site.

## DISCUSSION

Apicomplexan parasites have evolved highly specialized secretory organelles called rhoptries, which are key players for establishing successful infection. Rhoptry secretion is a complex process coupled with host membrane interaction and injection of materials into the host. The underlying mechanisms of this unique cell biological process remain largely unresolved, though hints regarding the exocytic step—the fusion between the rhoptry and parasite plasma membranes—have been recently obtained (Aquilini et al., 2021; Mageswaran et al., 2021). In the present study we took advantage of the relatively close evolutionary relationship between ciliates and apicomplexans, and in particular their sharing unique mechanisms for regulated secretion (Aquilini et al., 2021), to identify new rhoptry secretion factors in *Toxoplasma*. A *Tetrahymena*-based *in silico* screening led us to the identification of a key rhoptry secretion complex comprising TgCRMPa, TgCRMPb, Tg247195 and Tg277910 proteins. Our data suggest that these novel factors link the recognition of the host cell to the activation of the rhoptry exocytic machinery.

We showed that the TgCRMPs are present at the apex of extracellular parasites, the site where the parasite contacts the host cell and discharges its rhoptry content. CRMPs, together with their partners, are membrane proteins containing domains known to interact with proteins and glycans. Both predictions and our experimental data indicate these domains are exposed to the extracellular milieu, and thus likely capable of interacting with host cell membranes. This apical localization relies on the productive assembly of the CRMP-based complex. Moreover, both TgCRMPa and TgCRMPb partially co-localize with TgNd6 at the site of exocytosis. This association is transient since it is evident only in extracellular parasites, and prior to invasion. Once the parasite breaches the host membrane the apical TgCRMPs staining disappears, strongly arguing for a function specifically at the time of rhoptry exocytosis. CRMPs and their partners Tg247195 and Tg277910 are not part of the previously described Nd/NdP exocytic complex. In contrast with the depletion of Nd9, removal of CRMPb does not affect the structural organization of the RSA and Nd6, but not CRMPs, remains at the exocytic site after exocytosis, consistent with distinct roles for CRMPs and Nds/NdPs in the context of rhoptry secretion. This observation, together with translocation of CRMPs to the exocytic site at the time of secretion, and their topology at the membrane with extracellular domains that could interact with the host cell surface, all support a model where CRMPs and their associated factors interact with ligands presented by the host cell. We propose that these interactions activate a signaling cascade within the parasite, leading to rhoptry discharge (Figure 6F). Intriguingly, removal of CRMPb induces slight changes in the shape and anchoring angle of the AV. Albeit the changes are relatively minor, they infer that there could be a direct or indirect connection of CRMPb to the AV which in turn could potentially regulate the rhoptry fusion apparatus.

Our model is also supported by the fact that TgCRMPa and its ortholog PfCRMP1 are predicted to be GPCR-like proteins. Moreover, although there is no clear prediction for *Tetrahymena* TtCRMP1 and TtCRMP2 as GPCRs, they contain a GPCR-Autoproteolysis INducing (GAIN) domain as found by HHpred analysis (score 81.6% for CRMP1 and 95.4% for CRMP2) (Zimmermann et al., 2018)). CRMPs cannot be considered bona fide GPCRs because they do not have the classical seven transmembrane domains of GPCRs (five predicted for *Tetrahymena* CRMPs, and nine for *Toxoplasma* and *Plasmodium* CRMPs), but they may be divergent forms that have maintained similar activities. GPCR is the largest family of membrane-bound receptors known to sense diverse extracellular stimuli and initiate signaling cascades within the cell cytosol to activate cellular responses. GPCRs are involved in nearly all biological processes and represent the favorite therapeutic target for many pathologies (Hauser et al., 2018). Few GPCRs are annotated in the *Toxoplasma* and *P. falciparum* genomes (Fredriksson and Schiöth, 2005; Madeira et al., 2008), suggesting that they are highly divergent and therefore difficult to recognize in Apicomplexa. Because of CRMPs’ potential host-cell-binding domains, they might be analogous to adhesion-GPCRs, a sub-group of proteins with a large extracellular part containing structural modules typical of cell adhesion proteins (Liebscher et al., 2021; Yona et al., 2008). Adhesion-GPCRs convert the stimulus derived from cell-to-cell contact into intracellular signaling via their C-termini, but many lack identified activating ligands. Our data on the localization and topology of TgCRMPs support this scenario, in which the proteins’ N-termini are exposed extracellularly to capture the signal, and the C-termini face the cell cytoplasm to transduce the signal for exocytosis. The N-terminal extensions of the apicomplexan CRMPs are larger than the *Tetrahymena* counterparts and contain, in addition to the EGF receptor domain, both lectin and Kringle domain. This difference in complexity may reflect the need to respond to different stimuli for triggering exocytosis. Ciliates must sense environmental changes to trigger exocytosis, while the parasites response depends on intimate cell-cell contacts: they must interact first with a host to ensure that rhoptry secretion is effective. Interestingly, while *Plasmodium* CRMP1 and CRMP2 are dispensable for merozoite invasion of red blood cells, they are required for sporozoite entry into the salivary glands (Douradinha et al., 2011; Thompson et al., 2007), a step recently shown to be dependent on rhoptry secretion (Ishino et al., 2019). These findings suggest that CRMPs evolved differently to adapt to diverse environments or hosts.

The localization of CRMPs at the apical tip is evident, while their intracellular distribution is less clear. TgCRMPa and TgCRMPb are visible as small dots likely corresponding to vesicles, within the cytosol of intracellular parasites. An apical gradient reminiscent of micronemes was occasionally observed, but CRMPs do not appear to co-localize extensively with microneme markers. Moreover, the hyperLOPIT (spatial proteomics method hyperplexed Localisation of Organelle Proteins by Isotopic Tagging) analysis predicts that CRMPs, Tg247195 and Tg277910 are associated with micronemes when using the TAGM-MCMC method, but this prediction was not supported by data obtained with TAGM-MAP analysis (Barylyuk et al., 2020). These different observations might be reconciled by the long-standing hypothesis of different subsets of micronemes (Kremer et al., 2013). Unfortunately, immuno electron microscopy of TgCRMPa-and TgCRMPb-tagged lines was inconclusive, and thus did not further clarify the distribution of these proteins in intracellular parasites (data not shown).

In this study we described CRMPs as novel secretory factors shared between ciliates and apicomplexans, providing further support to the existence of conserved machinery for secretion in Alveolates. Our previous work showed that the fusion machinery responsible for the discharge of secretory organelles is conserved between Apicomplexa and Ciliata (Nd/NdP proteins). Here, we extend such conservation to the putative signaling pathway leading to exocytosis. CRMPs represent suitable target for new treatments against apicomplexan-related infections. Uncovering the host ligands for CRMPs, Tg247195 and Tg277910 proteins, as well as the signaling pathway downstream of their interaction, will greatly help to develop strategies for blocking rhoptry exocytosis and subsequent invasion, contributing further to fight human infections caused by apicomplexans.

## ACKNOWLEDGEMENTS

We thank Sebastian Lourido for the pU6-Universal plasmid, Dominique Soldati-Favre for providing the anti-ARM (ARO) antibodies and pLinker-2xTy-DHFR plasmid, Nicolas Dos Santos Pacheco for helping in setting up the Ultrastructure Expansion Microscopy, Anita Koshy for the toxofilin-Cre plasmid and Helen Blau’s lab for the Cre reporter DSred cell line. We thank Veronique Richard and Frank Godiard of the MEA platform, University of Montpellier for their assistance with electron microscopy and Pilar Ruga Fahy of the Pôle Facultaire de Microscopie Ultrastructurale, in Geneva for preparation of freeze-fracture replicas. We are also grateful to Elodie Jublanc, Vicky Diakou and the imaging facility MRI at the University of Montpellier, part of the national infrastructure France-BioImaging supported by the French National Research Agency (ANR-10-INBS-04, «Investments for the future»), and Christophe Duperray of the MRI-Cytometry at the Institute for Regenerative Medicine and Biotherapy for their assistance and technical support. Mass spectrometry experiments were carried out using the facilities of the Montpellier Proteomics Platform (PPM, BioCampus Montpellier). We thank Stefan Steimle for his technical assistance with the Krios G3i cryogenic electron microscope; the Singh Center for Nanotechnology and the Beckman Center for Cryogenic Electron Microscopy at the University of Pennsylvania for hosting and supporting the use of the Titan Krios. Dr Maryse Lebrun is an INSERM researcher. This work was supported by the Laboratoire d’Excellence (LabEx) (ParaFrap ANR-11-LABX-0024), and European Research Council (ERC advanced grant number 833309 KissAndSpitRhoptry) to M.L.; by the FACCTS (France and Chicago Collaborating in the Sciences) to A.P.T and M.L.; by NIH GM105783 to A.P.T.; by a David and Lucile Packard Fellowship for Science and Engineering (2019-69645) and a Pennsylvania Department of Health FY19 Health Research Formula Fund to Y.-W.C; by NIH R01 AI112427 to B. S. D.S. and M.M.C. are supported by the European Research Council (ERC) under the European Union’s Horizon 2020 research and innovation program under Grant agreement no. 833309 to M.L.

## AUTHOR CONTRIBUTION

Conceptualization: D.S. M.L. Investigation *Tetrahymena*: L.M.T. performed CDH analysis; D.S. generated and analyzed mutants. Investigation *Toxoplasma*: J.D., J.H., D.M.P.V. generated the tagged and iKD lines; J.D. performed phenotypic analysis of the mutants with the help of M.M., D.M.P.V., J.H.; D.S. generated and analyzed lines for co-IP and fluorescence microscopy of extracellular parasites; D.M.P.V., S.U., performed IP and mass spectrometry analysis; L.B.S., J.H. performed EM analysis; D.S., M.M.C., performed ultrastructure expansion microscopy. Phylogenies: D.S. Freeze fracture data: D.S., J.H. prepared samples; D.S., J.F.D., collected data. Cryo-ET data: A.G. cultured the cells; A.G. and S.K.M. prepared M.L.

## DATA AVAILABILITY

The mass spectrometry proteomics data have been deposited to the ProteomeXchange Consortium via the PRIDE (Perez-Riverol et al., 2019) partner repository with the dataset identifier PXD031161 and PXD031164.

## STAR METHODS

### *Tetrahymena* culture conditions

*Tetrahymena thermophila* strains used in this work are the wildtype CU428.1 (Ctrl), the mutants *Δfer2*, *Δ00442310* (*ΔCRMP1*) and *Δ00637180* (*ΔCRMP2*). Unless otherwise indicated, cells were grown overnight in SPP (2% proteose peptone, 0.1% yeast extract, 0.2% dextrose, 0.003% ferric-EDTA) supplemented with 250 ug/ml penicillin G and 250 ug/ml streptomycin sulfate, to medium density (1-3×10^5^ cells/ml). For biolistic transformation, growing cells were subsequently starved in 10 mM Tris buffer, pH 7.4. Fed and starved cells were both kept at 30°C with agitation at 60 rpm. Culture densities were measured using the Neubauer chamber.

### Homology searching and phylogenetic tree construction for *ferlin* genes

*Toxoplasma gondii* ferlin 2 protein sequence (TgFer2; TGME49_260470) was used as query against translated open reading frame (ORF) coding sequences from the genomes of selected Ciliata and Apicomplexa species using the BLAST algorithm. Positive BLAST hits against TgFer2 query were those with an E value < 0.001 and score >50, for which reciprocal BLAST against the genome containing the query sequence retrieved either the same sequence or an isoform of the sequence with similar E value or lower. Once homologs of TgFer2 were identified by BLAST, the phylogenetic relationships of ferlins within ciliates and apicomplexans were determined by maximum likelihood estimation. Homologs were aligned by MUSCLE and tree construction was performed by MEGA-X software (Kumar et al., 2018). The root of the tree was determined using *Toxoplasma* ferlin 3 (TgFer3) as outgroup. Ciliata and Apicomplexa sequences were retrieved from either the http://ciliate.org or http://veupathdb.org databases, respectively. Identification numbers and E-values of the proteins used for tree construction are reported in Table S4.

### Coregulation Data Harvester (CDH) analysis

The identification of co-regulated genes for *Tetrahymena Fer2* (TTHERM_00886960), *Nd6* (TTHERM_00410160), *NdP1* (TTHERM_01287970) and *NdP2* (TTHERM_00498010) was performed by Coregulation Data Harvester (CDH) software (http://ciliate.org/index.php/show/CDH) as previously described (Tsypin and Turkewitz, 2017). A list of co-regulated genes, with homologs in *T. gondii* and *P. falciparum*, was obtained for each query. We then generated a cross-list by selecting genes shared at least by three queries, for which reciprocal BLAST towards ToxoDB and PlasmoDB databases retrieved *Toxoplasma* and *Plasmodium* homologous genes with an E-value of at least 10^-4^, respectively (Table S1).

### Generation of *Tetrahymena* knockout strains

*ΔFer2*, *ΔCRMP1*, and *ΔCRMP2* mutants were generated by replacing the macronuclear ORF of *Fer2* (TTHERM_00886960), *CRMP1* (TTHERM_00442310) and *CRMP2* (TTHERM_00637180), with the paromomycin (Neo4) drug resistance cassette (Mochizuki, 2008) via homologous recombination with the linearized vectors p00886960-Neo4, p00442310-Neo4, and p00637180-Neo4. To generate the knockout constructs, 500-800bp fragments homologous to the genomic regions upstream (5’UTR) and downstream (3’UTR) of each ORF were PCR amplified with KOD HiFi polymerase (Merk), and cloned into SacI/PstI and XhoI/HindIII restriction sites, respectively, flanking the Neo4 cassette in the pNeo4 plasmid by Quick Ligation (New England, Biolabs Inc.). Specifically, for p00886960-Neo4, 707bp (5’UTR) and 754bp (3’UTR) homology regions (HRs) were amplified with primers ML4379/ML4380 and ML4381/ML4382, respectively; for p00442310-Neo4, 569bp (5’UTR) and 721bp (3’UTR) were amplified with primers ML3830/ML3831 and ML3832/ML3833, respectively; for p00637180-Neo4, 727bp (5’UTR) and 506bp (3’UTR) were amplified with primers ML4283/ML4284 and ML4285/ML4286, respectively. Each construct was linearized by digestion with SacI and KpnI, and delivered into CU428.1 cells by biolistic transformation. Primers are listed in Table S5.

### *Tetrahymena* biolistic transformation

*Tetrahymena* CU428.1 was grown to mid log phase and starved 18-24h in 10mM Tris, pH 7.4. Biolistic transformation was performed with 20µg synthetic DNA as described previously (Cassidy-Hanley et al., 1997; Chilcoat et al., 1996). Selection of positive transformants was initiated 5h after bombardment by adding 120µg/ml paromomycin sulfate and 1µg/ml CdCl_2_ to the cultures. Transformants were serially transferred 6x/week in increasing concentrations of drug and decreasing concentrations of CdCl_2_ (up to 3mg/ml paromomycin and 0.2 µg/ml CdCl_2_) for at least 6 weeks before further testing. Successful integration and replacement of all endogenous alleles at each genomic locus was tested by RT-PCR.

### RT-PCR assessment of gene disruption in *Tetrahymena*

Overnight cultures of mid log phase cells from each knockout strain were pelleted, washed once with 10mM Tris pH 7.4, and total RNA was isolated using NucleoSpin RNA, Mini kit for RNA purification (Macherey-Nagel), according to manufacturer’s instructions. The cDNA synthesis from 2-3µg of total RNA was performed using High-Capacity cDNA Reverse Transcription Kit (Applied Biosystems). The cDNA was PCR amplified with GoTaq DNA Polymerase (Promega) to assay the presence of the corresponding transcripts (200-300bp) in the knockout strains using primers listed in Table S5. To confirm that equal amounts of cDNA were amplified, reactions with primers specific for β-tubulin 1 (BTU1) were run in parallel. At least three clones each for the knockout strains were tested.

### *Tetrahymena* mucocysts secretion assay

Wildtype CU428.1 and knockout strains were grown to stationary phase (10^6^ cells/ml) in 30ml SPP for 48h, and then concentrated into 500µl loose pellet by centrifugation at 1800g for 3min. Cells were stimulated with 165µl of 25mM dibucaine, vigorously mixed for 30s and diluted to 15ml with 10mM HEPES pH 7.4, and 5mM CaCl_2_. Samples were then centrifuged at 1800g for 3min resulting in the formation of a cell pellet/flocculent bilayer. Quantification of exocytic competence was performed by measuring the ratio between flocculent layer and pellet volumes. At least three clones each for the knockout strains were tested.

### *Tetrahymena* western blotting

Whole cell lysates were collected from 5×10^4^ cells from overnight cultures, washed once with 10mM Tris pH 7.4, resuspended in 2x lithium dodecyl sulfate (LDS) sample buffer containing 40mM DTT, and denatured at 95°C for 10 min. Proteins were resolved with the Novex NuPAGE Gel system (10% Bis-Tris gels, Invitrogen) and transferred to 0.45µm PVDF membranes (Immobilon^®^-P, Millipore). Blots were blocked with 5% dried milk in 1x TNT (15 mM Tris, 140 mM NaCl, 0.05% Tween 20, pH 8). The rabbit anti-Grl1 serum (Turkewitz et al., 1991) was diluted 1:2000 in blocking solution. Proteins were visualized with anti-rabbit alkaline phosphatase (AP)-conjugated (Promega) diluted 1:7500 and with BCIP/NBT Color development substrate (Promega). At least three clones each for the knockout strains were tested.

### *Tetrahymena* immunofluorescence microscopy

Overnight cultures of mid-log phase *Tetrahymena* cells for CU428.1 (control), *ΔFer2*, *Δ00442310* (*ΔCRMP1*) and *Δ00637180* (*ΔCRMP2*) were washed once with 10mM Tris pH 7.4, and fixed with 4% paraformaldehyde (PFA) in 50mM HEPES pH 7.4 at room temperature. Cells were permeabilized with 0.1% Triton X-100 and blocking was performed with 1% bovine serum albumin (BSA) in TBS (25 mM Tris, 3 mM KCl, 140 mM NaCl, pH 7.4); mucocyst proteins Grl3 were visualized with mouse mAb 5E9 (1:10) (Cowan et al., 2005) followed by AlexaFluor488 goat anti-mouse antibody (1:450) (Invitrogen), both diluted in 1% BSA. Cells were mounted in 30% glycerol/TBS and imaged on a Leica Thunder microscope, with a 100x oil objective NA=1.4, equipped with the sCMOS 4.2MP camera, using Leica Application Suite X (LAS X) software (Leica Biosystems). Z-stacks were denoised, adjusted in brightness and contrast, and colored with the program Fiji (Schindelin et al., 2012). At least two clones each for the knockout strains were tested.

### *Toxoplasma* culture conditions

*Toxoplasma gondii* RH tachyzoites (type I strain) lacking the *Ku80* gene (*ΔKu80*) (Huynh and Carruthers, 2009) were used for genetic recombination. In particular, to generate inducible knockdown strains, we used either the *ΔKu80* line expressing the TATi transactivator for the TetOff System (*TATi-ΔKu80*) (Sheiner et al., 2011), or the Tir-1 receptor for the auxin inducible degron system (miniAID) (Brown et al., 2018) (*ΔKu80* Tir-1). Parasites were routinely cultured in Human Foreskin Fibroblasts (HFFs) monolayers (ATCC, CRL 1634) in standard medium (DMEM 5% Fetal Bovine Serum -FBS-, 2 mM glutamine, supplemented with penicillin and streptomycin from Gibco) at 37°C and 5% CO_2_. For SeCreEt Assays, parasites expressing the protein toxofilin fused with a Cre-recombinase (Koshy et al., 2010) were cultured in mouse fibroblast cell line 10T1/2, constitutively expressing a floxed red fluorescent protein DsRed (Koshy et al. 2010), used as Cre-reporter cell line for assessing rhoptry secretion. Parasites used for immunoprecipitation experiments were cultured in Vero Cells (ATCC, CCL 81) with DMEM 3% FBS supplemented with glutamine, penicillin and streptomycin. For positive selection via hypoxanthine-xanthine-guanine phosphoribosyl transferase (HXGPRT) drug resistance cassette, 25µg/ml mycophenolic acid plus 50µg/ml xanthine were added to the culture media; 2µM pyrimethamine and 20µM chloramphenicol (CHL) were used for selection with the Dihydrofolate Reductase Thymidylate Synthase (DHFR-TS), and Chloramphenicol Acetyl Transferase (CAT) drug resistance cassettes, respectively. For negative selection via Uracil Phosphoribosyl Transferase (UPRT) cassette, 5µM fluorodeoxyuridine (FUDR) was added to the medium. To induce protein depletion in the iKD lines, 1µg/ml anhydrotetracycline (ATc) (Fluka 37919) was added to the medium for 24h, 48h and 72h, depending on the strain.

### Generation of *Toxoplasma* tagged and knockdown strains

All *Toxoplasma*-related primers and RNA guides (gRNAs) used in this study are listed in table S5.

Genomic DNA was isolated using Wizard SV genomic DNA purification system (Promega). KOD HiFi polymerase (Merk) and GoTaq DNA Polymerase (Promega) were used to amplify gene fragments for cloning strategy, and for colony screening PCRs, respectively.

*TgCRMPa* (TGGT1_261080) and *TgCRMPb* (TGGT1_292020) were C-terminally fused with a triple hemagglutinin (HA_3_) tag followed by the chloramphenicol resistance cassette (CAT) for selection, in the *TATi-ΔKu80* line using CRISPR/Cas9. Briefly, gRNAs targeting the 3’UTR of the genes were generated by annealing primers ML3283/ML3284 and ML3279/ML3280 respectively. The annealed gRNAs were cloned in the pU6-Cas9-YFP plasmid using the BsaI restriction sites, to generate pU6-TgCRMPa_gRNA1 and pU6-TgCRMPb_gRNA1. DNA fragments containing gene-specific homologous regions flanking the triple HA tag and the CAT cassette, were amplified from pLIC_HA3_CAT vector (Huynh and Carruthers, 2009) using the primer pairs ML3287/ML3288 and ML3277/ML3278 for *TgCRMPa* and *TgCRMPb* respectively, containing ∼30bp of homology to the 3’ and 3’UTR of the gene of interest. pU6-TgCRMPa_gRNA1 and pU6-TgCRMPb_gRNA1 plasmids and the corresponding donor DNAs were mixed prior to be transfected. The resulting lines were named TgCRMPa-HA_3_ and TgCRMPb-HA_3_.

Tg277910 was tagged with a triple HA tag (HA_3_) at the C-terminal end of the protein in the *TATi-ΔKu80* line using the Ligation Independent Cloning (LIC) strategy (Huynh and Carruthers, 2009) and chloramphenicol selection. Briefly, 1485bp fragment corresponding to the 3’ part of TGGT1_277910 gene minus the stop codon, was amplified with primers ML4046/ML4047 and integrated in the pLIC_HA3-CAT (Huynh and Carruthers, 2009). The vector was then linearized with BaeI site prior to transfection. The tagged line was named Tg277910-HA_3_.

Inducible knockdowns (iKDs) of *TgCRMPa*, *TgCRMPb* and *Tg277910* were generated in TgCRMPa-HA_3_, TgCRMPb-HA_3_ and Tg277910-HA_3_ lines, respectively, using pyrimethamine selection; the resulting strains were named *TgCRMPa*_iKD, *TgCRMPb*_iKD and *Tg277910*_iKD. To create the iKD lines, the endogenous promotor of each gene was replaced by an anhydrotetracycline (ATc)-regulatable promotor (TetO7SAG4), preceded by the *DHFR* cassette, using CRISPR/Cas9, as described previously (Suarez et al., 2019). gRNAs targeting the 5’UTR of the genes were generated by annealing the primer pairs ML3342/ML3343, ML3338/ML3339 and ML3970/ML3971, for *TgCRMPa*, *TgCRMPb* and *Tg277910*, respectively, and introduced in the BsaI site of pU6-Cas9-YFP, to generate pU6-TgCRMPa_gRNA3, pU6-TgCRMPb_gRNA3 and pU6-Tg277910_gRNA1. Donor DNA fragments were obtained by amplifying the TetO7SAG4 promoter and the DHFR resistance cassette from the DHFR-TetO7SAG4 plasmid (Sheiner et al., 2011) with the following primers: ML3317/ML3318 (*TgCRMPa*_iKD), ML3315/ML3316 (*TgCRMPb*_iKD) and ML3966/ML3967 (*Tg277910*_iKD), respectively. Each pair of primers contains ∼30 bp of homology to the 5’UTR and to the 5’ coding region of the gene. The gRNAs and donor DNAs were mixed prior to parasite transfection.

Inducible auxin-inducible knockdown of *TgCRMPa* was generated in Tir-1-expressing line. *TgCRMPa* was C-terminally fused with the miniAID sequence followed by a triple HA tag. The resulting line was named TgCRMPa-miniAID-HA_3__iKD. A fragment containing the miniAID sequence followed by the *HXGPRT* cassette was amplified from the plasmid pTUB8YFP-mAID-3 (Brown et al., 2017) with primers ML4909/ML4910, and mixed with the pU6-TgCRMPa_gRNA1.

To tag TgCRMPa with a triple FLAG at the C-terminus, we used a marker-free strategy. Integration of the tag at the endogenous locus was achieved by homologous recombination of a 500bp DNA fragment (gBlock, Genescript) containing the triple FLAG tag flanked by 185bp and 228bp of homology to the 3’ coding sequence and 3’UTR of *TgCRMPa*, respectively. The 500bp donor DNA was amplified from the synthetic gBlock with primers ML3002/ML3003, and mixed with the pU6-Cas9-YFP plasmid containing pU6-TgCRMPa_gRNA1. *TgCRMPa* was fused with the triple FLAG tag in the *TATi-ΔKu80* and TgCRMPb-HA_3_ lines, which were then used to generate *TgCRMPa* knockdown lines as described earlier for *TgCRMPa*_iKD. The lines generated from *TATi-ΔKu80* were named TgCRMPa-FLAG_3_ and TgCRMPa-FLAG_3__iKD, those generated from TgCRMPb-HA_3_ were named TgCRMPa-FLAG_3_+TgCRMPb-HA_3_ and TgCRMPa-FLAG_3__iKD+TgCRMPb-HA_3_.

C-terminal tagging of TgNd6 with double TY tag in TgCRMPa-HA_3_ and TgCRMPb-HA_3_ lines was obtained by inserting the coding sequence of TY_2_, followed by the DHFR resistance cassette, immediately after the gene’s stop codon in the *TgNd6* locus. The gRNA primers ML3129/ML3130 tagging the 3’UTR of *TgNd6*, were annealed and then cloned into the pU6-Cas9-YFP plasmid using the BsaI restriction sites to generate pU6-TgNd6_CtgRNA. The plasmid was mixed with 4597bp donor DNA fragment prior to transfection. The 4597bp donor DNA was PCR-amplified from the pLinker-2xTy-DHFR plasmid (Suarez et al., 2019) with primers ML4734/ML4735, and mixed with pU6-TgNd6_CtgRNA prior to transfection. The lines were named TgCRMPa-HA_3_+TgNd6-TY_2_ and TgCRMPb-HA_3_+TgNd6-TY_2_. C-terminal HA_3_ tagging of *TgCRMPA* and *TgCRMPB* in *TgNd9*_iKD (Aquilini et al., 2021) was obtained as described earlier for TgCRMPA-HA_3_ and TgCRMPB-HA_3_ lines. To quantify rhoptry secretion using SeCreEt assays, the Toxofilin-Cre recombinase was introduced in *TATi-ΔKu80*, *TgCRMPa*_iKD, *TgCRMPb*_iKD and *Tg277910*_iKD lines at the uracil phosphoribosyl transferase (UPRT) locus, to generate *TATi-ΔKu80*_Toxofilin-Cre*, TgCRMPa*_iKD_Toxofilin-Cre, *TgCRMPb*_iKD_Toxofilin-Cre and *Tg277910*_iKD_Toxofilin-Cre. Toxofilin-Cre (3949bp) was amplified from ToxofilinCre plasmid (Koshy et al., 2010) using primers ML3522/ML3523, containing ∼30 bp of homology to the 5’ and 3’UTR of the *UPRT* gene and co-transfected with two specific single gRNAs cutting the 5’ (ML3445/ML3446) and 3’ (ML2087/ML2088) of the *UPRT* gene.

### *Toxoplasma* transfection and screening of positive transformants

For *T. gondii* transfection, 60µg of pLIC plasmid for Tg277910-HA_3_, or 100µl of purified PCR products (∼5µg) mixed with 15-20µg of corresponding pU6-CAS9-YFP plasmids, were introduced in 20×10^6^ tachyzoites by electroporation, using Electro Cell Manipulator 630 (BTX) with the following settings: 2.02 kV, 50 Ω, 25µF (Kim et al., 1993). After transfection, positive transformants were recovered by drug selection and clones were isolated by limiting dilution, or by fluorescence-activated cell sorting (FACS). Genomic DNA from isolated clones was purified as described earlier, and screened by PCR for correct integration with GoTaq DNA Polymerase (Promega). Alternatively, PCR screening of single clones directly from 96-well plates was performed with Phire^TM^ Tissue Direct PCR master mix (Thermo Scientific) protocol, as previously described (Piro et al., 2020). The primers used to test correct integration are listed in Table S5.

### Homology searching and phylogenetic tree construction for *CRMP* genes in Apicomplexa

*Toxoplasma gondii* CRMPa (TGME49_261080) and CRMPb (TGME49_292020) protein sequences were used as queries against translated ORF coding sequences from the genomes of selected Apicomplexa (see Table S4) using the BLAST algorithm. Positive BLAST hits against CRMPa and CRMPb queries were those for which reciprocal BLAST against the genome containing the query sequence, retrieved the same sequence with similar E-value or lower followed by the other TgCRMP. Once the homologs of TgCRMPa and b were identified by BLAST, the phylogenetic relationships of CRMPs within apicomplexans were determined as described earlier for ferlins. Apicomplexa sequences were retrieved from http://veupathdb.org databases. Identification numbers and E-values of the proteins used for tree construction are reported in Table S4.

### RT-PCR assessment of *Nd9* transcripts depletion in *Toxoplasma*

TgNd9_iKD and TATi-*ΔKu80* parasites were treated 72h with ATc and total RNA was isolated and reverse-transcribed as mentioned earlier for *Tetrahymena* samples. Untreated parasites were analyzed in parallel. The cDNA was PCR amplified with GoTaq DNA Polymerase (Promega) to assay the presence of the corresponding *Nd9* transcripts (∼250bp) in the knockdown strain using primers listed in Table S5. To confirm that equal amounts of cDNA were amplified, reactions with primers specific to TgGAPDH were run in parallel.

### *Toxoplasma* rhoptry secretion assay

To assess parasites competence for rhoptry secretion, SeCreEt (<u>Se</u>creted <u>Cre E</u>pitope-tagged) parasites expressing the toxofilin-Cre fusion protein were generated as described earlier, and used to infect murine fibroblasts (cell line 10T1/2) constitutively expressing a floxed red fluorescent protein DsRed. This mammalian Cre-reporter cell line is able to switch from DsRed to eGFP (enhanced green fluorescent protein) expression upon toxofilin-driven Cre-mediated recombination (Koshy et al., 2010). DsRed cells were grown to a density of 2×10^5^ cells/ml and infected in absence of ATc, with either ATc-pretreated (48h, 24h and 72h ATc incubation for *TgCRMPa*_iKD_Toxofilin-Cre, *TgCRMPb*_iKD_Toxofilin-Cre and *Tg277910*_iKD_Toxofilin-Cre, respectively) or untreated tachyzoites, at a multiplicity of infection (MOI) of 3. One day post-infection, infected cells were trypsinized, and DsRed and eGFP fluorescence signals were measured by fluorescence-activated cell sorting (FACS). The numbers of DsRed and GFP positive cells were used as measure of impaired or successful rhoptry secretion, respectively; the values were reported as fraction of GFP-positive cells over the total number of cells, and expressed as percentages. Each value was normalized to that of the control line (*TATi-ΔKu80*_Toxofilin Cre +ATc) arbitrarily fixed to 100%. Graphs show the mean of three independent experiments.

### *Toxoplasma* invasion assay

For the quantification of invasion in *TgCRMPa*_iKD, *TgCRMPb*_iKD and *Tg277910*_iKD lines, freshly egressed tachyzoites (5×10^6^/coverslips), ATc-pretreated (48h, 24h and 72h ATc incubation for *TgCRMPa*_iKD, *TgCRMPb*_iKD and *Tg277910*_iKD, respectively) or untreated, were added to HFF monolayers grown on coverslips in a 24-well plate, and let settle on ice for 20 min, prior to be transferred to a 38°C preheated water bath for 5 min, to allow invasion. Parasites were fixed with 4% PFA in Hank’s Balanced Salt Solution (HBSS) for 20 min at room temperature, and incubated with 10% FBS/HBSS blocking solution. In order to distinguish intracellular from extracellular parasites, a dual-antibody staining was performed as previously described (Cerede et al., 2005). First, non-permeabilized extracellular parasites were stained using the mouse mAbs T41E5 anti-SAG1 (1:2000) (Couvreur et al., 1988) in 2% FBS/HBSS. Parasites and infected cells were then permeabilized with 0.1% saponin, and incubated again with blocking solution. Secondly, parasites were stained with rabbit anti-ROP1 antibodies (1:3000) (Lamarque et al., 2014) in 2% FBS/PBS, to label intracellular parasites in parasitophorous vacuoles. Secondary antibody staining was performed with AlexaFluor594 goat anti-mouse (1:4000) and AlexaFluor488 goat anti-rabbit (1:10000) antibodies (Invitrogen). DNA was stained with 16µM Hoechst, and coverslips were mounted onto microscope slides using Immunomount (Calbiochem). Intracellular parasites were counted in 20 fields/coverslip (n=3 coverslips/experiment) with a Leica DM2500, 100x oil objective NA=1.4, microscope (Leica Biosystems). The values were expressed as the number of intracellular parasites per field and normalized to that of the control line (*TATi-ΔKu80* -ATc) arbitrarily fixed to 100%. Graphs show the mean of three independent experiments.

### *Toxoplasma* host-cell-attachment assay

The ability of TgCRMPa_iKD, TgCRMPb_iKD and Tg277910_iKD tachyzoites to attach to host cells, was assessed as previously described (Aquilini et al., 2021). HFFs grown on coverslips in a 24-well plate were fixed with cold 2% glutaraldehyde/PBS for 5 min at 4°C, washed three times with cold PBS, quenched with 100mM cold glycine for 2 min, and washed three more times in PBS, and then kept in preheated DMEM 5% FBS. Coverslips were infected with 1×10^7^ freshly egressed ATc-treated (48h, 24h and 72h ATc incubation for *TgCRMPa*_iKD, *TgCRMPb*_iKD and *Tg277910*_iKD, respectively) and untreated parasites resuspended in 300µl DMEM 5% FBS. Coverslips were also infected with parasites pre-treated with 20µM BAPTA-AM (Sigma), used as negative control for microneme-dependant attachment. Parasites were allowed to attach to host cells for 20 min at 37°C, carefully washed twice with DMEM 5% FBS and then fixed with 4% PFA/PBS for 30 min. Parasites were incubated with 1.5% BSA/PBS blocking solution and subsequently immunostained with mouse anti-SAG1 p30 hybridoma (1:50) (Couvreur et al., 1988) followed by secondary staining with AlexaFluor488 goat anti-mouse antibodies (1:4000) (Invitrogen), both diluted in 0.15% BSA/PBS. DNA was stained with 16µM Hoechst, and coverslips were mounted onto microscope slides using Immunomount (Calbiochem). Parasites were counted in 20 fields/coverslip (n=3 coverslips/experiment) with a Leica DM2500, 100x oil objective NA=1.4, microscope (Leica Biosystems). Attachment was reported as number of parasites found attached to the host cells per field and expressed as percentage. Values were normalized to that of the control line (iKD -ATc) arbitrarily fixed to 100%. Graphs show mean values of three independent experiments.

### *Toxoplasma* plaque assay

TgCRMPa_iKD, TgCRMPb_iKD and Tg277910_iKD tachyzoites pre-treated with ATc for 48h, 24h or 72h, respectively, were used to infect HFF monolayers grown in 24-well plates. Untreated parasites were analyzed in parallel. Roughly 3500 parasites were added in each well of the first row, then serial dilutions were performed by transferring ¼ of the parasites to the next well, and so on until the end of the plate. Lysis plaques formation was allowed for 7 days at 37°C and 5% CO_2_. HFFs were then fixed with 4% PFA/PBS and stained with 1% Crystal-violet. Images of the lysis plaques were collected with Olympus MVX10 macro zoom microscope, equipped with a Zeiss MRM2 Camera. Plaques area were measured using ImageJ (Schneider et al., 2012) and expressed as percentage. Values were normalized to that of the control line (*TATi-ΔKu80* -ATc) arbitrarily fixed to 100%. Graphs shows the mean of three independent experiments, including three technical replicates each.

### *Toxoplasma* replication assay

*In vitro* growth of intracellular parasites was performed as previously described (Aquilini et al., 2021). HFF monolayers grown on coverslips were infected with 2×10^5^ freshly egressed parasites, pre-treated with ATc (48h, 24h and 72h ATc incubation for *TgCRMPa*_iKD, *TgCRMPb*_iKD and *Tg277910*_iKD, respectively) or untreated. Invasion was allowed for 2h, HFFs were then washed five times with HBSS to remove extracellular parasites incapable of entering host cells, and intracellular parasites were let replicate for 24h at 37°C. Cells were then fixed using 4%PFA/PBS and permeabilized with 0.1% Triton X-100; parasites were immunostained with mouse mAbs T41E5 anti-SAG1 (1:2000) (Couvreur et al., 1988) and AlexaFluor488 goat anti-mouse (1:4000) (Invitrogen) antibodies diluted in 2% FBS/PBS. DNA was stained with 16µM Hoechst, and coverslips were mounted onto microscope slides using Immunomount (Calbiochem). The number of vacuoles containing either 2, 4, 6, 8, 16 or 32 parasites were counted using a Leica DM2500, 100x oil objective NA=1.4, microscope (Leica Biosystems), and expressed as percentage. 200 vacuoles/coverslips (n=3 coverslips/experiment) were analyzed. Graphs show means of two and three independent experiments for TgCRMPa_iKD and TgCRMPb_iKD, and Tg277910_iKD, respectively, including three technical replicates each.

### *Toxoplasma* egress assay

To assess the egress of *T. gondii* tachyzoites from host cells, HFF monolayers grown on coverslips were infected with 1×10^5^ parasites and incubated at 37°C for 30h. Egress was induced by stimulating microneme secretion and parasite motility with 3µM A23187 for 8 min. Parasites were fixed with 4% PFA/PBS, permeabilized with 0.1% Triton X-100, incubated with 10% FBS/PBS blocking solution and stained with mouse anti-GRA3 hybridoma (1:100) (Achbarou et al., 1991a) and rabbit anti-GAP45 (1:9000) (Frenal et al., 2014) antibodies, followed by AlexaFluor488 goat anti-mouse (1:4000) and AlexaFluor594 goat anti-rabbit (1:4000) antibodies (Invitrogen), respectively. Upon egress, the PVM is ruptured and GRA3 proteins are released from the PV space into the extracellular milieu. Thus, egress events were analysed by quantifying the presence of GRA3 staining in intact and ruptured PVs, 200 vacuoles/coverslip were analyzed (n=3 coverslips/experiment) with a Leica DM2500, 100x oil objective NA=1.4, microscope (Leica Biosystems). Values were reported as the fraction of ruptured vacuoles over the total number of vacuoles observed, and were expressed as percentages. Values were normalized to that of the control line (*TATI-ΔKu80* -ATc) arbitrarily fixed to 100%. Graphs show means of three independent experiments for TgCRMPa_iKD and TgCRMPb_iKD, and two independent experiments for Tg277910_iKD.

### *Toxoplasma* microneme secretion assay

The extent of microneme secretion in TgCRMPa_iKD, TgCRMPb_iKD and Tg277910_iKD lines was measured by evaluating the release of the TgMIC2 processed form in the supernatant. Freshly egressed, ATc-treated (48h, 24h and 72h ATc incubation for *TgCRMPa*_iKD, *TgCRMPb*_iKD and *Tg277910*_iKD, respectively) and untreated parasites were harvested by centrifugation at 600 g, washed twice in preheated intracellular buffer (5mM NaCl, 142mM KCL, 1mM MgCl2 2mM EGTA, 5.6mM glucose and 25mM HEPES, pH 7.2) and resuspended in DMEM (minus FBS) with or without 500µM propranolol (Sigma; P0884). Parasites were then incubated at 37°C for 20 min to induce the secretion of microneme contents into the supernatant. Supernatants were separated from parasite pellets by centrifugation at 2000g for 5 min at 4°C. Pellets were washed once in PBS and supernatants were additionally cleared by centrifugation at 4000g for 5 min. 1/10 and 1/5 of the total pellets and supernatants were subjected to SDS-PAGE and western blotting, respectively; full-length (∼115kDa) and processed TgMIC2 (∼100kDa) proteins were detected with mouse anti-MIC2 hybridoma (1:2) (Achbarou et al., 1991b) and Horseradish Peroxidase (HRP)-conjugated goat anti-mouse (1:10000) (Jackson Immuno Research) secondary antibodies. TgGRA3 proteins were used as loading control and detected with rabbit anti-GRA3 primary (1:500) (Achbarou et al., 1991a) and anti-rabbit alkaline-phosphatase (AP)-conjugated (1:7500) (Promega) secondary antibodies. Proteins were visualized with BCIP/NBT Color development (Promega) or Clarity Max^TM^ Western ECL (Bio-Rad) substrates. One representative experiment is shown for the iKD lines.

### *Toxoplasma* immunofluorescence microscopy

Unless otherwise specified, immunofluorescence assays (IFAs) of intracellular parasites were performed as previously described (El Hajj et al., 2008). Briefly, coverslips containing infected HFF monolayers were fixed with 4% PFA/PBS for 30 min at room temperature. Cells were washed three times with PBS, permeabilized with 0.15% Triton X-100/PBS for 10 min and then saturated with 10% FBS/PBS blocking solution for 1 h. Proteins were stained with primary antibodies for 1h, followed by six washes with PBS and secondary staining with proper fluorophore-conjugated antibodies for 1h. Antibodies were diluted in 2% FBS/PBS. HA_3_-tagged proteins were visualized with rat anti-HA 3F10 (1:1000) (Roche; 11867460001) or rabbit anti-HA (1:5000) (Abcam; ab9110) and AlexaFluor488 goat anti-rat (1:2000) or goat anti-rabbit (1:10000) (Invitrogen) antibodies; TgARO, TgAMA1, TgMIC2, TgGAMA and TgPLP1 were visualized with rabbit anti-ARM(ARO) (1:1000) (Mueller et al., 2013), rabbit anti-AMA1 folded (1:5000) (Lamarque et al., 2014), mouse anti-MIC2 hybridoma (1:50) (Achbarou et al., 1991b), rabbit anti-GAMA (1:500) (Huynh and Carruthers, 2016) and rabbit anti-PLP1 (1:500) (Roiko and Carruthers, 2013) primary antibodies, respectively, together with AlexaFluor594 goat anti-rabbit (1:4000) or goat anti-mouse (1:4000) (Invitrogen) secondary antibodies.

For detecting the apical accumulation of TgCRMPa-HA_3_ and TgCRMPb-HA_3_, either alone or in pairwise combination with TgNd6-TY_2_, 5×10^6^ parasites/condition were added to HFF monolayers grown on coverslips in a 24-well plate, and let settle on ice for 20 min, prior to be transferred to a 38°C preheated water bath to allow invasion. According to the experimental design, parasites were fixed with 4% PFA/PBS after 2, 3 and 5min incubation at 38°C. Fixation was allowed for 30 min at room temperature prior to permeabilization with 0.1% Triton X-100, blocking with 10% FBS/PBS and antibody staining of extracellular and invading parasites. Rat anti-HA 3F10 (1:1000) primary (Roche; 11867460001) and AlexaFluor488 goat anti-rat secondary (1:2000) (Invitrogen) antibodies diluted in 2% FBS/PBS, were used to visualize TgCRMPa-HA_3_ and TgCRMPb-HA_3_ alone, in *TATi-ΔKu80*, TgCRMPa_iKD and TgNd9_iKD backgrounds. The co-staining of TgCRMPa-HA_3_ and TgCRMPb-HA_3_ with TgNd6-TY_2_ was performed with rabbit anti-HA (1:5000) (Abcam; ab9110) and mouse anti-TY hybridoma (1:100) (Bastin et al., 1996) primary antibodies, followed by AlexaFluor488 goat anti-rabbit (1:10000) and AlexaFluor594 goat anti-mouse (1:4000) secondary antibodies (Invitrogen), diluted in 2% FBS/PBS. Intracellular parasites co-expressing TgCRMPs and TgNd6-TY were similarly stained.

For the time-course experiment during invasion, the co-staining of TgCRMPa-HA_3_ and TgCRMPb-HA_3_ with TgNd6-TY_2_ with primary antibodies was performed as described earlier, followed by AlexaFluor647 goat anti-rabbit (1:2000) and AlexaFluor488 goat anti-mouse highly cross adsorbed (HCA)(1:4000) (Invitrogen) secondary antibodies diluted in 10% FBS/PBS; to visualize the moving junction, parasites were incubated again with 10% FBS/PBS blocking solution, and stained with rat anti-RON5 (1:200) (Besteiro et al., 2009) followed by AlexaFluor594 goat anti-rat HCA (1:2000) (Invitrogen) antibodies. The use of highly cross-adsorbed (HCA) secondary antibodies limited cross-reactivity. DNA was stained with 16µM Hoechst, and coverslips were mounted onto microscope slides using Immunomount (Calbiochem). Imaging was performed either with a Leica Thunder microscope, with a 100x oil objective NA=1.4, equipped with the sCMOS 4.2MP camera, using Leica Application Suite X (LAS X) software (Leica Biosystems), or Zeiss Axioimager Z2 epifluorescence microscope, with a 100x oil objective NA=1.4, equipped with the CMOS Orca Flash 4.0 (Hammamatsu) camera, using Zen software (Zeiss, Intelligent Imaging Innovations), or Zeiss LSM880 confocal microscope equipped with Airyscan detector, with a 63x oil objective NA=1.4, using Zen Black software (Zeiss, Intelligent Imaging Innovations). Images of single focal planes and z-stacks were uniformly denoised, adjusted in brightness and contrast, and colored with the program Fiji (Schindelin et al., 2012). Images were collected at the Montpellier Ressources Imagerie (MRI) facility of the University of Montpellier.

### Co-localization analysis of *Toxoplasma* CRMPs with Nd6 and microneme proteins

To estimate the extent of co-localization, the Fiji-JACoP plugin was used to calculate Pearson’s correlation coefficient (PCC) (Bolte and Cordelieres, 2006). The overlap between CRMPs and Nd6 signals at the apical dot of A23187-treated extracellular parasites, was measured by creating a binary mask of the selected area covering the entire volume of the parasite at the extreme apex. PCC was calculated setting the threshold to the estimated value of background. Z-stacks of three parasites for each line were analyzed. The overlap between CRMPb-HA_3_ and the microneme proteins AMA1, MIC2, GAMA and PLP1 in intracellular parasites was measured as described above. Untagged parasites, equally stained with anti-HA Abs in pairwise combination with the four anti-MICs Abs, were analyzed in parallel to estimate the background noise. Untagged parasites were also stained with anti-AMA1 and anti-MIC2 antibodies to measure the overlap between the two microneme proteins. 20-32 parasites were analyzed for each pair of antibodies.

### Western blotting of *Toxoplasma* proteins

For western blotting of whole cell lysates, ∼10^7^ freshly egressed tachyzoites/condition were washed once in PBS and resuspended in 100°C Laemmli SDS or lithium dodecyl sulfate (LDS) sample buffer supplemented with 10-40mM dithiothreitol (DTT) (Sigma). Epitope-tagged TgCRMPa, TgCRMPb, Tg277910 and related proteins were resolved with either the Bio-Rad Gel system (10% gel: 10% acrylamide/bis, 0.4M Tris-HCl pH 8.8, 0.1% SDS, 0.1% APS, temed) or the Novex NuPAGE Gel system (3-8% Tris-Acetate gels, Invitrogen) and transferred either to 0.45µm nitrocellulose (Amersham Protron, GE Healthcare Life Science) or 0.45µm PVDF (Immobilon^®^-P, Millipore) membranes. Blots were blocked either with 5% dried milk or 3% BSA in TNT (15 mM Tris, 140 mM NaCl, 0.05% Tween 20, pH 8). The rat anti-HA 3F10 (1:1000) (Roche; 11867460001), mouse anti-ROP5 T53E2 (1:500) (El Hajj et al., 2007), rabbit anti-FLAG (1:5000) (Sigma, F7425) and mouse anti-TY hybridoma (1:200) (Bastin et al., 1996) were diluted in blocking solution. FLAG-tagged proteins were visualized with Horseradish Peroxidase (HRP)-conjugated donkey anti-rabbit (1:10000) (Jackson Immuno Research), HA-tagged proteins with anti-rat alkaline-phosphatase (AP)-conjugated (1:10000) (Invitrogen), TY-tagged proteins and TgROP5 were visualized with anti-mouse alkaline-phosphatase (AP)-conjugated (1:7500) (Promega) secondary antibodies. Proteins were visualized with BCIP/NBT Color development (Promega) or Clarity Max^TM^ Western ECL (Bio-Rad) substrates. ECL-based detection was performed with Chemidoc system (Bio-Rad).

### Immunoprecipitation and co-immunoprecipitation of *Toxoplasma* proteins

Immunoprecipitation (IP) of TgCRMPa-HA_3_ and TgCRMPb-HA_3_, and co-immunoprecipitation (co-IP) of TgCRMPa-FLAG_3_ and TgCRMPb-HA_3_, were performed from 500×10^6^ tachyzoites, resuspended in 1ml cold lysis buffer (1% NP40, 50mM Tris pH7.4, 150mM NaCl, 4mM EDTA) supplemented with protease inhibitor cocktail tablets (Roche; 11867460001) and gently mixed for 4h at 4°C. Lysates were cleared by centrifugation at 13500g for 30 min at 4°C, and supernatants were transferred in a 1.5ml tube for overnight incubation with proper antibody-conjugated magnetic beads. IP supernatants for mass-spectrometry analysis were incubated with 50µl anti-HA beads (Pierce; R88836) while, those for co-IP, were split in two tubes and separately incubated with 50µl anti-HA beads (Pierce; R88836) and 50µl anti-FLAG M2 beads (Sigma; M8823). The beads were then washed five times with lysis buffer and resuspended in 100°C Laemmli SDS or lithium dodecyl sulfate (LDS) sample buffer, supplemented with 10-40mM dithiothreitol (DTT) (Sigma). Untagged parasites were treated in parallel. Prior to mass spectrometry analysis, protein samples were loaded on a 3-8% gel for SDS-PAGE and stained with Coomassie Blue R-250 solution, to verify protein enrichment upon immunoisolation of the protein of interest. Protein samples from co-IP experiments were resolved with the Novex NuPAGE Gel system (3-8% Tris-Acetate gels, Invitrogen) and subjected to western blotting as described earlier. TgCRMPa-FLAG_3_ and TgCRMPb-HA_3_ proteins were co-stained with the rat anti-HA 3F10 (1:1000) (Roche; 11867460001) and rabbit anti-FLAG (1:5000) (Sigma; F7425) primary antibodies, in combination with anti-rabbit alkaline-phosphatase (AP)-conjugated (1:7500) (Promega) and Horseradish Peroxidase (HRP)-conjugated donkey anti-rat (1:10000) (Jackson Immuno Research) secondary antibodies, respectively. TgROP5 was used as negative control for co-IP experiments, and searched in the clear lysates (before incubation with beads) and IP eluates with mouse anti-ROP5 T53E2 (1:500) (El Hajj et al., 2007) antibodies, followed by anti-mouse alkaline-phosphatase (AP)-conjugated (1:7500) (Promega) secondary antibodies. Proteins were visualized with BCIP/NBT Color development (Promega) or Clarity Max^TM^ Western ECL (Bio-Rad) substrates. ECL-based detection was performed with Chemidoc system (Bio-Rad).

### Mass spectrometry analysis of *Toxoplasma* proteins

Proteins were digested in gel (2 bands per sample) as previously described (Skorupa et al., 2013). Peptides were loaded onto a 25 cm reversed phase column (75 mm inner diameter, Acclaim Pepmap 100® C18, Thermo Fisher Scientific) and separated with an Ultimate 3000 RSLC system (Thermo Fisher Scientific) coupled to a Q Exactive HFX (Thermo Fisher Scientific). MS/MS analyses were performed in a data-dependent mode. Full scans (375 – 1,500 m/z) were acquired in the Orbitrap mass analyzer with a resolution of 60,000 at 200 m/z. For the full scans, 3e6 ions were accumulated within a maximum injection time of 60 ms. The twelve most intense ions with charge states ≥ 2 were sequentially isolated (1e5) with a maximum injection time of 45 ms and fragmented by HCD (Higher-energy collisional dissociation) in the collision cell (normalized collision energy of 28) and detected in the Orbitrap analyzer at a resolution of 30,000. Raw spectra were processed using the MaxQuant (Cox and Mann, 2008) using standard parameters with match between runs (Cox et al., 2011). MS/MS spectra were matched against the UniProt Reference proteomes of *Toxoplasma gondii* and Human (respectively Proteome ID UP000001529, v2019_11 and UP000005640, v2020_01) and 250 frequently observed contaminants as well as reversed sequences of all entries (MaxQuant contaminant database). Statistical analysis were done using Perseus on intensity data (Tyanova et al., 2016).

### Freeze-fracture and transmission electron microscopy of *Toxoplasma* and *Tetrahymena* strains

*Tetrahymena ΔCRMP1* cells were grown overnight in 30 ml SPP to mid log phase density, harvested by centrifugation at 1800g for 3 min and washed once with 10mM Tris pH 7.4. Cells were resuspended in 5ml 20mM phosphate buffer (prepared from 0.1M stock pH 7.1, containing 0.1M sodium phosphate monobasic and 0.1M sodium phosphate dibasic at 1:4 ratio) and fixed by adding 5ml 3% glutaraldehyde (1.5% final conc.) diluted in 30mM phosphate buffer, for 4h at room temperature with gentle mixing. 3ml of fixed *Tetrahymena* cells were withdrawn from the total amount, pelleted to remove the fixative solution and resuspended in 30mM phosphate buffer. *Toxoplasma* TgCRMPa_iKD parasites were cultured for 48h in presence of ATc; freshly egressed tachyzoites were harvested by centrifugation at 650g for 5 min, and fixed as solid pellets with 2.5% glutaraldehyde in 0.1M phosphate buffer for 2h at room temperature. Upon removal of the fixative, pellets were maintained in 30% glycerol diluted in 0.1M phosphate buffer. Fixed *Tetrahymena* and *Toxoplasma* samples were subjected to freeze fracture as previously described (Aquilini et al., 2021). Briefly, cells were quickly frozen in liquid nitrogen and fractured in a Bal-Tec BAF060 apparatus. The fracture surface replica was obtained by evaporating platinum at a 45°angle (∼3.2nm thick) and carbon at a 90° angle (∼25nm thick) respectively. Replicas were washed in 6.5% sodium hypochlorite, rinsed first in chloroform solution (2:1, v/v) (this step was skipped for *Tetrahymena* replicas) and then in distilled water prior to mounting on copper grids. Images were acquired with a Jeol 1200 EXII transmission electron microscope FLAGat the Electron Microscopy Platform of the University of Montpellier, adjusted in brightness and contrast, with the program Fiji (Schindelin et al., 2012).

### Tachyzoites preparation for Cryo-ET of *Toxoplasma*

RH strain *T. gondii* tachyzoites (both wildtype and CRMPb_iKD mutant) were cultivated as described earlier (Suarez et al., 2019) with minor modifications. Tachyzoites were grown within monolayer human foreskin fibroblasts (HFF – ATCC, CRL 1634) in culture media composed of DMEM-10 (ThermoFisher, Cat# 10313039) supplemented with 5% fetal calf serum, 2 mM glutamine and a cocktail of Penicillin–Streptomycin. Extracellular parasites freshly egressed were isolated and concentrated in culture media before freezing 4 µl of this suspension (∼4 × 10^6^ tachyzoites) in each EM grid. For the purpose of tomogram reconstruction, the cell suspension was premixed with 10 nm colloidal gold fiducials (Ted Pella, Cat# 15703) prior to freezing. The CRMPb_iKD parasites were pre-treated with 1 µM ATc (Sigma-Aldrich, Cat# 37919) for 48 h before freezing.

### Cryo-electron tomography (cryo-ET) and subtomogram averaging

Cryo-ET and subtomogram averaging were performed as previously described (Mageswaran et al., 2021). Briefly, projection images were recorded on a Thermo Fisher Titan Krios G3i 300 keV field emission gun cryogenic electron microscope equipped with a K3 direct electron detector (Gatan Inc., Pleasanton, CA, USA) using SerialEM software (Mastronarde, 2005). The camera was operated in the electron-counted mode and images were dose-fractionated at 10 frames per second. Images were motion corrected using the AlignFrames function in IMOD software package (Kremer et al., 1996). Volta phase plate (Danev et al., 2014; Fukuda et al., 2015) and Gatan Imaging Filter (Gatan Inc., Pleasanton, CA, USA;(Krivanek et al., 1995)) with a slit width of 20 eV were used to increase contrast. The imaging workflow is as follows: cells were initially assessed at lower magnifications for ice thickness and plasma membrane integrity, following which tilt series were collected with a span of 120° (−60° to +60°; bi-directional or dose-symmetric scheme) with 2° increments accounting for a total dosage of 120-140 e−/Å^2^ per tilt series. Tilt series were collected at 33,000X magnification with a corresponding pixel size of 2.65 Å (it is noteworthy that a part of the CRMPb_iKD dataset was collected on a replacement K3 camera that reported a slightly increased pixel size of 2.72 Å). Each tilt series had a fixed defocus value between 1 and 3 µm under focus. Our in-house automated computation pipeline (built on functions from the IMOD software package) was used to align tilt series and reconstruct tomograms; the 10 nm colloidal gold served as fiducials for the alignment procedure. IMOD’s slicer program was used to visualize tomograms. After orienting the 3-D volume and sectioning through the desired location, we generally averaged a few slices above and below to enhance contrast. Subtomogram averaging was performed on the RSA (including the AV) of *CRMPb_iKD* cells using PEET (Nicastro et al., 2006). These features were first located in several parasite tomograms by manually inspecting their apical region and subtomograms (a.k.a. particles) were extracted from each in a manually pre-oriented fashion. Subsequently, these particles were computationally aligned over four iterations, each one using reduced angular and translational search parameters compared to the previous. A template was generated for the first alignment iteration by computing an initial average from the manually pre-oriented particles; the template for each of the subsequent iterations was computed by averaging the aligned particles from the corresponding previous iteration. An alignment mask encompassing the RSA (along with the plasma membrane) and the anterior region of the AV was used. After noticing a conserved 8-fold rotational symmetry for the RSA along the longitudinal axis (one that is roughly perpendicular to the patch of plasma membrane where the RSA is anchored), we generated 8-fold more particles by iteratively rotating each particle and repeated our alignment and averaging procedure. We thus enhanced the signal-to-noise ratio, which allowed us to resolve the finer details of the RSA ultrastructure in CRMPb_iKD. In total, we used 22 unique particles, which contributed 176 particles while exploiting the 8-fold symmetry, to generate the final average.

### Quantifications of Cryo-ET data, statistics and reproducibility

We obtained a total of 100 tomograms (over 7 days spanning 2 independent imaging sessions) for wild-type *T. gondii* and 59 tomograms (over 7 days spanning 3 independent imaging sessions) for CRMPb_iKD *T. gondii*, each dataset from multiple frozen grids. The wild-type dataset is the same as the one previously published (Mageswaran et al., 2021). Each of the quantifications (described below) was performed on a randomly chosen subset of these tomograms that resolved the feature of interest. In the case of the wildtype, quantifications were performed again independent of the previous quantifications in Mageswaran et al., 2021 to control for small discrepancies in measurements that could arise from different users or from different attempts at the same analysis. Parasites showed some flattening on the EM grid, likely due to blotting. However, this flattening did not reflect on the shape of the AV or the RSA. Flattening could have caused relatively small displacements of these features but their organizational patterns in the wildtype and CRMPb_iKD cells were evident despite the presence of such potential noise.

#### AV measurements

AV_dist_ (AV anchoring distance): the shortest distance measured from the parasite apex to the AV membrane. The apex is defined as the central position on the PPM where the RSA is anchored.

Ψ’ (a measure of AV offset under the RSA): the angle formed between the orthogonal from the apex and the line connecting the AV centroid to the apex.

AV dimensions: Each AV was described by approximating it to a 2-D ellipse using only 2 axes for simplicity (instead of describing in 3-D using 3 axes). The longest axis for each vesicle in 3-D was marked as the major axis (labeled as AV_maj_) while the shortest axis orthogonal to the major axis and intersecting it at the centroid was marked the minor axis (labeled as AV_min_). In other words, one of the central slices of the AV (representing an ellipse approximation) was used to describe the vesicle. Eccentricity (or Ecc) is calculated as (1-b2/a2)1/2, where ‘a’ is the semi-major axis, and ‘b’ is the semi-minor axis.

Sample size: 22 tomograms each of wildtype and CRMPb_iKD cells were used for the above quantifications except for Ψ’, which used 37 tomograms for each sample.

Quantifications were performed using models generated in IMOD slicer. Models were exported as text files, parsed and analyzed using Python 3.8 or 3.9. Numpy, Matplotlib and Seaborn libraries were used for plotting. Boxplots show the distribution of measurements; in each dataset, the lower and upper boundaries of the box represent the 1^st^ and 3^rd^ quartiles (Q1 and Q3), whiskers extend to 1.5 times the interquartile range (Q3-Q1) below and above Q1 and Q3, and points outside (diamonds) are regarded as outliers (NOTE: the whiskers on either side are shortened if there no data points spanning the previously calculated whisker length). The horizontal divider within the box represents the median. The boxplots are overlaid with swarmplots, each data point representing a measurement from a tomogram. Jointplots are a combination of a bivariate scatterplot and two marginal univariate kernel density estimate plots (a.k.a. probability density plots). Mann–Whitney U test, which is a non-parametric alternative for unpaired Student’s t-test (available within Python’s Scipy package), was used to calculate the p-values. Actual p-values are presented in the plots for values <0.1. For values greater than 0.1, they are replaced with n.s (not significant).

### *Toxoplasma* conoid extrusion assay

To induce conoid extrusion in TgCRMPa-HA_3_ and TgCRMPb-HA_3_ lines, 150-300µl of freshly egressed parasites were added to poly-D-lysine-coated coverslips preheated at 37°C in a 24-well plate. The plate was centrifuged at 400 g for 1min to attach the parasites to the coverslips, and the medium was carefully removed. 300µl of preheated HEPES buffer (274mM NaCl, 10mM KCl, 2mM Na_2_HPO_4_, 11mM glucose, 42mM HEPES, pH 7.05) supplemented with 5mM CaCl_2_ and 5µM A23187 ionophore, were added to each coverslip. HEPES buffer without A23187 was added to control coverslips where, spontaneous conoid extrusion may occur. The plate was incubated at 37°C for 8 min to stimulate A23187-dependent conoid extrusion.

Parasites were fixed with 4% PFA/PBS for 30 min at room temperature upon removal of the buffer, quenched with 100mM glycine/PBS for 10 min, washed with PBS, permeabilized with 0.1% Triton X-100/PBS, and incubated with 10%FBS/PBS blocking solution for 1h. Parasites staining was performed with rat anti-HA 3F10 (1:1000) (Roche; 11867460001) and AlexaFluor488-conjugated goat anti-rat (1:2000) (Invitrogen) antibodies. DNA was stained with 16µM Hoechst, and coverslips were mounted onto microscope slides using Immunomount (Calbiochem). Imaging was performed with a Leica Thunder microscope, with a 100x oil objective NA=1.4, equipped with the sCMOS 4.2MP camera, using Leica Application Suite X (LAS X) software (Leica Biosystems). Z-stacks were denoised, adjusted in brightness and contrast, and colored with the program Fiji (Schindelin et al., 2012).

### Ultrastructure expansion microscopy (U-ExM) of *Toxoplasma* tachyzoites

This technique allows a near-native expansion of cell structures, enabling parasites to stretch up to four times their initial size. Freshly egressed tachyzoites, co-expressing TgCRMPa-HA_3_ and TgCRMPb-HA_3_ with TgNd6-TY_2_, were treated with A23187 as described earlier, to induce extrusion of the conoid. The untagged line was processed in parallel. Upon fixation with 4%PFA/PBS and quenching with 100mM glycine/PBS, PBS-washed coverslips were transferred to a 12-well plate and protein cross-linking was allowed for 5h at 37°C with 1.4% formaldehyde and 2% acrylamide diluted in PBS. Parasites were then embedded in a gel made of a monomer solution (19% sodium acrylate, 10% acrylamide, 0.1% N, N’-methylenebisacrylamide in PBS) supplemented with 10% TEMED and 10% APS; gelation proceeded for 1h at 37°C. Gels containing the parasites were detached from coverslips while dipped in a denaturation buffer (200mM SDS, 200mM NaCl, 50mM Tris, pH 9), and heated at 70°C for 90 min to denature proteins. Gels were expanded in ddH_2_O overnight at room temperature, and then shrank in PBS for antibody incubation. Staining of TgCRMPa-HA_3_ and TgCRMPb-HA_3_ with TgNn6-TY_2_ was performed with rabbit anti-HA (1:2500) (Abcam; ab9110) and mouse anti-TY hybridoma (1:50) (Bastin et al., 1996) primary antibodies, together with guinea pig anti-αtubulin (1:200) (AA345; University of Geneva) and guinea pig anti-βtubulin (1:200) (AA344; University of Geneva) antibodies, used to label subpellicular microtubules. Secondary antibody staining was performed with AlexaFluor647 goat anti-rabbit (1:1000), AlexaFluor594 goat anti-mouse (1:2000), AlexaFluor488 goat anti-guinea pig HCA (1:1500) secondary antibodies (Invitrogen). Gels were incubated with primary and secondary antibodies for 3h at 37°C each time. Antibody were diluted in 2% BSA/PBS and washes after each antibody incubation were performed with PBS containing 0.1% Tween20. Gels were subjected to a second round of expansion in ddH_2_O overnight prior to microscopy imaging. Expanded parasites were imaged with Zeiss LSM880 confocal microscope equipped with Airyscan detector, with a 63x oil objective NA=1.4, using Zen Black software (Zeiss, Intelligent Imaging Innovations). Z-stacks were denoised, adjusted in brightness and contrast, colored and processed to obtain maximum intensity projections, with the program Fiji (Schindelin et al., 2012).

## REFERENCES

Achbarou, A., Mercereau-Puijalon, O., Sadak, A., Fortier, B., Leriche, M.A., Camus, D., and Dubremetz, J.F. (1991a). Differential targeting of dense granule proteins in the parasitophorous vacuole of Toxoplasma gondii. Parasitology 3, 321–329.

Achbarou, A., Mercereau-Puijalon, O., Autheman, J.M., Fortier, B., Camus, D., and Dubremetz, J.F. (1991b). Characterization of microneme proteins of Toxoplasma gondii. Mol Biochem Parasitol 47, 223–233.

Adams, J.C., and Tucker, R.P. (2000). The thrombospondin type 1 repeat (TSR) superfamily: diverse proteins with related roles in neuronal development. Dev Dyn 218, 280–299.

Aquilini, E., Cova, M.M., Mageswaran, S.K., Dos Santos Pacheco, N., Sparvoli, D., Penarete-Vargas, D.M., Najm, R., Graindorge, A., Suarez, C., Maynadier, M., et al. (2021). An Alveolata secretory machinery adapted to parasite host cell invasion. Nat Microbiol 6, 425–434.

Barylyuk, K., Koreny, L., Ke, H., Butterworth, S., Crook, O.M., Lassadi, I., Gupta, V., Tromer, E., Mourier, T., Stevens, T.J., et al. (2020). A Comprehensive Subcellular Atlas of the Toxoplasma Proteome via hyperLOPIT Provides Spatial Context for Protein Functions. Cell Host Microbe 28, 752–766.e9.

Bastin, P., Bagherzadeh, Z., Matthews, K.R., and Gull, K. (1996). A novel epitope tag system to study protein targeting and organelle biogenesis in Trypanosoma brucei. Mol Biochem Parasitol 77, 235–239.

Beisson, J., Lefort-Tran, M., Pouphile, M., Rossignol, M., and Satir, B. (1976). Genetic analysis of membrane differentiation in Paramecium. Freeze-fracture study of the trichocyst cycle in wild-type and mutant strains. The Journal of Cell Biology 69, 126–143.

Besteiro, S., Michelin, A., Poncet, J., Dubremetz, J.-F., and Lebrun, M. (2009). Export of a Toxoplasma gondii Rhoptry Neck Protein Complex at the Host Cell Membrane to Form the Moving Junction during Invasion. PLOS Pathogens 5, e1000309.

Besteiro, S., Dubremetz, J.F., and Lebrun, M. (2011). The moving junction of apicomplexan parasites: a key structure for invasion. Cell Microbiol 13, 797–805.

Blumenschein, T.M., Friedrich, N., Childs, R.A., Saouros, S., Carpenter, E.P., Campanero-Rhodes, M.A., Simpson, P., Chai, W., Koutroukides, T., Blackman, M.J., et al. (2007). Atomic resolution insight into host cell recognition by Toxoplasma gondii. Embo J 26, 2808–2820.

Bolte, S., and Cordelieres, F.P. (2006). A guided tour into subcellular colocalization analysis in light microscopy. J Microsc 224, 213–232.

Brown, K.M., Long, S., and Sibley, L.D. (2017). Plasma Membrane Association by N-Acylation Governs PKG Function in Toxoplasma gondii. MBio 8, e00375–17.

Brown, K.M., Long, S., and Sibley, L.D. (2018). Conditional Knockdown of Proteins Using Auxin-inducible Degron (AID) Fusions in Toxoplasma gondii. Bio Protoc 8, e2728.

Burrell, A., Marugan-Hernandez, V., Moreira-Leite F, Djp, F., Tomley, F, and Vaughan, S (2021). Cellular electron tomography of the apical complex 1 in the apicomplexan 2 parasite Eimeria tenella shows a highly organised gateway for regulated secretion.

Carruthers, V.B., and Sibley, L.D. (1997). Sequential protein secretion from three distinct organelles of Toxoplasma gondii accompanies invasion of human fibroblasts. Eur J Cell Biol 73, 114–123.

Carruthers, V.B., and Sibley, L.D. (1999). Mobilization of intracellular calcium stimulates microneme discharge in Toxoplasma gondii. Mol Microbiol 31, 421–428.

Cassidy-Hanley, D., Bowen, J., Lee, J.H., Cole, E., VerPlank, L.A., Gaertig, J., Gorovsky, M.A., and Bruns, P.J. (1997). Germline and somatic transformation of mating Tetrahymena thermophila by particle bombardment. Genetics 146, 135–147.

Cerede, O., Dubremetz, J.F., Soete, M., Deslee, D., Vial, H., Bout, D., and Lebrun, M. (2005). Synergistic role of micronemal proteins in Toxoplasma gondii virulence. J Exp Med 201, 453–463.

Chilcoat, N.D., Melia, S.M., Haddad, A., and Turkewitz, A.P. (1996). Granule lattice protein 1 (Grl1p), an acidic, calcium-binding protein in Tetrahymena thermophila dense-core secretory granules, influences granule size, shape, content organization, and release but not protein sorting or condensation. J Cell Biol 135, 1775–1787.

Coleman, B.I., Saha, S., Sato, S., Engelberg, K., Ferguson, D.J.P., Coppens, I., Lodoen, M.B., and Gubbels, M.J. (2018). A Member of the Ferlin Calcium Sensor Family Is Essential for Toxoplasma gondii Rhoptry Secretion. MBio 9.

Couvreur, G., Sadak, A., Fortier, B., and Dubremetz, J.F. (1988). Surface antigens of Toxoplasma gondii. Parasitology 97 (Pt 1), 1–10.

Cowan, A.T., Bowman, G.R., Edwards, K.F., Emerson, J.J., and Turkewitz, A.P. (2005). Genetic, genomic, and functional analysis of the granule lattice proteins in Tetrahymena secretory granules. Mol. Biol. Cell 16, 4046–4060.

Cox, J., and Mann, M. (2008). MaxQuant enables high peptide identification rates, individualized p.p.b.-range mass accuracies and proteome-wide protein quantification. Nature Biotechnology 26, 1367–1372.

Cox, J., Neuhauser, N., Michalski, A., Scheltema, R.A., Olsen, J.V., and Mann, M. (2011). Andromeda: a peptide search engine integrated into the MaxQuant environment. Journal of Proteome Research 10, 1794–1805.

Danev, R., Buijsse, B., Khoshouei, M., Plitzko, J.M., and Baumeister, W. (2014). Volta potential phase plate for in-focus phase contrast transmission electron microscopy. Proc Natl Acad Sci U S A 111, 15635–15640.

Douradinha, B., Augustijn, K.D., Moore, S.G., Ramesar, J., Mota, M.M., Waters, A.P., Janse, C.J., and Thompson, J. (2011). Plasmodium Cysteine Repeat Modular Proteins 3 and 4 are essential for malaria parasite transmission from the mosquito to the host. Malar J 10, 71.

Dubremetz, J.F. (1998). Host cell invasion by Toxoplasma gondii. Trends Microbiol. 6, 27–30.

El Hajj, H., Lebrun, M., Fourmaux, M.N., Vial, H., and Dubremetz, J.F. (2007). Inverted topology of the Toxoplasma gondii ROP5 rhoptry protein provides new insights into the association of the ROP2 protein family with the parasitophorous vacuole membrane. Cell Microbiol 9, 54–64.

El Hajj, H., Papoin, J., Cérède, O., Garcia-Réguet, N., Soête, M., Dubremetz, J.-F., and Lebrun, M. (2008). Molecular Signals in the Trafficking of Toxoplasma gondii Protein MIC3 to the Micronemes. Eukaryot Cell 7, 1019–1028.

Fredriksson, R., and Schiöth, H.B. (2005). The repertoire of G-protein-coupled receptors in fully sequenced genomes. Mol Pharmacol 67, 1414–1425.

Frenal, K., Marq, J.B., Jacot, D., Polonais, V., and Soldati-Favre, D. (2014). Plasticity between MyoC- and MyoA-glideosomes: an example of functional compensation in Toxoplasma gondii invasion. PLoS Pathogens 10, e1004504.

Friedrich, N., Santos, J.M., Liu, Y., Palma, A.S., Leon, E., Saouros, S., Kiso, M., Blackman, M.J., Matthews, S., Feizi, T., et al. (2010). Members of a novel protein family containing microneme adhesive repeat domains act as sialic acid-binding lectins during host cell invasion by apicomplexan parasites. J Biol Chem 285, 2064–2076.

Froissard, M., Keller, A.M., and Cohen, J. (2001). ND9P, a novel protein with armadillo-like repeats involved in exocytosis: physiological studies using allelic mutants in paramecium. Genetics 157, 611–620.

Fukuda, Y., Laugks, U., Lučić, V., Baumeister, W., and Danev, R. (2015). Electron cryotomography of vitrified cells with a Volta phase plate. J Struct Biol 190, 143–154.

Ghosh, S., Kennedy, K., Sanders, P., Matthews, K., Ralph, S.A., Counihan, N.A., and de Koning-Ward, T.F. (2017). The Plasmodium rhoptry associated protein complex is important for parasitophorous vacuole membrane structure and intraerythrocytic parasite growth. Cellular Microbiology 19.

Gilbert, L.A., Ravindran, S., Turetzky, J.M., Boothroyd, J.C., and Bradley, P.J. (2007). Toxoplasma gondii targets a protein phosphatase 2C to the nuclei of infected host cells. Eukaryot Cell 6, 73–83.

Gogendeau, D., Keller, A.M., Yanagi, A., Cohen, J., and Koll, F. (2005). Nd6p, a novel protein with RCC1-like domains involved in exocytosis in Paramecium tetraurelia. Eukaryotic Cell 4, 2129–2139.

Guerin, A., Corrales, R.M., Parker, M.L., Lamarque, M.H., Jacot, D., El Hajj, H., Soldati-Favre, D., Boulanger, M.J., and Lebrun, M. (2017). Efficient invasion by Toxoplasma depends on the subversion of host protein networks. Nature Microbiology 2, 1358–1366.

Haddad, A., and Turkewitz, A.P. (1997). Analysis of exocytosis mutants indicates close coupling between regulated secretion and transcription activation in Tetrahymena. Proceedings of the National Academy of Sciences of the United States of America 94, 10675–10680.

Hakimi, M.-A., Olias, P., and Sibley, L.D. (2017). Toxoplasma Effectors Targeting Host Signaling and Transcription. Clin. Microbiol. Rev. 30, 615–645.

Hanssen, E., Dekiwadia, C., Riglar, D.T., Rug, M., Lemgruber, L., Cowman, A.F., Cyrklaff, M., Kudryashev, M., Frischknecht, F., Baum, J., et al. (2013). Electron tomography of Plasmodium falciparum merozoites reveals core cellular events that underpin erythrocyte invasion. Cell Microbiol 15, 1457–1472.

Hauser, A.S., Chavali, S., Masuho, I., Jahn, L.J., Martemyanov, K.A., Gloriam, D.E., and Babu, M.M. (2018). Pharmacogenomics of GPCR Drug Targets. Cell 172, 41–54.e19.

Huynh, M.-H., and Carruthers, V.B. (2009). Tagging of endogenous genes in a Toxoplasma gondii strain lacking Ku80. Eukaryotic Cell 8, 530–539.

Huynh, M.-H., and Carruthers, V.B. (2016). A Toxoplasma gondii Ortholog of Plasmodium GAMA Contributes to Parasite Attachment and Cell Invasion. MSphere 1, e00012–16.

Ishino, T., Murata, E., Tokunaga, N., Baba, M., Tachibana, M., Thongkukiatkul, A., Tsuboi, T., and Torii, M. (2019). Rhoptry neck protein 2 expressed in Plasmodium sporozoites plays a crucial role during invasion of mosquito salivary glands. Cell Microbiol 21, e12964.

Kemp, L.E., Yamamoto, M., and Soldati-Favre, D. (2012). Subversion of host cellular functions by the apicomplexan parasites. FEMS Microbiol Rev.

Kessler, H., Herm-Gotz, A., Hegge, S., Rauch, M., Soldati-Favre, D., Frischknecht, F., and Meissner, M. (2008). Microneme protein 8--a new essential invasion factor in Toxoplasma gondii. J Cell Sci 121, 947–956.

Kim, K., Soldati, D., and Boothroyd, J.C. (1993). Gene replacement in Toxoplasma gondii with chloramphenicol acetyltransferase as selectable marker. Science 262, 911–914.

Koshy, A.A., Fouts, A.E., Lodoen, M.B., Alkan, O., Blau, H.M., and Boothroyd, J.C. (2010). Toxoplasma secreting Cre recombinase for analysis of host-parasite interactions. Nat Methods 7, 307–309.

Kremer, J.R., Mastronarde, D.N., and McIntosh, J.R. (1996). Computer visualization of three-dimensional image data using IMOD. Journal of Structural Biology 116, 71–76.

Kremer, K., Kamin, D., Rittweger, E., Wilkes, J., Flammer, H., Mahler, S., Heng, J., Tonkin, C.J., Langsley, G., Hell, S.W., et al. (2013). An overexpression screen of Toxoplasma gondii Rab-GTPases reveals distinct transport routes to the micronemes. PLoS Pathog 9, e1003213.

Krivanek, O.L., Friedman, S.L., Gubbens, A.J., and Kraus, B. (1995). An imaging filter for biological applications. Ultramicroscopy 59, 267–282.

Kumar, S., Briguglio, J.S., and Turkewitz, A.P. (2014). An aspartyl cathepsin, CTH3, is essential for proprotein processing during secretory granule maturation in Tetrahymena thermophila. Molecular Biology of the Cell 25, 2444–2460.

Kumar, S., Stecher, G., Li, M., Knyaz, C., and Tamura, K. (2018). MEGA X: Molecular Evolutionary Genetics Analysis across Computing Platforms. Mol Biol Evol 35, 1547–1549.

Lamarque, M.H., Roques, M., Kong-Hap, M., Tonkin, M.L., Rugarabamu, G., Marq, J.B., Penarete-Vargas, D.M., Boulanger, M.J., Soldati-Favre, D., and Lebrun, M. (2014). Plasticity and redundancy among AMA-RON pairs ensure host cell entry of Toxoplasma parasites. Nature Communications 5, 4098.

Lebrun, M, Carrthers, V., and Cesbron-Delauw, MF (2020). The Model Apicomplexan -Perspectives and Methods. Chapter 14 - Toxoplasma secretory proteins and their roles in parasite cell cycle and infection. 607–704.

Liebscher, I., Cevheroğlu, O., Hsiao, C.-C., Maia, A.F., Schihada, H., Scholz, N., Soave, M., Spiess, K., Trajković, K., Kosloff, M., et al. (2021). A guide to adhesion GPCR research. FEBS J.

Madeira, L., Galante, P.A.F., Budu, A., Azevedo, M.F., Malnic, B., and Garcia, C.R.S. (2008). Genome-wide detection of serpentine receptor-like proteins in malaria parasites. PLoS One 3, e1889.

Mageswaran, S.K., Guérin, A., Theveny, L.M., Chen, W.D., Martinez, M., Lebrun, M., Striepen, B., and Chang, Y.-W. (2021). In situ ultrastructures of two evolutionarily distant apicomplexan rhoptry secretion systems. Nat Commun 12, 4983.

Mastronarde, D.N. (2005). Automated electron microscope tomography using robust prediction of specimen movements. Journal of Structural Biology 152, 36–51.

Meissner, M., Schluter, D., and Soldati, D. (2002). Role of Toxoplasma gondii myosin A in powering parasite gliding and host cell invasion. Science 298, 837–840.

Mi, H., Ebert, D., Muruganujan, A., Mills, C., Albou, L.-P., Mushayamaha, T., and Thomas, P.D. (2021). PANTHER version 16: a revised family classification, tree-based classification tool, enhancer regions and extensive API. Nucleic Acids Research 49, D394–D403.

Mochizuki, K. (2008). High efficiency transformation of Tetrahymena using a codon-optimized neomycin resistance gene. Gene 425, 79–83.

Mondragon, R., and Frixione, E. (1996). Ca(2+)-dependence of conoid extrusion in Toxoplasma gondii tachyzoites. J Eukaryot Microbiol 43, 120–127.

Mueller, C., Klages, N., Jacot, D., Santos, J.M., Cabrera, A., Gilberger, T.W., Dubremetz, J.-F., and Soldati-Favre, D. (2013). The Toxoplasma protein ARO mediates the apical positioning of rhoptry organelles, a prerequisite for host cell invasion. Cell Host Microbe 13, 289–301.

Nicastro, D., Schwartz, C., Pierson, J., Gaudette, R., Porter, M.E., and McIntosh, J.R. (2006). The molecular architecture of axonemes revealed by cryoelectron tomography. Science 313, 944–948.

Nichols, B.A., Chiappino, M.L., and O’Connor, G.R. (1983). Secretion from the rhoptries of Toxoplasma gondii during host-cell invasion. J Ultrastruct Res 83, 85–98.

Nishimura, K., Fukagawa, T., Takisawa, H., Kakimoto, T., and Kanemaki, M. (2009). An auxin-based degron system for the rapid depletion of proteins in nonplant cells. Nat Methods 6, 917–922.

Patthy, L., Trexler, M., Váli, Z., Bányai, L., and Váradi, A. (1984). Kringles: modules specialized for protein binding. Homology of the gelatin-binding region of fibronectin with the kringle structures of proteases. FEBS Lett 171, 131–136.

Perez-Riverol, Y., Csordas, A., Bai, J., Bernal-Llinares, M., Hewapathirana, S., Kundu, D.J., Inuganti, A., Griss, J., Mayer, G., Eisenacher, M., et al. (2019). The PRIDE database and related tools and resources in 2019: improving support for quantification data. Nucleic Acids Res 47, D442–D450.

Pietrzyk-Brzezinska, A.J., and Bujacz, A. (2020). H-type lectins – Structural characteristics and their applications in diagnostics, analytics and drug delivery. International Journal of Biological Macromolecules 152, 735–747.

Piro, F., Carruthers, V.B., and Di Cristina, M. (2020). PCR Screening of Toxoplasma gondii Single Clones Directly from 96-Well Plates Without DNA Purification. Methods Mol Biol 2071, 117–123.

Plattner, H., Miller, F., and Bachmann, L. (1973). Membrane specializations in the form of regular membrane-to-membrane attachment sites in Paramecium. A correlated freeze-etching and ultrathin-sectioning analysis. Journal of Cell Science 13, 687–719.

Riglar, D.T., Richard, D., Wilson, D.W., Boyle, M.J., Dekiwadia, C., Turnbull, L., Angrisano, F., Marapana, D.S., Rogers, K.L., Whitchurch, C.B., et al. (2011). Super-resolution dissection of coordinated events during malaria parasite invasion of the human erythrocyte. Cell Host Microbe 9, 9–20.

Roiko, M.S., and Carruthers, V.B. (2013). Functional dissection of Toxoplasma gondii perforin-like protein 1 reveals a dual domain mode of membrane binding for cytolysis and parasite egress. The Journal of Biological Chemistry 288, 8712–8725.

Satir, B. (1977). Dibucaine-induced synchronous mucocyst secretion in Tetrahymena. Cell Biol Int Rep 1, 69–73.

Satir, B., Schooley, C., and Satir, P. (1972). Membrane reorganization during secretion in Tetrahymena. Nature 235, 53–54.

Schindelin, J., Arganda-Carreras, I., Frise, E., Kaynig, V., Longair, M., Pietzsch, T., Preibisch, S., Rueden, C., Saalfeld, S., Schmid, B., et al. (2012). Fiji: an open-source platform for biological-image analysis. Nature Methods 9, 676–682.

Schneider, C.A., Rasband, W.S., and Eliceiri, K.W. (2012). NIH Image to ImageJ: 25 years of image analysis. Nat Methods 9, 671–675.

Sheiner, L., Demerly, J.L., Poulsen, N., Beatty, W.L., Lucas, O., Behnke, M.S., White, M.W., and Striepen, B. (2011). A systematic screen to discover and analyze apicoplast proteins identifies a conserved and essential protein import factor. PLoS Pathog 7, e1002392.

Sidik, S.M., Huet, D., Ganesan, S.M., Huynh, M.H., Wang, T., Nasamu, A.S., Thiru, P., Saeij, J.P., Carruthers, V.B., Niles, J.C., et al. (2016). A Genome-wide CRISPR Screen in Toxoplasma Identifies Essential Apicomplexan Genes. Cell 166, 1423–1435 e12.

Singh, S., Alam, M.M., Pal-Bhowmick, I., Brzostowski, J.A., and Chitnis, C.E. (2010). Distinct external signals trigger sequential release of apical organelles during erythrocyte invasion by malaria parasites. PLoS Pathog 6, e1000746.

Skorupa, A., Urbach, S., Vigy, O., King, M.A., Chaumont-Dubel, S., Prehn, J.H., and Marin, P. (2013). Angiogenin induces modifications in the astrocyte secretome: relevance to amyotrophic lateral sclerosis. Journal of Proteomics 91, 274–285.

Suarez, C., Lentini, G., Ramaswamy, R., Maynadier, M., Aquilini, E., Berry-Sterkers, L., Cipriano, M., Chen, A.L., Bradley, P., Striepen, B., et al. (2019). A lipid-binding protein mediates rhoptry discharge and invasion in Plasmodium falciparum and Toxoplasma gondii parasites. Nature Communications 10, 4041.

Suss-Toby, E., Zimmerberg, J., and Ward, G.E. (1996). Toxoplasma invasion: the parasitophorous vacuole is formed from host cell plasma membrane and pinches off via a fission pore. Proc Natl Acad Sci U S A 93, 8413–8418.

Thompson, J., Fernandez-Reyes, D., Sharling, L., Moore, S.G., Eling, W.M., Kyes, S.A., Newbold, C.I., Kafatos, F.C., Janse, C.J., and Waters, A.P. (2007). Plasmodium cysteine repeat modular proteins 1-4: complex proteins with roles throughout the malaria parasite life cycle. Cell Microbiol 9, 1466–1480.

Tsypin, L.M., and Turkewitz, A.P. (2017). The Co-regulation Data Harvester: automating gene annotation starting from a transcriptome database. SoftwareX 6, 165–171.

Turkewitz, A.P., Madeddu, L., and Kelly, R.B. (1991). Maturation of dense core granules in wild type and mutant Tetrahymena thermophila. EMBO J 10, 1979–1987.

Tyanova, S., Temu, T., Sinitcyn, P., Carlson, A., Hein, M.Y., Geiger, T., Mann, M., and Cox, J. (2016). The Perseus computational platform for comprehensive analysis of (prote)omics data. Nature Methods 13, 731–740.

Yona, S., Lin, H.-H., Siu, W.O., Gordon, S., and Stacey, M. (2008). Adhesion-GPCRs: emerging roles for novel receptors. Trends Biochem Sci 33, 491–500.

Zimmermann, L., Stephens, A., Nam, S.-Z., Rau, D., Kübler, J., Lozajic, M., Gabler, F., Söding, J., Lupas, A.N., and Alva, V. (2018). A Completely Reimplemented MPI Bioinformatics Toolkit with a New HHpred Server at its Core. J Mol Biol 430, 2237–2243.

