## Supplementary figures for "An apical membrane complex controls rhoptry exocytosis and invasion in *Toxoplasma*"

Supplementary figure 1

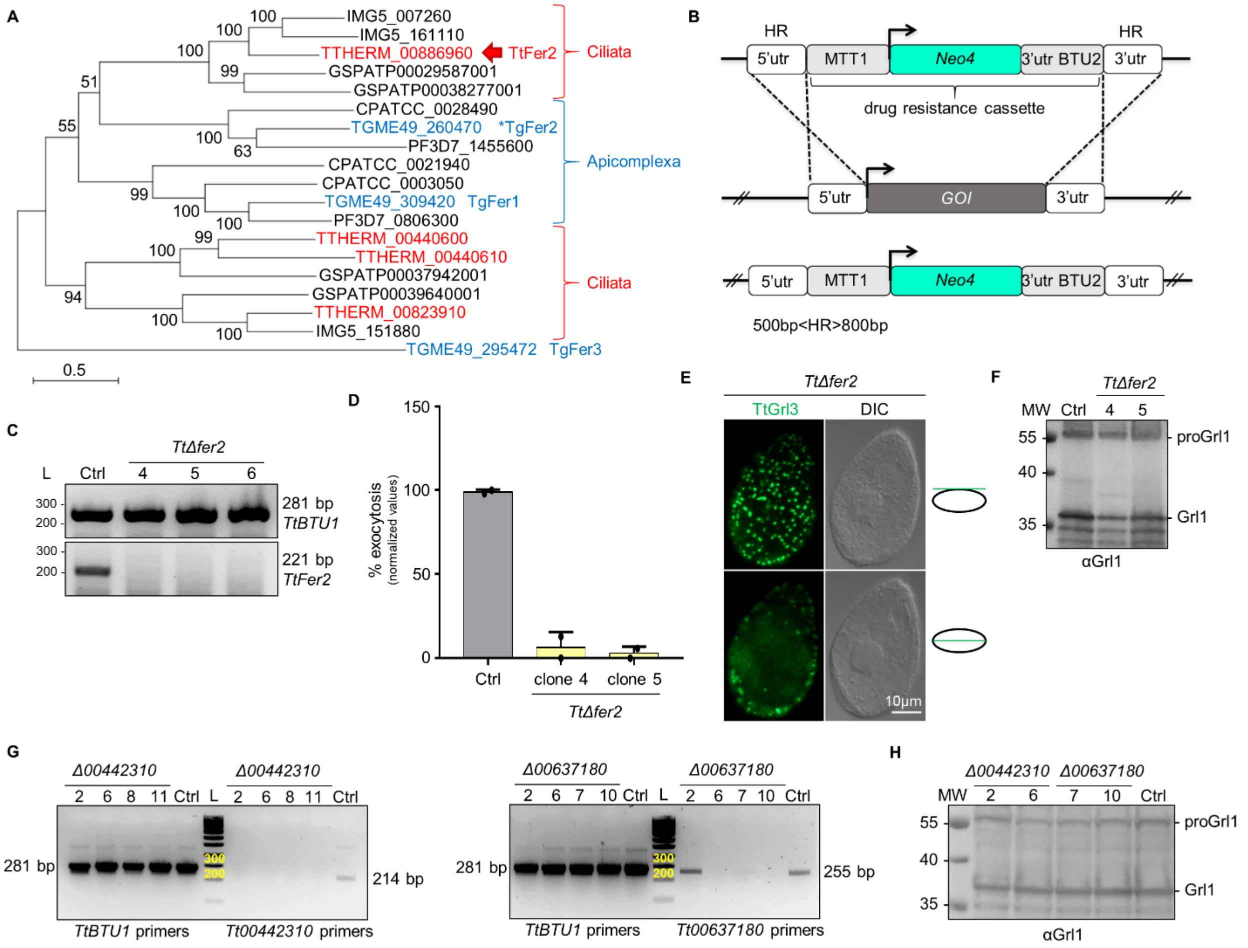


**Figure S1. *Tetrahymena* TtFer2, Tt00442310 and Tt00637180 are essential for mucocyst secretion. Related to figure 1.**

(A) Phylogeny depicting the relationships between Ciliata and Apicomplexa ferlins. The Maximum likelihood phylogenetic tree was obtained with the protein sequences of ferlin genes retrieved for the ciliates *Tetrahymena thermophila* (TTHERM), *Paramecium tetraurelia* (GSPATP) and *Ichthyophthirius multifiliis* (IMG5), and for the apicomplexans *Toxoplasma gondii* (TGME49), *Plasmodium falciparum* (PF3D7) and *Cryptosporidium parvum* (CPATCC). *Tetrahymena* and *Toxoplasma* ferlins are highlighted in red and blue, respectively. The *Tetrahymena* homolog of the rhoptry-related TgFer2 (asterisk) is indicated by the red arrow. The scale bar represents the branch length.

(B) Strategy for the macronuclear knockout of *Tetrahymena thermophila* genes of interest (GOI). A linearized construct carrying fragments (HR) homologous to the 5' and 3'-untranslated regions

(UTR) of the GOI, and flanking the drug resistance cassette, was used to replace the GOI at the endogenous locus. The $CdCl_2$-inducible MTT1 promoter drives the expression of a paromomycin-resistance gene (Neo4) used for selecting positive transformants.

(C) Disruption of the macronuclear copies of *TtFer2* (ferlin 2; TTHERM_00886960) was assessed by RT-PCR. cDNA from wildtype (Ctrl) and three clones of putative knockout cells (*Δfer2)*, was PCR-amplified with primers specific for *TtBTU1* (β-tubulin 1; upper panel), and *TtFer2* (lower panel). The 221bp products corresponding to transcripts from *Fer2* are absent in the *Δfer2* clones, indicating that all the wildtype copies of *TtFer2* were efficiently replaced with the Neo4 cassette. All samples show wildtype levels of *BTU1* transcripts. L: DNA ladder (bp). Primers are listed in Table S5.

(D) Quantification of exocytosis in *Tetrahymena Δfer2* cells stimulated with dibucaine. The values are expressed as percentage of release relatively to the control strain (Ctrl). Mutant cells are severely impaired in mucocyst secretion. (n = 2 biological replicates).

(E) Immunofluorescence images of a *TtΔfer2* cell with paired differential interference contrast (DIC) images. Mucocysts were immunostained with mAbs 5E9 which label the granule protein Grl3, and appeared similar to wildtype (see figure 1E) in shape and docking. Single focal planes of surface (upper) and cross (lower) sections are shown for the same cell.

(F) Western blot of whole-cell lysates from wildtype (Ctrl) and *Δfer2* cells. In both, wildtype and mutant extracts, anti-Grl1 antibodies recognized the ~60kDa precursor of the granule protein 1, proGrl1, and the processed form of Grl1, between 35-40kDa, indicating non-significant defects in proteolytic maturation. MW: molecular weight standards.

(G) Disruption of the macronuclear copies of TTHERM_00442310 and TTHERM_00637180, was assessed by RT-PCR as in C). Four clones for each putative knockout cells were tested. The 214bp and 255bp fragments corresponding to transcripts for TTHERM_00442310 and TTHERM_00637180, respectively, are absent in all *Δ00442310* clones, and nearly undetectable in clones 6, 7 and 10 for *Δ00637180*, indicating the achievement of full knockout. Clones 2 and 6 for *Δ00442310*, and clones 7 and 10 for *Δ00637180*, were selected for further analysis. All samples show wildtype levels of *BTU1* transcripts. L: DNA ladder (bp). Primers are listed in Table S5.

(H) Western blot of whole-cell lysates from wildtype (Ctrl), *Δ00442310* and *Δ00637180* cells. In both, wildtype and mutants extracts, anti-Grl1 antibodies recognized processed Grl1 between 35-

40kDa, and the precursor proGrl1 at ~60kDa, indicating non-significant defects in proteolytic maturation. MW: molecular weight standards.

Supplementary figure 2

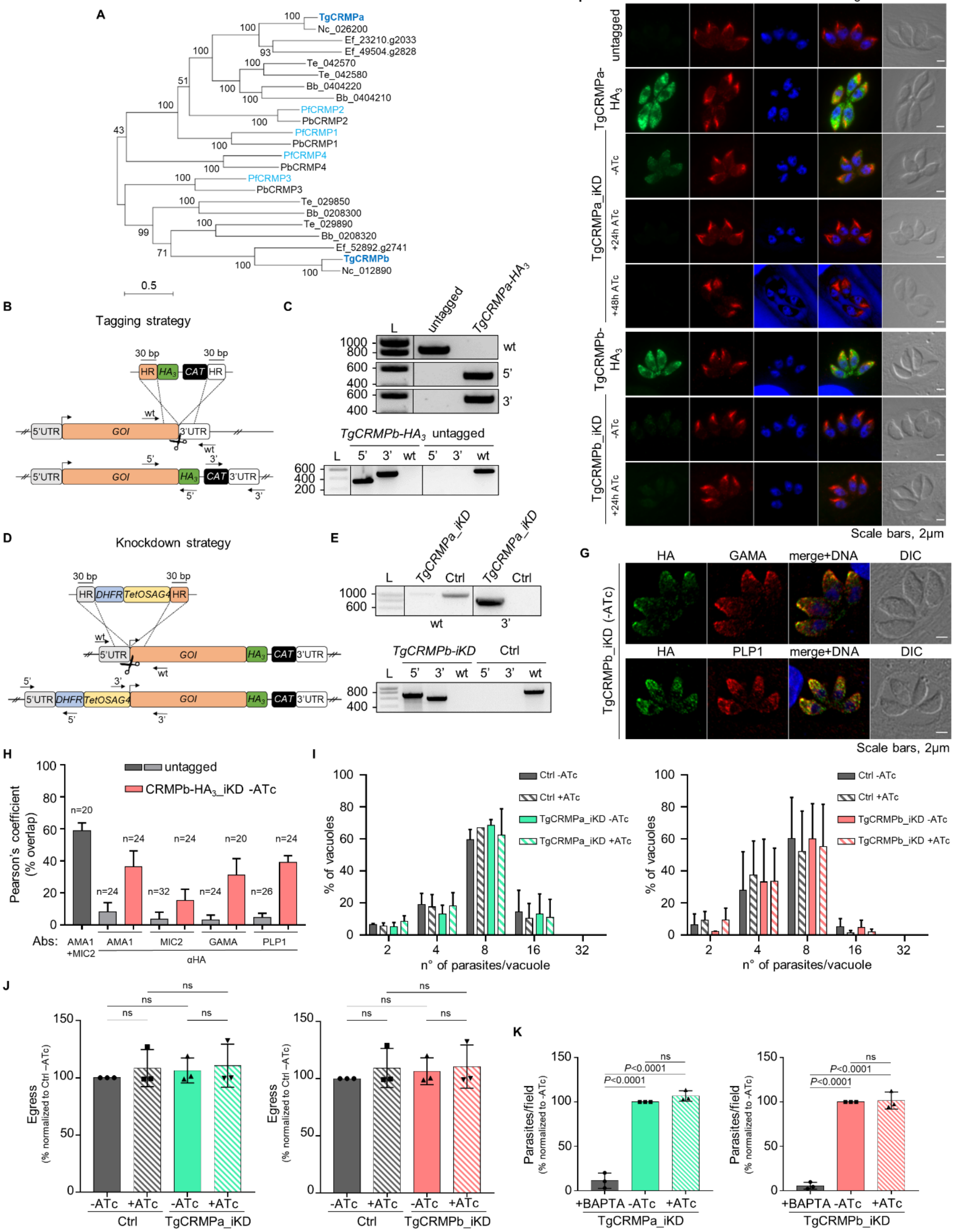

**Figure S2. TgCRMPa- and TgCRMPb-depleted tachyzoites have normal rhoptries, and show no defects in replication, egress and attachment. Related to figure 2.**

(A) Phylogeny depicting the relationships between Apicomplexa CRMPs. The Maximum likelihood phylogenetic tree was obtained with the protein sequences of *CRMP* genes retrieved for the apicomplexans *Toxoplasma gondii* (TgCRMP), *Plasmodium falciparum* (PfCRMP), *Plasmodium berghei* (PbCRMP), *Neospora caninum* (Nc), *Eimeria falciformis* (Ef), *Theileria equi* (Te), and *Babesia bigemina* (Bb). *Toxoplasma* and *P. falciparum* CRMPs are highlighted in bold-blue and light blue, respectively. The scale bar represents the branch length.

(B) Strategy for tagging genes of interest (GOI) in *Toxoplasma*. To generate C-terminal $HA_3$-fusions of TgCRMPa and TgCRMPb, a DNA fragment was amplified from a donor vector containing the $HA_3$ tag and the drug resistance cassette (CAT). Primers to amplify the DNA fragment were designed to contain ~30bp-long stretches (HR) homologous to the GOI regions flanking the insertion site for the epitope tag. Upon CRISPR-cas9 cut (scissors), the PCR-amplified DNA fragment efficiently recombines into the targeted endogenous locus. The arrows indicate the binding sites of the primers used in (C).

(C) Integration of the $HA_3$ tag and CAT cassette at the C-terminus of TgCRMPa (upper panel) and TgCRMPb (lower panel) was tested by PCR. Genomic DNA from the untagged line and a clonal population for each of the putative $HA_3$-tagged lines, was amplified with primers binding to the 3' C-terminus and 3'UTR of each TgCRMP gene, in pairwise combination with primers binding the $HA_3$ and CAT sequences, respectively. The fragments corresponding to the $HA_3$ tag (5') and the resistance cassette (3') were correctly amplified in the putative tagged lines, indicating that they were efficiently integrated at the TgCRMPs loci. As expected, the wildtype fragment of each gene (wt) was detected only in the untagged line. L: DNA ladder (bp). Primers are listed in table S5.

(D) Strategies for the inducible depletion (iKD) of genes of interest (GOI) in *Toxoplasma*. The iKD lines for TgCRMPs were generated starting from the $HA_3$-tagged lines previously produced. In order to conditionally deplete the proteins, the endogenous promoter of each gene was replaced with an ATc-regulatable TetOSag4 promoter, preceded by the DHFR resistance cassette. The DNA fragment containing the cassette and the promoter was PCR-amplified from a donor vector with primers containing ~30bp-long homology regions (HR) specific for each gene, and introduced upstream the starting codon via CRISPR-cas9 technology (scissors) and homologous recombination. The arrows indicate the binding sites of the primers used in (E) and figure S3G.

(E) Integration of the TetOSag4 promoter in TgCRMPa-$HA_3$ (upper panel) and TgCRMPb-$HA_3$ (lower panel) lines was tested by PCR. Integration of the DHFR resistance cassette was successfully PCR-amplified only for TgCRMPb-$HA_3$ (lower panel) line. Genomic DNA from untagged parasites, and putative TgCRMPa_iKD and TgCRMPb_iKD clonal populations, was amplified with primers binding to the 5'UTR and the 5' N-terminus of the GOI, flanking the DHFR-TetOSag4 insert, and used also in pairwise combination with primers binding the DHFR cassette and the TetOSag4 promoter, respectively. The fragments corresponding to the DHFR integration (5') and TetOSag4 integration (3') were detected exclusively in the putative iKD lines, while the wildtype fragment (wt) was amplified only in the untagged line. L: DNA ladder (bp). Primers are listed in table S5.

(F) Immunofluorescence images of untagged, TgCRMPa-$HA_3$ and TgCRMPb-$HA_3$ lines, and TgCRMPs-depleted (iKD) intracellular tachyzoites. Parasites were stained with anti-HA and with anti-ARM (ARO) Abs to visualize TgCRMPs and rhoptries, respectively. The nuclei (DNA) are stained with Hoechst. TgCRMPs pattern mirrors that of figure 2B. Rhoptries show a wildtype appearance in the TgCRMPs-depleted parasites. Shown are single focal planes.

(G) Confocal immunofluorescence images of TgCRMPb-depleted (iKD) intracellular tachyzoites. Parasites were stained with anti-HA and with anti-GAMA and anti-PLP1 Abs to visualize TgCRMPb and micronemes, respectively. The nuclei (DNA) are stained with Hoechst. Shown are single focal planes.

(H) Extent of colocalization between TgCRMPb-$HA_3$ (light red) and microneme proteins AMA1, MIC2, GAMA and PLP1 shown in (G) and figure 2D. Untagged parasites were analyzed in parallel to estimate the background noise (light grey), and the extent of overlap between the microneme proteins AMA1 and MIC2 (dark grey). Pearson's correlation coefficient was measured using the Fiji-JACoP plugin. Values are expressed as mean ± SD; n: number of parasites analyzed.

(I) Replication measured for TgCRMPa- and TgCRMPb-depleted parasites. The percentage of vacuoles with 2, 4, 8, 16, & 32 parasites was calculated for control (Ctrl), TgCRMPa_iKD and TgCRMPb_iKD lines, in absence of ATc and upon 48 and 24h ATc treatment, respectively. Both iKD mutants (+ATc) are capable of efficient replication. Data are reported as mean ± SD (n = 2 biological replicates, each with 3 technical replicates).

(J) Egress was quantified for TgCRMPa- and TgCRMPb-depleted parasites. Infected cells with intact vacuoles were treated with A23187 to induce parasites egress, measured as number of burst vacuoles over the total number of vacuoles. Egress was tested for control (Ctrl), TgCRMPa_iKD and TgCRMPb_iKD lines, in absence of ATc and upon 48 and 24h ATc treatment, respectively. Values are reported as mean ± SD (n = 3 biological replicates, each with 3 technical replicates). *P* values are non-significant for all datasets (two-tailed *t*-test).

(K) Attachment measured for TgCRMPa- and TgCRMPb-depleted parasites. The number of parasites attached to the host cell were counted for control (Ctrl), TgCRMPa_iKD and TgCRMPb_iKD lines, in absence of ATc and upon 48 and 24h ATc treatment, respectively. BAPTA treatment was used as control since it prevents attachment. TgCRMPa- and TgCRMPb-depleted parasites were able to attach to host cells. Values are reported as mean ± SD (n = 3 biological replicates, each with 3 technical replicates). *P* values were measured by two-tailed *t*-test.

Supplementary figure 3

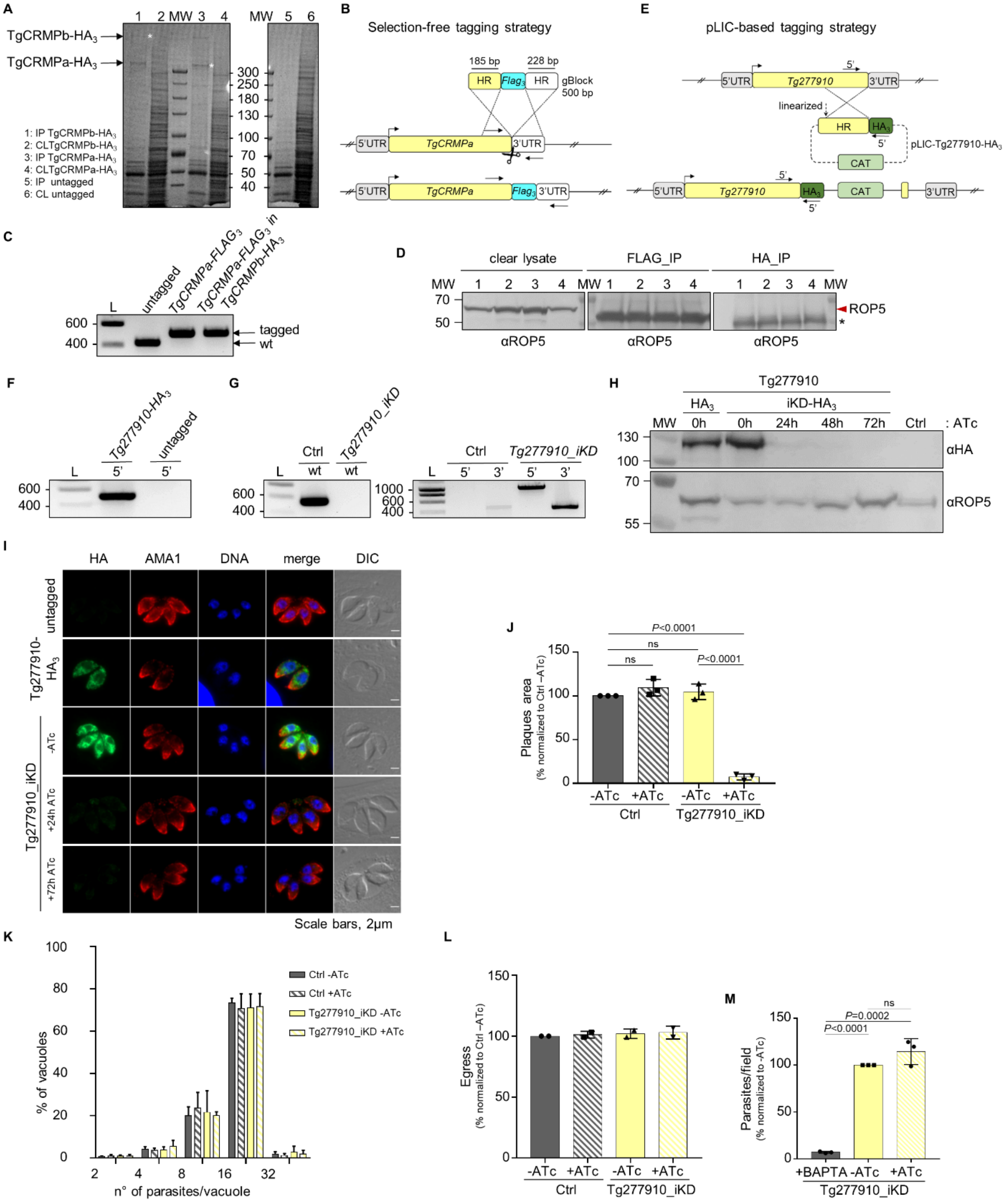

**Figure S3. Tg277910-depleted tachyzoites with a disrupted lytic cycle, show no defects in microneme staining, replication, egress and attachment. Related to figure 3.**

(A) Coomassie Blue staining of eluted proteins (1, 3, 5) immunoprecipitated (IP) with anti-HA beads, and protein fractions of the corresponding clear lysates (CL; 2, 4, 6) prior to beads incubation, from TgCRMPa-$HA_3$, TgCRMPb-$HA_3$ and untagged lines. The TgCRMP protein used as bait in each IP lane is indicated by the asterisk. Samples in lanes 1, 3 and 5 were analyzed by mass spectrometry. MW: molecular weight standards.

(B) Marker-free strategy for $FLAG_3$-tagging of TgCRMPa. To generate a C-terminal $FLAG_3$-fusion of TgCRMPa, a gBlock containing the $FLAG_3$ tag flanked by ~30bp-long TgCRMPa homology regions (HR), was amplified and integrated into the TgCRMPa endogenous locus via CRISPR-cas9 technology (scissors). The $FLAG_3$-tagged TgCRMPa was generated also in the TgCRMPb-$HA_3$ line. The arrows indicate the binding sites of the primers used in (C).

(C) Integration of the $FLAG_3$ tag was tested by PCR in putative TgCRMPa-$FLAG_3$ and TgCRMPa-$FLAG_3$+TgCRMPb-$HA_3$ lines. The addition of the tag at the C-terminus of the TgCRMPa gene corresponds to the insertion of additional 74bp to the wildtype sequence. A higher band was observed in the putative tagged lines compared to the untagged one. DNA ladder (L) is shown on the left of each panel. Primers are listed in table S5.

(D) Eluates from figure 3B and 1/20 of the clear lysates (before beads incubation) were also immunoblotted with anti-ROP5 antibodies, to confirm the specificity of the immunoprecipitation experiments. The red arrowhead indicates TgROP5 protein, the asterisk indicates unspecific bands detected in the eluates, likely corresponding to the light chain of the beads-conjugated antibody. MW: molecular weight standards.

(E) Strategy based on the pLIC system (Huynh and Carruthers, 2009) for tagging Tg277910 with triple HA. The arrows indicate the binding sites of the primers used in (F).

(F) Integration of the $HA_3$ tag and CAT cassette at the C-terminus of TGGT1_277910 was tested by PCR. Genomic DNA from an untagged line and a clonal population for the putative $HA_3$-tagged line, was amplified with primers binding to the 3' C-terminus of TGGT1_277910 and to the $HA_3$ sequence. The $HA_3$ tag (5') was correctly amplified indicating that it was efficiently integrated at the TGGT1_277910 locus. L: DNA ladder (bp). Primers are listed in table S5.

(G) Integration of the DHFR cassette followed by the TetOSag4 promoter in TGGT1_277910 line was tested by PCR as in figures S2D-E. Genomic DNA from untagged parasites, and putative

Tg277910_iKD clonal population, was amplified with primers binding the gene's 5'UTR and the 5' N-terminus, flanking the DHFR-TetOSag4 insert, and used also in pairwise combination with primers binding the DHFR cassette and the Sag4 promoter, respectively. The wildtype fragment (wt) was amplified only in the control line (Ctrl), while the fragments corresponding to the DHFR integration (5') and TetOSag4 integration (3') were detected exclusively in the putative iKD line. A low-abundant unspecific band of similar size to the 3' fragment, was observed in the untagged line. L: DNA ladder (bp). Primers are listed in table S5.

(H) Whole-cell lysates were collected from Tg277910-$HA_3$ parasites ($HA_3$), and from the line generated for the inducible-knockdown (iKD), treated with ATc for 24, 48 and 72h, and untreated. The samples were immunoblotted with anti-HA Abs (upper panel) to visualize Tg277910 protein under all mentioned conditions. TgROP5 was used as loading control (lower panel). A band corresponding to the predicted size for Tg277910 (~138kDa), was detected in the untreated samples (-) and decreased overtime in the ATc-treated ones (+), to completely disappear upon 72h of ATc treatment. Protein molecular weight standards (MW) are shown on the left of each panel.

(I) Immunofluorescence images of untagged, Tg277910-$HA_3$ and Tg277910-depleted (iKD) intracellular tachyzoites. Parasites were stained with anti-HA and anti-AMA1 Abs to label Tg277910 and micronemes, respectively. The nuclei (DNA) are stained with Hoechst. Tg277910-$HA_3$ pattern mirrors that of figure 3D. Micronemes show a wildtype appearance in the Tg277910-depleted parasites. Shown are single focal planes.

(J) Quantification of plaques for Tg277910-depleted parasites. Lysis plaque areas were measured for untreated and 72h ATc-treated control and iKD lines. Values are reported as mean ± SD (n = 3 biological replicates, each with 3 technical replicates).

(K) Quantification of replication, (L) egress and (M) attachment for control (Ctrl) and Tg277910-depleted (iKD) lines were performed as in figures S2H, S2I, S2J, respectively, with 72h ATc-treated and untreated parasites. Tg277910-depleted parasites replicate, egress and attach normally. Values are reported as mean ± SD (n = 3 biological replicates, each with 3 technical replicates). *P* values were measured by two-tailed *t*-test and were non-significant specifically for "replication" and "egress" datasets.

Supplementary figure 4

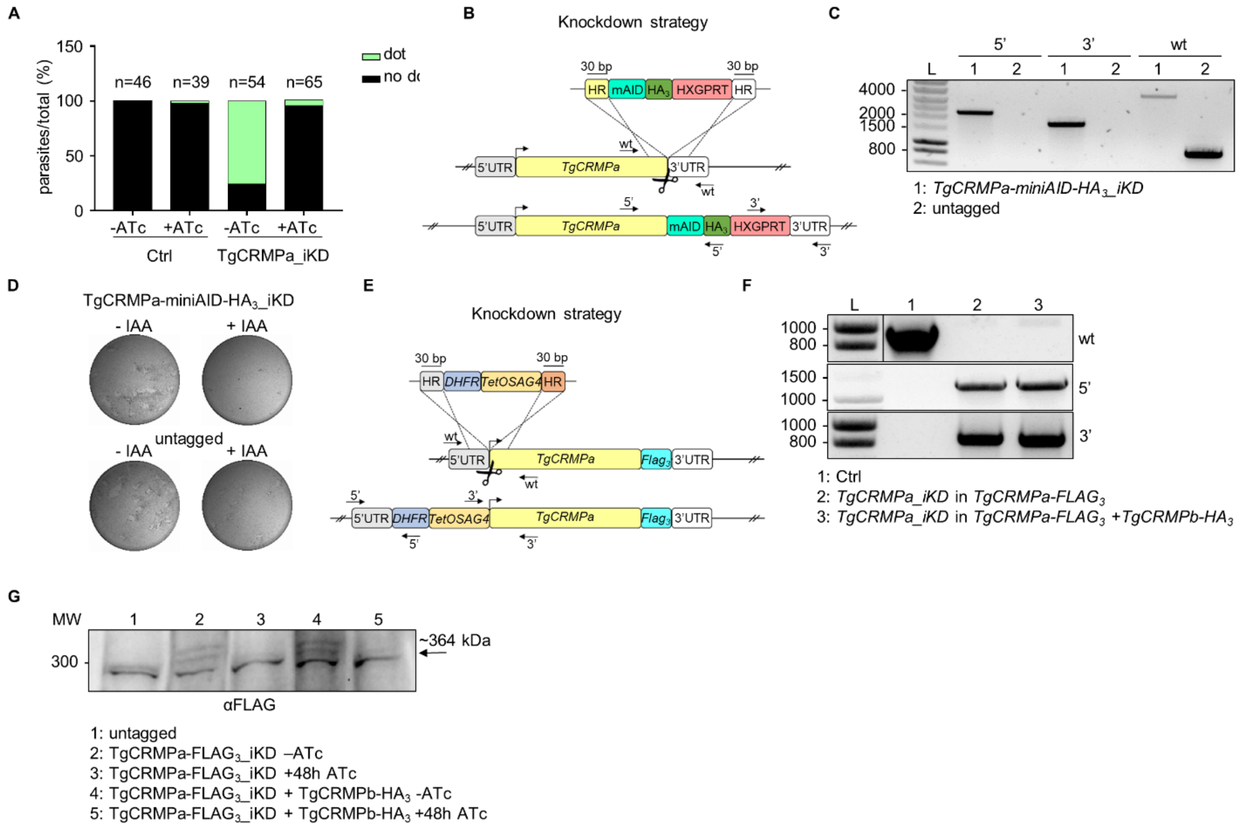


**Figure S4. TgCRMPa and TgCRMPb accumulate at the tip of the extruded conoid. Related to figure 5.**

(A) Quantification of the dot pattern for TgCRMPa-$HA_3$ in TgCRMPa-depleted (iKD) tachyzoites. TgCRMPa accumulation at the apical tip of extracellular parasites was measured upon incubation with host cell monolayers for 2min, to stimulate natural conoid extrusion. CRMPa signal at the apical dot disappeared after 48h ATc treatment, indicating that the association with the tip of the extruded conoid was specific. No significant apical signal was detected for the control line (Ctrl), as in figure 5B. Numbers are expressed as percentage of parasites showing (dot) or lacking (no dot) the tip accumulation of TgCRMPa. The number of parasites (n) analyzed for each line is reported on the column tops.

(B) Auxin-degron strategy used for generating TgCRMPa-miniAID-$HA_3$ strain. The integration of the tag and drug resistance cassette into the TgCRMPa locus is ensured by ~30bp-long homology regions (HR) upon CRISPR-Cas9 activity (scissors). The arrows indicate the binding sites of the primers used in (C).

(C) Integration of the miniAID-$HA_3$ and HXGPRT cassette at the *TgCRMPa* locus in the Tir-1 line was tested by PCR as in figure S2D. The fragments corresponding to the miniAID-$HA_3$ (5') and HXGPRT cassette (3') integration were detected exclusively in the putative iKD line, while the wildtype fragment (wt) was amplified only in the untagged line. A ~4000bp fragment corresponding to the miniAID-$HA_3$+HXGPRT cassette and amplified with primers binding the wildtype sequence, was detected in the iKD line. L: DNA ladder (bp). Primers are listed in table S5.

(D) Representative images of lytic plaques formation in HFF monolayers infected with IAA-treated and untreated Tir-1 control and TgCRMPa-miniAID-$HA_3$_iKD lines.

(E) Strategy for the inducible depletion (iKD) of TgCRMPa-$FLAG_3$. The iKD lines were generated starting from the $FLAG_3$-tagged lines previously produced. In order to conditionally deplete the protein, the endogenous promoter of the TgCRMPa-$FLAG_3$ gene was replaced with an ATc-regulableTetOSag4 promoter, preceded by the DHFR resistance cassette. The DNA fragment containing the cassette and the promoter was PCR-amplified from a donor vector with primers containing ~30bp-long homology regions (HR) specific for *TgCRMPa* gene, and introduced upstream the starting codon via CRISPR-cas9 technology (scissors) and homologous recombination. The arrows indicate the binding sites of the primers used in (G).

(F) Integration of the DHFR cassette followed by the TetOSag4 promoter in the putative TgCRMPa-$FLAG_3$ and TgCRMPa-$FLAG_3$+TgCRMPb-$FLAG_3$ iKD lines was tested by PCR as in figure S2D. The fragments corresponding to the DHFR integration (5') and TetOSag4 integration (3') were detected exclusively in the putative iKD lines, while the wildtype fragment (wt) was amplified only in the control line (Ctrl). L: DNA ladder (bp). Primers are listed in table S5.

(G) Whole-cell lysates from untagged, TgCRMPa-$FLAG_3$_iKD and TgCRMPa-$FLAG_3$_iKD+TgCRMPb-$HA_3$ lines, were immunoblotted with anti-FLAG Abs to visualize tagged CRMPa in ATc-treated and untreated samples. CRMPa disappeared upon 48h ATc incubation in both lines. A ~300kDa unspecific cross-reactive band was observed in all samples. MW: molecular weight standards.

Supplementary figure 5

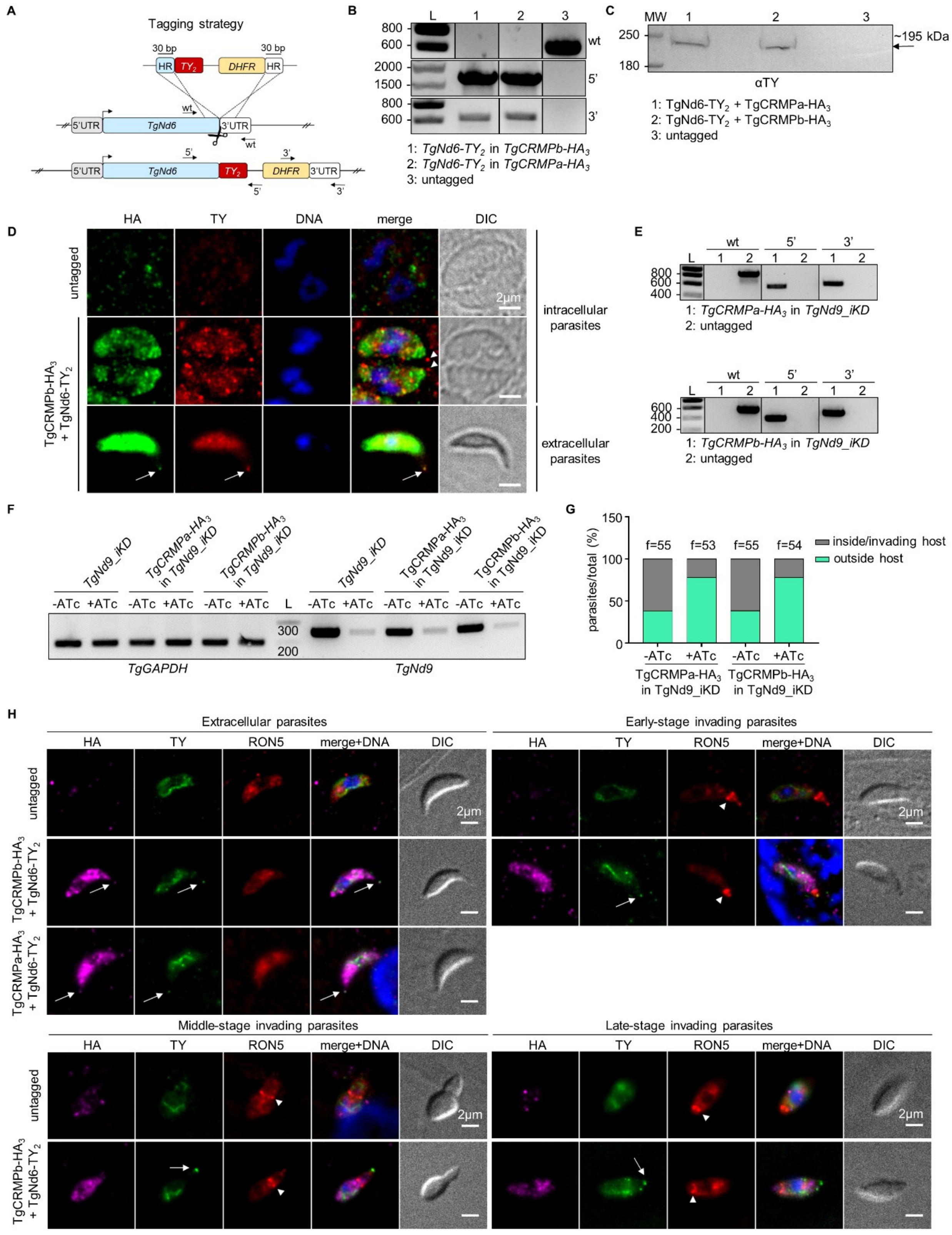

**Figure S5. CRMP and Nd complexes show different dynamics at the exocytic site in *Toxoplasma gondii*. Related to figure 6.**

(A) Strategy for $TY_2$-tagging of TgNd6 in TgCRMPa-$HA_3$ and TgCRMPb-$HA_3$ lines. To generate C-terminal $TY_2$-fusion of TgNd6, a DNA fragment was amplified from a donor vector containing the $TY_2$ tag and the drug resistance cassette (DHFR). Primers to amplify the DNA fragment were designed to contain 30bp-long stretches (HR) homologous to TgND6 regions flanking the insertion site for the epitope tag. Upon CRISPR-cas9 cut (scissors), the PCR-amplified DNA fragment efficiently recombines into the targeted endogenous locus. The arrows indicate the binding sites of the primers used in (B).

(B) Integration of the $TY_2$ tag and DHFR cassette at the C-terminus of TgNd6 was tested by PCR. Genomic DNAs from an untagged line and clonal populations for TgNd6-$TY_2$+TgCRMPa-$HA_3$ and TgNd6-$TY_2$+TgCRMPb-$HA_3$ lines, were amplified with primers binding to the 3' C-terminus and 3'UTR of *TgNd6*, and also in pairwise combination with primers binding the $TY_2$ and DHFR sequences, respectively. The fragments corresponding to the $TY_2$ tag (5') and the resistance cassette (3') were correctly amplified in the putative tagged lines, indicating that they were efficiently integrated at the *TgNd6* locus. As expected, the wildtype fragment for *TgNd6* (wt) was detected only in the untagged line. L: DNA ladder (bp). Primers are listed in table S5.

(C) Whole-cell lysates from untagged, TgNd6-$TY_2$+TgCRMPa-$HA_3$ and TgNd6-$TY_2$+TgCRMPb-$HA_3$ parasites, were immunoblotted with anti-TY Abs to detect tagged Nd6. A band around the expected size (~195kDa) for TgNd6-$TY_2$ was observed exclusively for the tagged lines. MW: molecular weight standards.

(D) Immunofluorescence images of intracellular (upper and middle panels) and extracellular (lower panel) tachyzoites from untagged and TgCRMPb-$HA_3$+TgNd6-$TY_2$ lines. Extracellular parasites were incubated with host cell monolayers for 2min prior to fixation. Parasites were stained with anti-HA and anti-TY Abs to label CRMPb and Nd6, respectively. Nd6, but not CRMPb, accumulates at the tachyzoite apex in intracellular parasites (arrowheads), while both proteins localize at the tip of the extruded conoid in extracellular parasites (arrows). DNA is labeled by Hoechst. Single focal planes are shown. DIC, differential interference contrast.

(E) Integration of the $HA_3$ tag and CAT cassette at the C-terminus of *TgCRMPa* and *TgCRMPb* genes in TgNd9_iKD line was tested by PCR as in figures S2B-C. The fragments corresponding to the $HA_3$ tag (5') and the resistance cassette (3') were correctly amplified in the putative tagged lines, indicating that they were efficiently integrated at the TgCRMPs loci. As expected, the wildtype fragment of each gene (wt) was detected only in the untagged line. L: DNA ladder. Primers are listed in table S5.

(F) Depletion of *TgNd9* transcripts was assessed by RT-PCR for the experiment shown in figure 6D. Total RNAs from TgCRMPa-$HA_3$ and TgCRMPb-$HA_3$ expressed in TgNd9_iKD (minus epitope tag) parasites and parental line were subjected to reverse transcription and PCR-amplified with primers binding *TgNd9* transcripts. *TgGAPDH* was used as housekeeping gene. *TgNd9* transcripts strongly decreased upon 72h ATc treatment (+ATc). L: DNA ladder (L). Primers are listed in table S5.

(G) Depletion of TgNd9 proteins in the lines used for the experiment in figure 6D was also assessed by quantifying the defect in invasion of ATc-treated TgNd9_iKD parasites expressing TgCRMPa-$HA_3$ and TgCRMPb-$HA_3$, versus untreated. The values are reported as percentages of the number of invading/intracellular and extracellular parasites, over the total number of parasites. The number of fields (f) analyzed for each line is reported on the column tops.

(H) Immunofluorescence images of extracellular parasites, and of parasites in early, middle and late stages of host cell invasion. Parasites co-expressing TgCRMPa-$HA_3$ and TgCRMPb-$HA_3$ with TgNd6-$TY_2$ were incubated with host cell monolayers and stained as in figure 6D. Untagged parasites were treated in parallel. In contrast with TgNd6 (arrow), the apical accumulation of TgCRMPa and TgCRMPb observed in extracellular parasites, disappears upon entering the host and remains undetected for the entire process. TgCRMPa data for invading parasites are shown in figure 6D. The moving junction is indicated by the arrowhead. Non-specific anti-TY labeling of mitochondria was detected for both untagged and tagged lines. DIC: differential interference contrast. Single focal planes are shown.
