## Supplementary material for "An apical membrane complex controls rhoptry exocytosis and invasion in *Toxoplasma*": Table S4

### Ferlins retrieved by BLAST and reciprocal BLAST for phylogenetic tree construction

| Gene ID | Species | BLAST E-value (TgFer2 query) | RECIPROCAL BLAST E-value (TgFer2 query top hit) | RECIPROCAL BLAST E-value (TgFer1 second hit) | RECIPROCAL BLAST E-value (TgFer1 top hit) | RECIPROCAL BLAST E-value (TgFer2 query second hit) |
|---|---|---|---|---|---|---|
| **TGME49 260470 (TgFer2; query)** | Toxoplasma gondii strain ME49 | 0.00E+00 | n.a. | n.a. | n.a. | n.a. |
| TGME49 309420 (TgFer1) | Toxoplasma gondii strain ME49 | 2.00E-24 | n.a. | n.a. | 0.00E+00 | 2.00E-24 |
| PF3D7 0806300 | Plasmodium falciparum strain 3D7 | 1.00E-23 | n.a. | n.a. | 0.00E+00 | 2.00E-25 |
| PF3D7 1455600 | Plasmodium falciparum strain 3D7 | 5.00E-128 | 2.00E-131 | 2.00E-20 | n.a. | n.a. |
| CPATCC 0028490 | Criptosporidium parvum strain IOWA-ATCC | 0.00E+00 | 0.00E+00 | 3.00E-25 | n.a. | n.a. |
| CPATCC 0003050 | Criptosporidium parvum strain IOWA-ATCC | 2.00E-25 | n.a. | n.a. | 0.00E+00 | 1.00E-25 |
| CPATCC 0021940 | Criptosporidium parvum strain IOWA-ATCC | 1.00E-21 | n.a. | n.a. | 2.00E-62 | 3.00E-21 |
| TTHERM 00440600 | Tetrahymena thermophila | 1.00E-35 | 9.00E-37 | 3.00E-29 | n.a. | n.a. |
| TTHERM 00440610 | Tetrahymena thermophila | 2.00E-13 | n.a. | n.a. | 4.00E-24 | 6.00E-20 |
| TTHERM 00823910 | Tetrahymena thermophila | 2.00E-26 | 7.00E-27 | 3.00E-25 | n.a. | n.a. |
| TTHERM 00886960 | Tetrahymena thermophila | 7.00E-30 | n.a. | n.a. | 7.00E-40 | 2.00E-31 |
| IMG5 007260 | Ichthyophthirius multifiliis | 1.00E-26 | n.a. | n.a. | 1.00E-38 | 4.00E-28 |
| IMG5 161110 | Ichthyophthirius multifiliis | 2.00E-24 | n.a. | n.a. | 6.00E-31 | 2.00E-25 |
| IMG5_151880 | Ichthyophthirius multifiliis | 3.00E-23 | n.a. | n.a. | 8.00E-24 | 2.00E-23 |
| GSPATP00029587001 | Paramecium tetraurelia strain d4-2 protein (v1.0) | 2.00E-30 | n.a. | n.a. | 1.00E-33 | 8.00E-31 |
| GSPATP00038277001 | Paramecium tetraurelia strain d4-2 protein (v1.0) | 2.00E-31 | n.a. | n.a. | 4.00E-40 | 7.00E-32 |
| GSPATP00039640001 | Paramecium tetraurelia strain d4-2 protein (v1.0) | 6.00E-28 | 6.00E-28 | 6.00E-28 | 6.00E-28 | 6.00E-28 |
| GSPATP00037942001 | Paramecium tetraurelia strain d4-2 protein (v1.0) | 3.00E-20 | n.a. | n.a. | 1.00E-22 | 1.00E-20 |
| TGME49 295472 (TgFer3; outgroup) | Toxoplasma gondii strain ME49 | | | | | |

### CRMPs retrieved by BLAST and reciprocal BLAST for phylogenetic tree construction

| Gene ID | Species | BLAST E-value (TgCRMPa query) | BLAST E-value (TgCRMPb query) | RECIPROCAL BLAST E-value (TgCRMPa query top hit) | RECIPROCAL BLAST E-value (TgCRMPb second hit) | RECIPROCAL BLAST E-value (TgCRMPa query second hit) | RECIPROCAL BLAST E-value (TgCRMPb fir |
|---|---|---|---|---|---|---|---|
| **TGME49 261080 (CRMPa query)** | *Toxoplasma gondii* | **0.00E+00** | 5.00E-35 | n.a. | n.a. | n.a. | n.a. |
| **TGME49 292020 (CRMPb query)** | *Toxoplasma gondii* | 1.00E-30 | **0.00E+00** | n.a. | n.a. | n.a. | n.a. |
| PF3D7 0718300 | *Plasmodium falciparum* | 0.00E+00 | 5.00E-34 | 0.00E+00 | 3.00E-35 | n.a. | n.a. |
| PF3D7 1208200 | *Plasmodium falciparum* | 2.00E-22 | 3.00E-70 | n.a. | n.a. | 4.00E-20 | 1.00E-78 |
| PF3D7 1475400 | *Plasmodium falciparum* | 6.00E-16 | 7.00E-41 | n.a. | n.a. | 9.00E-21 | 8.00E-42 |
| PF3D7 0911300 | *Plasmodium falciparum* | 4.00E-137 | 6.00E-24 | 2.00E-149 | 3.00E-25 | n.a. | n.a. |
| PBANKA 0615900 | *Plasmodium berghei* | 0.00E+00 | 7.00E-33 | 0.00E+00 | 2.00E-28 | n.a. | n.a. |
| PBANKA 0812400 | *Plasmodium berghei* | 2.00E-136 | 1.00E-24 | 2.00E-143 | 8.00E-26 | n.a. | n.a. |
| PBANKA 0606700 | *Plasmodium berghei* | 7.00E-28 | 2.00E-69 | n.a. | n.a. | 8.00E-26 | 4.00E-75 |
| PBANKA 1300800 | *Plasmodium berghei* | 7.00E-23 | 4.00E-29 | n.a. | n.a. | 2.00E-24 | 5.00E-27 |
| NCLIV 026200 | *Neospora caninum* | 0.00E+00 | 3.00E-34 | 0.00E+00 | 1.00E-30 | n.a. | n.a. |
| NCLIV 012890 | *Neospora caninum* | 3.00E-29 | 0.00E+00 | n.a. | n.a. | 1.00E-34 | 0.00E+00 |
| EfaB MINUS 23210.g2033 | *Eimeria falciformis* | 0.00E+00 | 1.00E-33 | 0.00E+00 | 3.00E-32 | n.a. | n.a. |
| EfaB MINUS 49504.g2828 | *Eimeria falciformis* | 0.00E+00 | 8.00E-29 | 0.00E+00 | 5.00E-30 | n.a. | n.a. |
| EfaB PLUS 52892.g2741 | *Eimeria falciformis* | 4.00E-71 | 0.00E+00 | n.a. | n.a. | 8.00E-71 | 0.00E+00 |
| BEWA 042570 | *Theileria equi* | 0.00E+00 | 2.00E-31 | 0.00E+00 | 1.00E-29 | n.a. | n.a. |
| BEWA 042580 | *Theileria equi* | 0.00E+00 | 3.00E-29 | 0.00E+00 | 5.00E-32 | n.a. | n.a. |
| BEWA 029890 | *Theileria equi* | 8.00E-31 | 1.00E-115 | n.a. | n.a. | 4.00E-38 | 7.00E-108 |
| BEWA 029850 | *Theileria equi* | 3.00E-27 | 2.00E-75 | n.a. | n.a. | 3.00E-38 | 2.00E-77 |
| BBBOND 0404220 | *Babesia bigemina* | 0.00E+00 | 3.00E-25 | 0.00E+00 | 7.00E-29 | n.a. | n.a. |
| BBBOND 0404210 | *Babesia bigemina* | 0.00E+00 | 9.00E-33 | 0.00E+00 | 9.00E-35 | n.a. | n.a. |
| BBBOND 0208300 | *Babesia bigemina* | 8.00E-33 | 4.00E-78 | n.a. | n.a. | 2.00E-28 | 5.00E-79 |
| BBBOND 0208320 | *Babesia bigemina* | 8.00E-17 | 7.00E-71 | n.a. | n.a. | 8.00E-17 | 1.00E-127 |
