## Supplementary material for "An apical membrane complex controls rhoptry exocytosis and invasion in *Toxoplasma*": Table S5

### Primers *Toxoplasma gondii*

#### guideRNAs

| | | |
|---|---|---|
| | **TgCRMPA** | |
| **gRNA at the C-terminus** | | |
| ML3283 | aagttGTGAAGACGCTGTCTTTGCACg | F_gRNA1_261080 integration HA3-CAT, mAID, FLAG3 C-terminus |
| ML3284 | aaaacGTGCAAAGACAGCGTCTTCACa | R_gRNA1_261080 integration HA3-CAT, mAID, FLAG3 C-terminus |
| **gRNA at the N-terminus** | | |
| ML3342 | aagttGCTTTCGACTGCAGCAAGCGGg | F_gRNA3_261080 integration tetOFF |
| ML3343 | aaaacCCGCTTGCTGCAGTCGAAAGCa | R_gRNA3_261080 integration tetOFF |
| | **TgCRMPB** | |
| **gRNA at the C-terminus** | | |
| ML3279 | aagttCCATGTGAAGGCCGATCAGGAg | F_gRNA1_292020 Integration HA3-CAT |
| ML3280 | aaaacTCCTGATCGGCCTTCACATGGa | R_gRNA1_292020 Integration HA3-CAT |
| **gRNA at the N-Terminus** | | |
| ML3338 | aagttGAAACAGGGAGAGTGGGCACg | F_gRNA3_292020 integration TetOff |
| ML3339 | aaaacGTGCCCACTCTCCCTGTTTCa | R_gRNA3_292020 integration TetOff |
| | **Tg277910** | |
| **gRNA at the N-Terminus** | | |
| ML3970 | aagttGTCACACCTGAGAAAAGTGAGg | F_gRNA1 277910 TetOff |
| ML3971 | aaaacCTCACTTTTCTCAGGTGTGACa | R_gRNA1 277910 TetOff |
| | **Toxofilin Strains** | |
| ML2087 | AAGTTGGCTCCCACGTCCCTCACCATG | 3' guide UPRT Toxofilin Frwd |
| ML2088 | AAAACATGGTGAGGGACGTGGGAGCCA | 3' guide UPRT Toxofilin Rev |
| ML3445 | AAGTTGCAGGGCTTCTAAAA GGCGCG | 5' guide UPRT Toxofilin Frwd |
| ML3446 | AAAACGCGCCATTTTAGAAGCCCTGCA | 5' guide UPRT Toxofilin Rev |
| | **TgND6** | |
| **gRNA at the C-terminus** | | |
| ML3129 | AAGTTGTTTTATGCGCTCTACTGTGGG | F_248640_TY_DHFR C-terminus (Tgnd6CtgRNA_Fw) |
| ML3130 | AAAACCCACAGTAGAGCGCATAAAACA | R_248640_TY_DHFR C-terminus (Tgnd6CtgRNA_Rev) |

#### Integration_specific to GOI

| | | |
|---|---|---|
| | **TgCRMPA** | |
| ML3348 | GACACTGCGTAAGCTCGAA | F_int_HA_261080 |
| ML3349 | CTCCTATCGTACAAGCTGTGAAC | R_int_HA_261080 |
| ML3388 | TACAGACGGCCACTGCTTC | F_intTeTOSAG4_261080 |
| ML3389 | CGAAGACACTTCGACTGCAAGTGCT | R_intTeTOSAG4_261080 |
| ML3002 | CGTCTCTCACTTGTGTCATGT | gBlockCRMPA-flag_fw C-terminus |
| ML3003 | AAAGAACAAAGAAGTGCTTG | gBlockCRMPA-flag_rev C-terminus |
| | **TgCRMPB** | |
| ML3346 | GACGTGAAGAATCTGGATCGGA | F_int HA_292020 |
| ML3347 | CCTGCTCACGTTAGCGATAGT | R_int_HA 292020 |
| ML3386 | GCCAATGCGACATAAATCCACACA | F_intTeTOSAG4_292020 |
| ML3387 | TCGATCGACTTCGCGTCTCA | R_intTeTOSAG4_292020 |
| | **Tg277910** | |
| ML3968 | GACGCGCGCATGCACTGATG | Frwd_int_277910 TetOff |
| ML3969 | GCAGAGGATGAAGCAGAGCGCG | Rev_int_277910 TetOff |
| ML3972 | GAAGCGCTCAGAGACTGTG | Frwd int HA3-CAT 277910 |
| | **Toxofilin Strains** | |
| ML3187 | CTCGGCTCCATCTCATTCC | Toxofilin in UPRT Frwd |
| ML3546 | GAGGGCTAGCAAAGCGTCA | Toxofilin in UPRT Rev |
| ML3547 | GTATCAGTTGTGCGCGGAAG | Toxofilin in UPRT Frwd |
| ML3190 | TCCCGTTACAGGTGTACGGG | Toxofilin in UPRT Rev |
| | **TgND6** | |
| ML3006 | CGTTCAGAAGGTCGAGCAAC | C-terminal TY-DHFR fw (Tgnd6_checkFW) |
| ML3007 | CGTCTGCAGCCTTATTCCAC | C-terminal TY-DHFR rev (Tgnd6_3'checkRev) |

#### Integration_specific to DNA cassette

| | | |
|---|---|---|
| ML1476 | CAGCGTAGTCCGGGACGTCGTAC | HA rev |
| ML388 | ACAGTACTGCGATGAGTGGC | CAT fw |
| ML2456 | ACGCGTCGCAGCATACT | TetOff TeTOSAG4promoter fw |
| ML2454 | ACGGGAGGCCGTTGTTGAT | TetOff DHFR rev |
| ML687 | GTTTGAATGCAAGGTTTCGTGCTGTCG | TetOff_DHFR_rev |
| ML2873 | GGATCTGCACACCTGGTCTCGATG | TetOff_DHFR_rev & DHFR+TY_rev |
| ML1041 | CGGATCATTTGAAAACATCGTGAGGCTGG | TetOff_TeTOSAG4promoter_fw |
| ML4875 | ATGTGGACACAGTCGGTTGA | DHFR+TY fw |
| ML4136 | CGACAACACCTTCTACAACGC | mAID HA3 FW |
| ML4137 | TCAGGCGGACATAGTGCTC | mAID HA3 REV |

#### Cassette amplification

| | | |
|---|---|---|
| | **TgCRMPA** | |
| ML3287 | AGGGAACTACGAGGCGGCTCCCGACGACGGTCTgTACCCGTACGACGTCC | F_281080_HA_CAT |
| ML3288 | TGCTTGAAAAAAGCGCTATTCCTTGTTGTGTCTAGAACTAGTGGATCCCC | R_281080_HA_CAT |
| ML4909 | ACTACGAGGCGGCTCCCGACGACGGTCTGCCTAGGATGGTGAGCGCTAGC | Frwd mAID HA3 C-terminus |
| ML4910 | TGACAAGTGTTCTGGCAGGCTTTGCTCGAACTTTAAAGCTACGCGTTGTA | Rev mAID HA3 C-terminus |
| ML3317 | TCTGCAGCCGCTGTGGACGCTTGTCTGGACTCAAGCTTCGCCAGGCTGTA | F_TetOff_261080 |
| ML3318 | AACGCGGAGATGTTCCGTTGCGCTCTGCATAGATCTGGTTGAAGACAGAC | R_TetOff_261080 |
| | **TgCRMPB** | |
| ML3315 | CCCCGCGGACACCGATGCCAGGGCAAGCGACGAAGCTTCGCCAGGCTGTA | F_TetOff_292020 |
| ML3316 | GTTGTTCGCATCCGACGCCAGAGAACTCATAGATCTGGTTGAAGACAGAC | R_TetOff_292020 |
| ML3277 | ACAATTGAAGAAGCCCGGGTCTCCGAAACAGGAgTACCCGTACGACGTCC | F_292020_HA_CAT |
| ML3278 | ACATTTCGCCGGGGTCTTTTTTATTCTCGGTCTAGAACTAGTGGATCCCC | R_292020_HA_CAT |
| | **Tg277910** | |
| ML3966 | CCATTCTCCGAGCTTCGTCTCGCAGCTGTCGATAAGCTTCTTCGCCAGGCTG | TetOff DHFR 277910_fw |
| ML3967 | GAGAGAGGCGCTTCTGCGAGTCGCCGGCATCATAGATCTGGTTGAAGACAGACG | TetOff DHFR 277910_rev |
| ML4046 | TACTTCCAATCCAATTTAATGCATGCGTTTCCGTGGAT | HA-CAT_fw for pLIC |
| ML4047 | TCCTCCACTTCCAATTTTAGCCATGGACCCAAAGGATTCGG | HA-CAT_rev for pLIC |
| | **Toxofilin in UPRT locus** | |
| ML3522 | TTGGTCAGCCTCCACAAGGCGTATTCCTCAAAGCACGGAGGAGAGACG | Toxofilin Cassette_fw + Homology for UPRT |
| ML3523 | CTTGTCGCGCACCAGGCAGCCTCACAACTATAGGACCCCAGGCTTTACA | Toxofilin Cassette_rev + Homology for UPRT |
| | **TgND6** | |
| ML4734 | CAAGCGCTCGACCAAGTGGGGGGGCAGGAGGCCGCTAGCAAGGGCTCGGG | F_248640_TY_DHFR C-terminus |
| ML4735 | TCCCGACGAAAGGACACGGCGCGAACTCACATACGACTCACTATAGGG | R_248640_TY_DHFR C-terminus |

#### RT-PCR

| | | |
|---|---|---|
| | **TgND9** | |
| ML4741 | GTGGAGCATGCGCATTAGTCAG | F_RT_249730 |
| ML4742 | GGTCGTTGAAGACATAGTCGCG | R_RT_249730 |
| | **TgGAPDH** | |
| ML3617 | TGTGCAAGCTGGGTATTAAC | Fw_RT_GAPDH |
| ML3618 | CCGACAATGAGCTTGCC | Rv_RT_GAPDH |

### Primers *Tetrahymena thermophila*

#### Knockout (KO) construct

| | | |
|---|---|---|
| | **TtFER2** | |
| ML4379 | aaagagctctacaatgtgtgaattgatcaaggaaag | 5'UTR_00886960_sacI_fw |
| ML4380 | aaactgcagtgataaagaatattcaaaatttatgttgc | 5'UTR_00886960_pstI_rev |
| ML4381 | attaagcttgataattaattgtattaaacgtgtaac | 3'UTR_00886960_hindIII_fw |

| | | |
|---|---|---|
| ML4382 | aaactcgagttacaagtttttatattctagaggtc | 3'UTR_00886960_xhoI_rev |
| | **TtCRMP1** | |
| ML3830 | GCATgagctcccattttattgaaatactaatgcaaaatttactg | 5'UTR_sacI_00442310 |
| ML3831 | tcactgcagcatcacagaattattattttaaaacgaattaac | 5'UTR_pstI_00442310 |
| ML3832 | GCATaagcttgtgaaacctgtctaaactgggaaccta | 3'UTR_HindIII_00442310 |
| ML3833 | TCAGctcgaggcatgaatgactgattatcttgatagag | 3'UTR_XhoI_00442310 |
| | **TtCRMP2** | |
| ML4283 | aaaGAGCTCtgatcagcttagattaaatttctgt | 5'utr00637180_sacI_fw |
| ML4284 | aaaCTGCAGgttccaatttaaaaaatctaataagatc | 5'utr00637180_pstI_rev |
| ML4285 | attAAGCTTtacttacttacgcagagatttccact | 3'utr00637180_hindIII_fw |
| ML4286 | aaaCTCGAGgatatattaaatgatgaaacaaccagaaagt | 3'utr00637180_XhoI_rev |
| | **RT-PCR to assess complete KO** | |
| | **TtFER2** | |
| ML4518 | attagaacagtgatgataaaaggcgt | RT_00886960_fw2 |
| ML4519 | taactttcttgctgtagatgaagtag | RT_00886960_rev2 |
| | **TtCRMP1** | |
| ML4510 | gattagcttaaatacgtaactcatgct | RT_00442310_fw2 |
| ML4511 | taacaaatcttgattagtagctgtcat | RT_00442310_rev2 |
| | **TtCRMP2** | |
| ML4508 | tgtagtgcttgtaattattatccaact | RT_00637180_fw2 |
| ML4509 | atcttcacactactttaaacaacttg | RT_00637180_rev2 |
| | **BTU1 (TTHERM_00348510)** | |
| ML3890 | ATGAGAGAAATCGTTCACATC | RT_BTU1_fw |
| ML3889 | TGACCGAAAACGAAGTTATC | RT_BTU1_rev |
